# Species-Dependent Heterophilic and Homophilic Cadherin Interactions in Intestinal Intermicrovillar Links

**DOI:** 10.1101/2020.09.01.278846

**Authors:** Michelle E. Gray, Zachary R. Johnson, Debadrita Modak, Matthew J. Tyska, Marcos Sotomayor

**Author notes:** Co-first authors (Michelle E. Gray, Zachary R. Johnson, Debadrita Modak). Marshall University, Joan C. Edwards School of Medicine, Huntington, WV 25701.

## Abstract

Enterocytes are specialized epithelial cells lining the luminal surface of the small intestine that build densely packed arrays of microvilli known as brush borders. These microvilli drive nutrient absorption and are arranged in a hexagonal pattern maintained by intermicrovillar links formed by two non-classical members of the cadherin superfamily of calcium-dependent cell adhesion proteins: protocadherin-24 (PCDH24, also known as CDHR2) and the mucin-like protocadherin (CDHR5). The extracellular domains of these proteins are involved in heterophilic and homophilic interactions important for intermicrovillar function, yet the structural determinants of these interactions remain unresolved. Here we present X-ray crystal structures of the PCDH24 and CDHR5 extracellular tips and analyze their species-specific features relevant for adhesive interactions. In parallel, we use binding assays to identify the PCDH24 and CDHR5 domains involved in both heterophilic and homophilic adhesion for human and mouse proteins. Our results suggest that homophilic and heterophilic interactions involving PCDH24 and CDHR5 are species dependent with unique and distinct minimal adhesive units.

## INTRODUCTION

Enterocytes are specialized epithelial cells lining the luminal surface of the small intestine and are fundamental players in nutrient absorption with a role in host defense (Maroux et al., 1988; Mukherjee et al., 2008; Selsted and Ouellette, 2005; Shifrin and Tyska, 2012). Key to the aforementioned processes is the brush border, so named because the apical surface of the enterocytes is coated by thousands of microvilli of similar length and size organized in a hexagonal arrangement (Mooseker, 1985). The organization and structure of the microvilli is maintained by a network of cytoplasmic and transmembrane proteins known as the intermicrovillar adhesion complex (IMAC), which includes proteins with extracellular adhesive domains and cytoplasmic parts tethered to the actin cytoskeleton that forms the interior of the microvilli (Crawley et al., 2016, 2014b, 2014a; Li et al., 2016; Pinette et al., 2018; Weck et al., 2016). Perturbations of brush border function are often associated with disease (Bailey et al., 1989; Khubchandani et al., 2011; Vallance et al., 2002; Wilson et al., 2001) and, not surprisingly, disruption of the IMAC components results in intestinal dysfunction (Bitner-Glindzicz et al., 2000; Crawley et al., 2014b; Hussain et al., 2004).

The extracellular parts of the IMAC stem from a pair of cellular adhesion proteins that belong to the cadherin superfamily, protocadherin-24 (PCDH24, also known as CDHR2) and mucin-like protocadherin (CDHR5) (Crawley et al., 2014b; McConnell et al., 2011). These two non-classical protocadherins form intermicrovillar links essential for brush border morphogenesis and function (Crawley et al., 2014a, 2014b), stabilizing the microvillar hexagonal patterns as shown by freeze-etch electron microscopy and PCDH24 immuno-labeling combined with transmission electron microscopy (Crawley et al., 2014b). Mutations that impair the PCDH24 and CDHR5 interaction, along with knockdowns of PCDH24 and CDHR5, cause a remarkable reduction in microvillar clustering *ex vivo* (Crawley et al., 2014b). A PCDH24 knockout mouse model was found to be viable but body weight of mutant mice was lower than wild-type and there were defects in the packing of the microvilli in the brush border (Pinette et al., 2018). Despite the important physiological role played by PCDH24 and CDHR5, little is known about the molecular details of how these proteins interact to form the intermicrovillar links.

PCDH24 and CDHR5 are members of a large superfamily of proteins with extracellular domains that often mediate adhesion and that have contiguous and similar, but not identical, extracellular cadherin (EC) repeats. The linker regions between the EC repeats typically bind three calcium ions essential for adhesive function (Frank and Kemler, 2002; Gumbiner, 2005; Sano et al., 1993; Takeichi, 1990). PCDH24 belongs to the Cr-2 subfamily and has 9 EC repeats, a membrane adjacent domain (MAD10), a transmembrane domain, and a C-terminal cytoplasmic domain (Bork and Patthy, 1995; De-la-Torre et al., 2018; Ge et al., 2018; Hulpiau and van Roy, 2009; McConnell et al., 2011; Pei and Grishin, 2017). CDHR5 belongs to the Cr-3 subfamily and has 4 EC repeats, a mucin-like domain (MLD), a transmembrane domain, and a C-terminal cytoplasmic domain. Some human CDHR5 isoforms lack the MLD, which is not essential for the interaction with human PCDH24 (Crawley et al., 2014b). Sequence alignments of the N-terminal repeats (EC1-3) suggest that PCDH24 and CDHR5 are similar to another pair of heterophilic interacting cadherins: cadherin-23 (CDH23) and protocadherin-15 (PCDH15), the cadherins responsible for the formation of inner-ear tip links (Elledge et al., 2010; Kazmierczak et al., 2007; Sotomayor et al., 2014, 2012, 2010). PCDH24 is most similar to CDH23, having the residues and elongated N-terminal β-strand that are predicted to favor the formation of an atypical calcium-binding site at its tip (Elledge et al., 2010; Sotomayor et al., 2010). CDHR5 is most similar to PCDH15, and features cysteine residues predicted to form a disulfide bond at its tip (Sotomayor et al., 2014, 2012). In addition, a mutation in CDHR5, R84G (residue numbering throughout the text corresponds to processed proteins, see Methods), which mimics a deafness-related mutation in PCDH15, interferes with the intermicrovillar links formed by PCDH24 and CDHR5 (Crawley et al., 2014b), suggesting that these proteins interact in a similar fashion to the heterophilic tip-link “handshake” formed by CDH23 and PCDH15 (Behrendt et al., 2012; Sotomayor et al., 2014, 2012).

To better understand how PCDH24 and CDHR5 interact to form intermicrovillar links, we used a combination of structural and bead aggregation assays to characterize the adhesive properties and mechanisms of these cadherin family members. We present the X-ray crystallographic structures of *Homo sapiens* (*hs*) PCDH24 EC1-2, *Mus musculus* (*mm*) PCDH24 EC1-3, and *hs* CDHR5 EC1-2 refined at 2.3 Å, 2.1 Å, and 1.9 Å resolution, respectively. The structures give insight into possible binding mechanisms among species, which are further probed by bead aggregation assays revealing that the human and mouse intermicrovillar cadherins do not engage in the same homophilic and heterophilic interactions. Based on these results, we suggest models for the PCDH24/CDHR5 heterophilic interaction utilized to form intermicrovillar links.

## RESULTS

### PCDH24 and CDHR5 tip sequences are poorly conserved across species

Intermicrovillar links formed by PCDH24 and CDHR5 in the enterocyte brush border are similar to inner-ear tip links formed by CDH23 and PCDH15. Both types of links are essential for the development, assembly, and function of actin-based structures (microvilli in the gut and stereocilia in the inner ear), and both are made of long cadherin proteins that have similar cytoplasmic partners involved in interactions with the cytoskeleton and in signaling (Crawley et al., 2014a; Pepermans and Petit, 2015). Sequence analyses have revealed similarities between the tips of PCDH24 and CDH23, and between the tips of CDHR5 and PCDH15 (Sotomayor et al., 2014). Given that CDH23 and PCDH15 engage in a heterophilic “handshake” complex involving their EC1-2 tips, it has been proposed that PCDH24 and CDHR5 might use a similar binding mechanism (Crawley et al., 2014b; Sotomayor et al., 2014). To further explore this hypothesis, we performed sequence analyses comparing the tips of these cadherins to each other, to classical cadherins, and across species.

Interestingly, alignments of EC repeat sequences across up to 20 species for intermicrovillar and tip-link cadherins reveal poor conservation for PCDH24 and CDHR5 when compared to CDH23 and PCDH15. Average percent identity is 51.6% for CDH23 EC repeats and 45.5% for PCDH15 repeats, with the N- and C-terminal ends being more conserved than the middle EC repeats (Choudhary et al., 2019; Jaiganesh et al., 2018). In contrast, average percent identity is 16.4% for PCDH24 and 11% for CDHR5, considerably lower when compared to values for CDH23 and PCDH15. Moreover, the N- and C-terminal ends of PCDH24 and CDHR5 tend to be less conserved than the middle region of these proteins (Figure 1A and Figure S1 and S2; Tables S1-S5). Nevertheless, multiple sequence alignments of PCDH24 and CDH23 EC1 repeats confirm that PCDH24 has an elongated N-terminus that should facilitate the formation of calcium-binding site 0 as observed in structures of CDH23 (Figure S3A). Binding of calcium at this site in CDH23 is mediated by several acidic residues, which are also present and conserved in PCDH24. Similarly, sequence alignments of CDHR5 and PCDH15 EC1 repeats confirm that CDHR5 has the conserved cysteine residues that form a stabilizing disulfide bond at the tip of PCDH15 EC1 (Figure S3B). None of these proteins feature the tryptophan residues that mediate homophilic binding in classical cadherins (Brasch et al., 2012; Parisini et al., 2007; Shapiro et al., 1995), supporting the hypothesis that PCDH24 and CDHR5 might form a tip-link-like handshake complex mediating adhesion.

**Figure 1.**
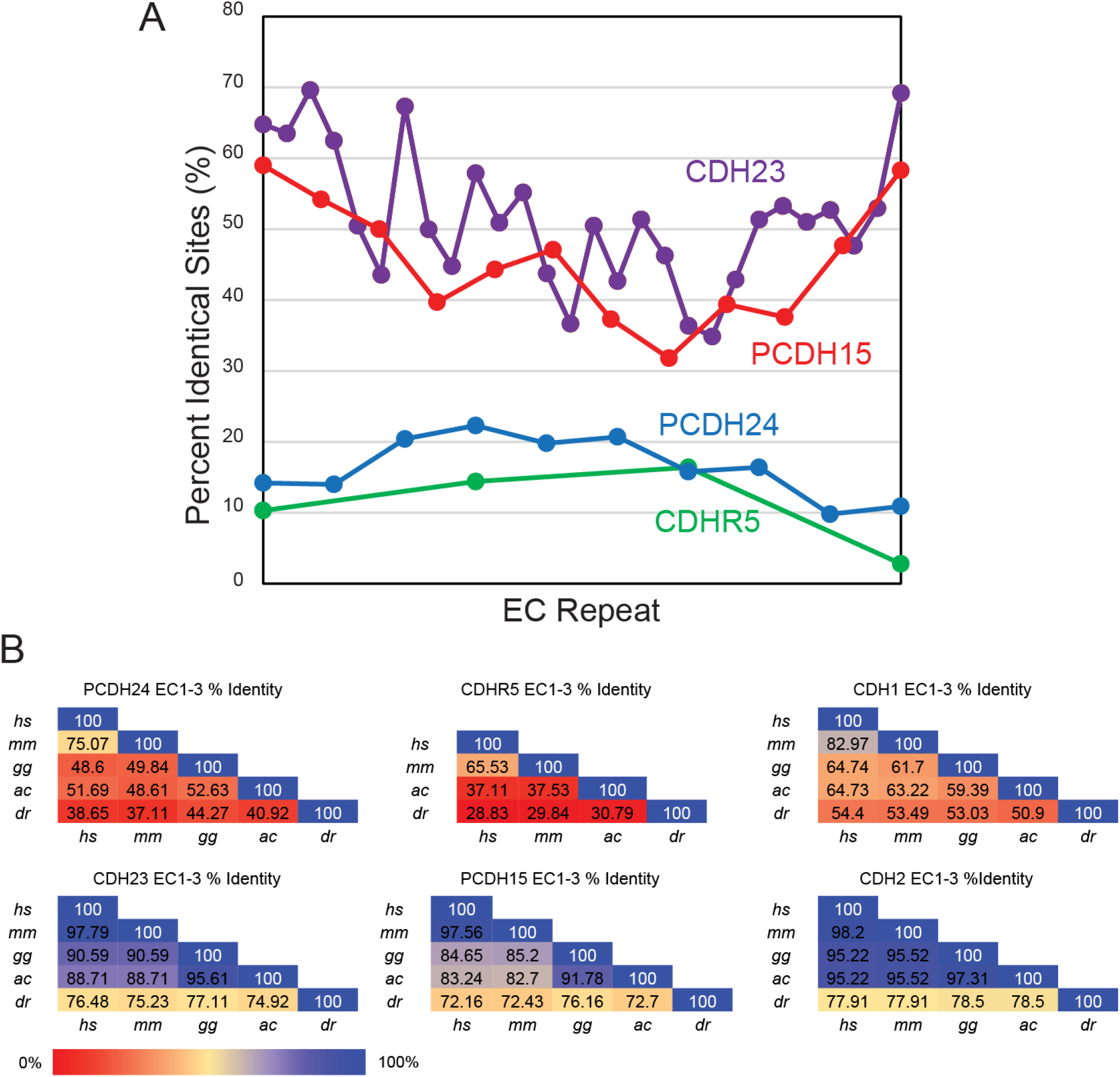
Comparison of sequence conservation across species. (A) Percent identity across the extracellular domains of PCDH24 (EC1-MAD10), CDH23 (EC1-MAD28), CDHR5 (EC1-4), and PCDH15 (EC1-MAD12) plotted against EC repeat numbers. Overall the cadherins of the inner-ear tip link, CDH23 and PCDH15, have higher identity across a variety of species while the intermicrovillar-link cadherins, PCDH24 and CDHR5, have poor sequence conservation across species (Figure S1, S2; Tables S2-S5). N- and C-termini are more conserved in CDH23 and PCDH15 than the middle region of these proteins. An opposite trend is observed for PCDH24 and CDHR5. (B) Percent identity of the first three EC repeats of PCDH24, CDHR5, CDH23, PCDH15, CDH1 and CDH2 demonstrate that PCDH24 and CDHR5 sequences are not highly conserved across several species (*Homo sapiens, hs*; *Mus musculus, mm*; *Gallus gallus, gg*; *Anolis carolinesis, ac*; *Danio rerio, dr*). However, the N-terminal repeats of CDH23, PCDH15, CDH1, and CDH2 are highly conserved. Sequences were obtained from NCBI (Table S6 and Methods), except for *gg* CDHR5, which is unavailable.

Pairwise sequence identity computed across up to five species for the N-terminal repeats (EC1-3) of PCDH24, CDHR5, CDH23, PCDH15, and the classical cadherins E-cadherin (CDH1) and N-cadherin (CDH2), also show different trends of sequence conservation (Figure 1B and Table S6). Average percent identity in pairwise comparisons is high for CDH2 (89%), CDH23 (86%), and PCDH15 (82%), moderate for CDH1 (61%) and lowest for PCDH24 (49%) and CDHR5 (38%). This is evident when comparing sequences from the most evolutionary distant species (human and fish). For instance, *hs* and *Danio rerio (dr)* PCDH24 EC1-3 sequences are 39% identical, while *hs* and *dr* CDH1 EC1-3 sequences are 54% identical. Sequence differences are still large for the human and mouse PCDH24 and CDHR5 protein tips, with 75% identity for the *hs* an *mm* PCDH24 EC1-3 sequences and 66% identity for the *hs* and *mm* CDHR5 EC1-3 sequences (compared to CDH1, CDH2, CDH23, and PCDH15 EC1-3 sequences with *hs* an *mm* pairwise identities between 83% and 98%) (Figure 1B). Proteins with sequence identity as low as 30% can share similar folds (Gilson et al., 2017), yet the low PCDH24 and CDHR5 sequence conservation in multiple sequence alignments and pair-wise comparisons suggests that details of their structures and function differ across species.

### Structures of human and mouse PCDH24 tips reveal N-terminal conformational variability

Previous bead aggregation assays using the *hs* PCDH24 protein revealed that its full-length extracellular domain can mediate adhesion through homophilic interactions with itself and through heterophilic interactions with *hs* CDHR5 (Crawley et al., 2014b). Members of the cadherin superfamily of proteins often engage in homophilic and heterophilic interactions using their N-terminal tips (EC1 for classical cadherins, EC1-2 for tip-link cadherins, and EC1-4 for clustered, δ1, and δ2 protocadherins) (Cooper et al., 2016; Goodman et al., 2016; Harrison et al., 2020; Modak and Sotomayor, 2019; Nagar et al., 1996; Nicoludis et al., 2016, 2015; Rubinstein et al., 2015; Schreiner and Weiner, 2010; Shapiro et al., 1995; Sotomayor et al., 2012; Takeichi, 1990). Therefore, to gain insights into the molecular basis of PCDH24-mediated adhesion and to explore the consequences of poor sequence conservation across species, we solved the crystal structures of *hs* PCDH24 EC1-2 and *mm* PCDH24 EC1-3 refined at 2.3 Å and 2.1 Å resolution, respectively (Table 1). These structures, obtained from crystals of proteins expressed in *E. coli* and refolded from inclusion bodies (see Methods), revealed both expected and unexpected architectural features.

**Table 1.** Statistics for PCDH24 and CDHR5 structures

| **Table 1. Statistics for PCDH24 and CDHR5 structures** |  |  |  |
| --- | --- | --- | --- |
| **Data Collection and Refinement** | ***Hs* PCDH24 EC1-2** | ***Mm* PCDH24 EC1-3** | ***Hs* CDHR5 EC1-2** |
| Space Group | P2<sub>1</sub> | P321 | I222 |
| Unit cell parameters |  |  |  |
| a, b, c (Å) | 270.9, 119.8, 74.3 | 74.8, 74.8, 140.0 | 50.4, 78.4, 121.4 |
| α, β, γ (°) | 90, 104.12, 90 | 90, 90, 120 | 90, 90, 90 |
| Molecules per asymmetric unit | 4 | 1 | 1 |
| Beam Source | APS 24-ID-C | APS 24-ID-C | APS 24-ID-E |
| Wavelength (Å) | 0.97910 | 0.97910 | 0.97918 |
| Resolution limit (Å) | 2.3 | 2.1 | 1.9 |
| Unique Reflections | 50945 (2128) | 27040 (1237) | 19398 (943) |
| Redundancy | 3.1 (2.6) | 8.60 (5.20) | 24.10 (21.80) |
| Completeness (%) | 95.5 (80.3) | 99.2 (90.6) | 99.7 (99.7) |
| Average I/σ (I) | 10.9 (3) | 27 (4.4) | 46.14 (14.50) |
| R<sub>merge</sub> | 0.105 (0.255) | 0.078 (0.337) | 0.074 (0.263) |
| **Refinement** |  |  |  |
| Resolution range (Å) | 50 – 2.3 (2.34 – 2.3) | 50 – 2.1 (2.14 – 2.1) | 50-1.9 (1.93-1.9) |
| Residues (atoms) | 861 (6610) | 318 (2447) | 204 (1656) |
| Water Molecules | 238 | 98 | 128 |
| R<sub>work</sub> (%) | 20.5 (27.8) | 21.0 (34.3) | 19.7 (20.9) |
| R<sub>free</sub> (%) | 23.5 (33.0) | 24.3 (39.6) | 23.4 (25.9) |
| Rms deviations |  |  |  |
| Bond lengths (Å) | 0.013 | 0.014 | 0.013 |
| Bond angles (°) | 1.460 | 1.632 | 1.652 |
| B factor average |  |  |  |
| Protein | 41.00 | 64.31 | 30.53 |
| Ligand/ion | 36.90 | 51.49 | 39.42 |
| Water | 31.65 | 50.99 | 33.6 |
| **Ramachandran plot regions** |  |  |  |
| Most favored (%) | 91.5 | 90.7 | 91.2 |
| Additionally allowed (%) | 8.3 | 8.9 | 8.8 |
| Generously allowed (%) | 0.3 | 0.4 | 0.0 |
| Disallowed (%) | 0.0 | 0.0 | 0.0 |
| **PDB ID code** | **5CZR** | **5CYX** | **6OAE** |

The *hs* PCDH24 EC1-2 structure has four molecules in the asymmetric unit including residues N1 to P215 for each protomer, while the *mm* PCDH24 EC1-3 structure has one molecule in the asymmetric unit including residues N1 to T324. Both structures had good-quality electron density maps that allowed the unambiguous positioning of side chains for most of the residues (Figure S4A-B). As expected, all EC repeats in the PCDH24 structures adopt the typical seven β-strand Greek-key fold seen in extracellular domains of cadherin proteins, with β-strands labeled A to G for each repeat (Figure 2A-D). Both *hs* and *mm* PCDH24 EC2 repeats have extended F and G strands, which are linked by a disulfide bond (S187:S201) that stabilizes an unusually long FG β-hairpin near the EC1-2 linker (Figure 2A-D, Figure S4A).

**Figure 2.**
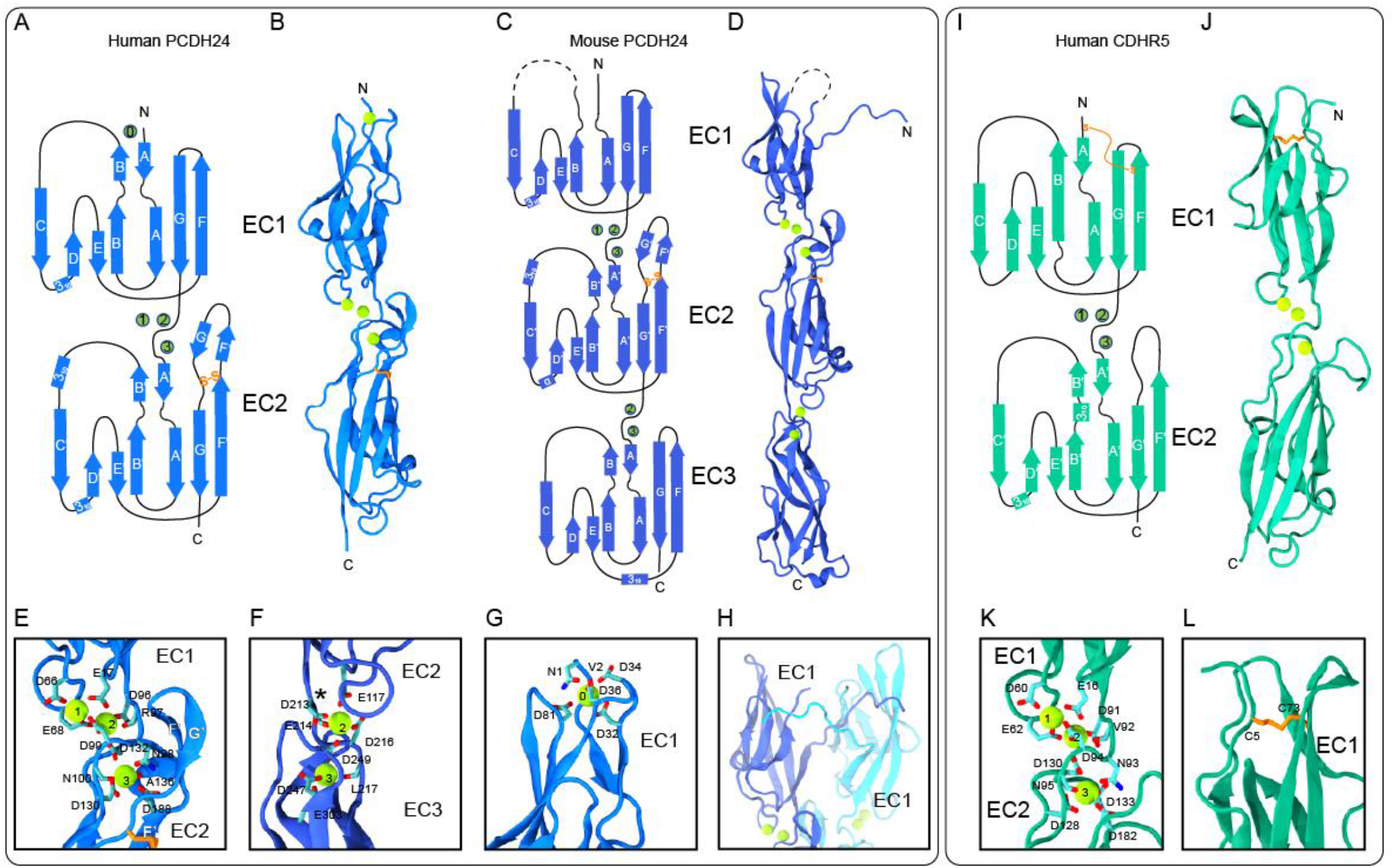
Structures of human (*hs*) PCDH24 EC1-2, mouse (*mm*) PCDH24 EC1-3, and human (*hs*) CDHR5 EC1-2. (A) Topology of *hs* PCDH24 EC1-2. The EC1 and EC2 repeats have a typical cadherin fold with 7 β-strands. A disulfide bond in EC2 is highlighted in orange. Calcium ions are shown as green circles. (B) A ribbon representation of *hs* PCDH24 EC1-2 with calcium ions in green. (C) Topology of the *mm* PCDH24 EC1-3 shown as in (A). The dashed line indicates a loop not resolved. (D) A ribbon representation of *mm* PCDH24 EC1-3 as in (B). The structure is similar to *hs* PCDH24 EC1-2, but the N-terminus projects away from EC1. A N-terminal calcium ion was not present. (E) Detail of the linker between EC1 and EC2 of *hs* PCDH24 showing canonical calcium-binding sites. Some sidechain and backbone atoms are not shown for clarity. (F) Detail of the *mm* PCDH24 EC2-3 linker, which lacks several canonical calcium-binding residues and thus only has two calcium ions bound. Shown as in (E). An asterisk (*) indicates the position where a calcium residue would be found in the canonical linker. (G) Detail of the N-terminal calcium bound at the tip of EC1 in the *hs* PCDH24 EC1-2 structure. A similar calcium-binding motif is seen at the tip of CDH23. Residues coordinating calcium are shown in stick with backbone atoms omitted. (H) Detail of the N-terminus of *mm* PCDH24 EC1-3 (blue) showing its interaction with another protomer (cyan) in the asymmetric unit. (I) Topology of *hs* CDHR5 EC1-2 shown as in (A). (J) A ribbon representation of *hs* CDHR5 EC1-2 with calcium ions in green. (K) Detail of the linker between EC1 and EC2 of *hs* CDHR5 showing canonical calcium-binding sites. Shown as in (I-J). (L) Detail of the disulfide bond in EC1 in the *hs* CDHR5 EC1-2 structure. A similar disulfide bond is seen in PCDH15 EC1.

The PCDH24 EC1-2 linker region is canonical in both human and mouse structures and features calcium ions bound to sites 1, 2, and 3, which are coordinated by acidic residues from the cadherin motif NTerm-XEX-DXD-D(R/Y)(D/E)-XDX-DXNDN-CTerm (Figure 2A-E and Figure S5). Interestingly, the structure of mouse PCDH24 has a non-canonical calcium-binding linker between repeats EC2 and EC3, with only two bound calcium ions and a straight conformation (Figure 2C-D, F and Figure S5). This linker lacks necessary conserved residues to bind the calcium ion that would occupy site 1 as the canonical DXE sequence is replaced with residues 172SYN174. Although the canonical DXNDN sequence motif is 213DXPDL217 at the EC2-3 linker, typical coordination of calcium at site 3 by the asparagine carboxamide oxygen is replaced by coordination through the backbone carbonyl group of a glutamate residue in our structure (E214 in mouse, Q214 in human). The last residue of the motif (typically asparagine, replaced here by L217) coordinates calcium at site 3 through its backbone carbonyl group, an interaction that is unchanged by residue variability at this position. Sequence conservation across species for the DXE/SYN and DXNDN/DXPDL motifs suggest that most species have a non-canonical linker region between repeats EC2 and EC3 with two bound calcium ions as observed in our *mm* PCDH24 EC1-3 structure. While non-canonical linker regions with two or less bound calcium ions have been shown to be flexible (Araya-Secchi et al., 2016; Jin et al., 2012; Powers et al., 2017), equilibrium and steered molecular dynamics (SMD) simulations of *mm* PCDH24 EC1-3 show that rigidity and mechanical strength are not compromised in this non-canonical linker (Figure S6).

As expected for members of the Cr-2 family of non-clustered protocadherins, the *hs* PCDH24 EC1-2 structure shows an occupied site 0 calcium-binding site at the tip of EC1, with calcium-coordinating residues similar to those observed in structures of CDH23 (1N, 32DXDXD36, 80XDX^TOP^; Figure 2A-B, G, Figure S7A). Unexpectedly, the *mm* PCDH24 EC1-3 structure shows an unoccupied site 0, with an extended N-terminus protruding away from EC1 as observed for classical cadherins that exchange their N-terminal strands to form adhesive bonds (Figure 2C-D, H, Figure S7B). The sequence motifs involved in calcium coordination at site 0 are conserved, suggesting that crystallization conditions, including 3M LiCl, facilitated the opening of this site. Alternatively, other differences between mouse and human sequences might be responsible for the structural divergence. Most notably, residue R86 near the FG loop and the XDX^TOP^ motif of *mm* PCDH24 EC1 seems to be incompatible with a closed N-terminal conformation and an occupied calcium-binding site 0. The open and closed conformations observed for PCDH24 EC1 will likely determine the type of homophilic or heterophilic interactions that mediate adhesion.

### Crystal contacts in human and mouse PCDH24 structures suggest distinct adhesive interfaces

Crystallographic contacts observed in cadherin structures have revealed multiple physiologically relevant interfaces (Brasch et al., 2011; Cooper et al., 2016; Goodman et al., 2016; Modak and Sotomayor, 2019; Nicoludis et al., 2016). The crystallographic packing observed for the *hs* and *mm* PCDH24 structures shows various interfaces that further highlight differences across species. The asymmetric unit of the *hs* PCDH24 EC1-2 structure shows 4 molecules with two similar anti-parallel *trans* dimers (Figure S8A) arranged perpendicular to one another to form a dimer of dimers (Figure S8B). Each antiparallel dimer positions EC1 from one protomer in front of EC2 from the next protomer, slightly shifted with respect to each other and with the extended FG β-hairpins near the EC1-2 linker regions mediating the interaction head to head. The antiparallel dimers have interface areas of 979.2 Å^2^ and 938.6 Å^2^ for protomers A:D and B:C, respectively. These values are both larger than an empirical threshold (856 Å^2^) that distinguishes biologically relevant interactions from crystallographic packing artifacts (Ponstingl et al., 2000). Two other interfaces are small and unlikely to be biologically relevant (Figure S8C-D). Previous data has shown that *hs* PCDH24 mediates homophilic *trans* adhesion when *hs* CDHR5 is not present (Crawley et al., 2014b). It is possible that the antiparallel EC1-2 interface facilitates homophilic adhesion mediated by *hs* PCDH24, yet our SMD simulations suggest that this interface is weak (Figure S9).

In contrast to the crystal contacts observed in the *hs* PCDH24 EC1-2 structure, the crystal packing of *mm* PCDH24 EC1-3 (space group *P*321) results in multiple large interfaces generated from rotations of a single molecule in the asymmetric unit. A parallel trimer is present around the axis of symmetry, with the EC1 N-termini from each protomer protruding away and inserting into pockets of adjacent EC1 repeats in a clockwise fashion, thereby interlocking the EC1 domains (Figure S10A-B). This is possible because of the open conformation observed for the mouse N-terminus of EC1 without a bound calcium ion at site 0, and it is reminiscent of how the EC1 N-terminus of classical cadherins is exchanged resulting in the insertion of W2 into an hydrophobic binding pocket of the partner molecule (Figure S7C-D) (Parisini et al., 2007; Shapiro et al., 1995). Interestingly, a glycosylation site is predicted at N9, which might interfere with or regulate the formation of interlocking EC1 repeats, as has been observed for other cadherins (Brasch et al., 2011).

The interlocking of mouse EC1 repeats in the trimer facilitates the formation of two parallel *cis* interfaces between EC1-3 protomers involving the FG β-hairpin interacting with the neighboring EC2-3 linker. These *cis* interfaces are large with an interface area of 1352.6 Å^2^ between two monomers (Figure S10A). In addition, a fully overlapped antiparallel *trans* interface is observed between EC1-3 protomers with an area of 1221.2 Å^2^ (Figure S10C). Sugars at a second predicted glycosylation site located in the center of the *cis* trimer (p.N161), and at a third predicted glycosylation site near the *trans* interface between EC1 and EC3 (p.N280), may interfere with or regulate the formation of these interfaces. Together, the *cis* trimer and *trans* dimer could form a glycosylation-modulated interlocking arrangement of PCDH24 molecules that is different from the *trans* interaction in the *hs* PCDH24 EC1-2 structure. Sequence conservation mapped to the *mm* EC1-3 structure reveals that only core residues are highly conserved, highlighting poor conservation for surface exposed residues potentially involved in adhesive interactions (Figure S11). Motivated by the sequence and structural differences observed for the human and mouse PCDH24 tips, we used bead aggregation assays to determine the minimum adhesive unit of PCDH24 homophilic binding in both species.

### Human, but not mouse PCDH24 mediates homophilic adhesion

To determine which EC repeats are essential for *hs* and *mm* PCDH24 homophilic adhesion we created a library of constructs encoding for the full-length extracellular domains (EC1-MAD10) and various truncations (EC1-7, EC1-6, EC1-5, EC1-4, EC1-3, EC1-2, and EC1), all fused to a C-terminal Fc tag. These protein fragments were produced in HEK293T cells and used for aggregation assays with Protein G magnetic beads (see Methods).

As expected, bead aggregation experiments using *hs* PCDH24 EC1-MAD10Fc showed large calcium-dependent aggregates (Figure 3A, L-M, Figure S15A, I). Similar large calcium-dependent aggregates were seen for *hs* PCDH24 EC1-7Fc through EC1-3Fc (Figure 3B-F, L-M, Figure S15B-F). Further truncations indicate that smaller *hs* PCDH24 fragments, including just EC1, are capable of facilitating the formation of small bead aggregates that increase in size with rocking but never reach the larger sizes observed when using fragments with 3 or more EC repeats (Figure 3G-H, K-M, Figure S15G-H). Controls performed using EDTA showed that constructs are unable to aggregate when calcium is depleted, except when using EC1Fc, possibly stabilized by the adjacent Fc-tag (Figure 3I-M, Figure S15I). These results suggest that *hs* PCDH24 mediates calcium-dependent homophilic adhesion, with at least three distinct modes of adhesion, two calcium-dependent mediated by *hs* PCDH24 EC1-3 and PCDH24 EC1-2, and another one calcium-independent mediated by *hs* PCDH24 EC1 alone. It is possible that each of these modes uses a different interface, and that the antiparallel interface observed in our *hs* PCDH24 EC1-2 structure is a low-affinity intermediate state that facilitates further interdigitation for the adhesive mode that drives the formation of larger bead aggregates.

**Figure 3.**
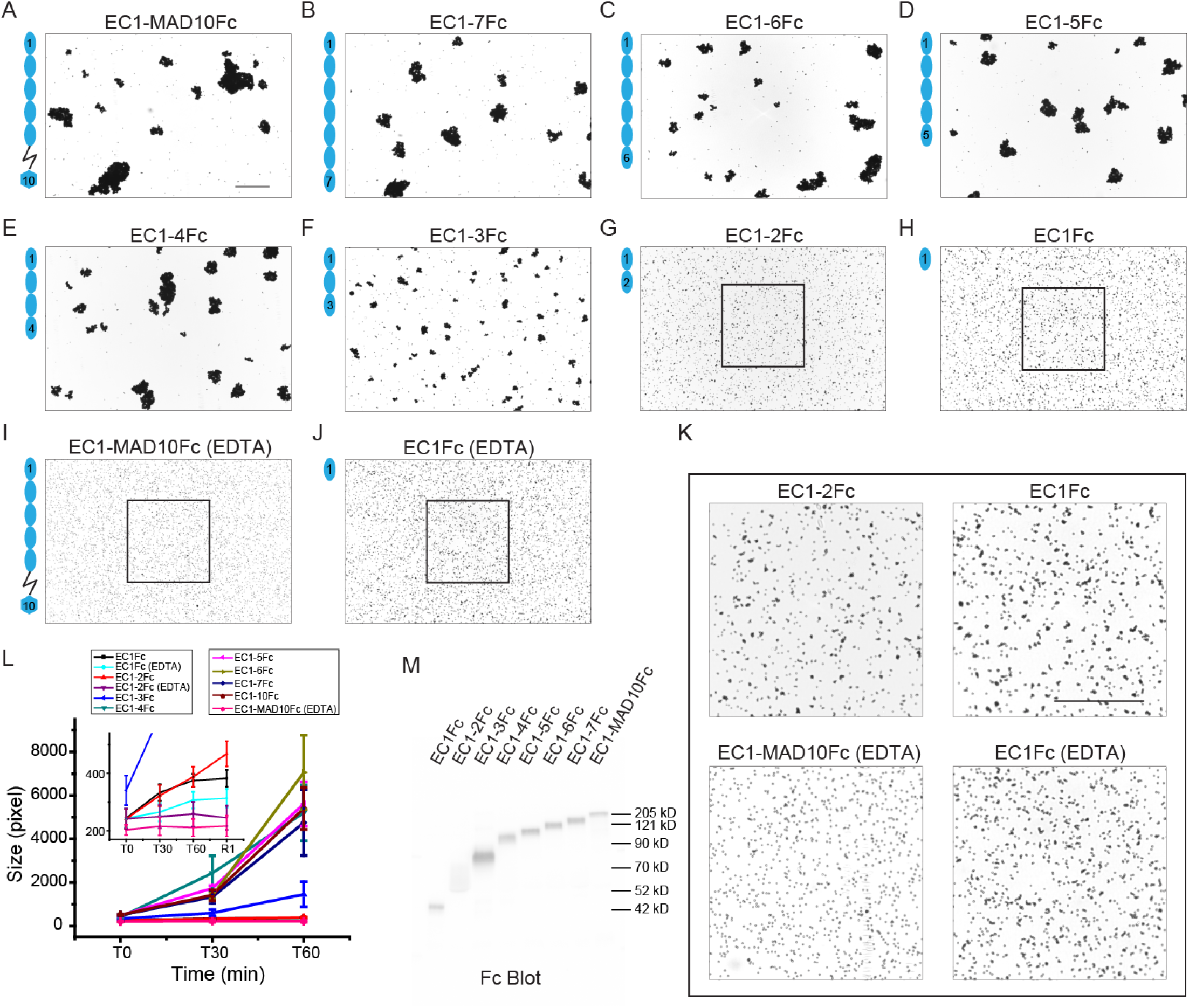
Homophilic binding assays of *hs* PCDH24. (A-H) Protein G beads coated with the full-length *hs* PCDH24 extracellular domain (A) and its C-terminal truncation versions (B-H). Each image shows bead aggregation observed after 60 min in the presence of 2 mM CaCl_2_. (I-J) Protein G beads coated with the full-length *hs* PCDH24 extracellular domain (I) and EC1 (J) in the presence of 2 mM EDTA, shown as in (A-H). (K) Detail of binding assay results for EC1-2Fc, EC1Fc, EC1-MAD10Fc with EDTA, and EC1Fc with EDTA. Aggregation is still present for the shortest constructs. Bar – 500 µm. (L) Aggregate size for full-length and truncated versions of *hs* PCDH24 extracellular domains at the start of the experiment (T0), and after 30 and 60 min (T30, T60). Inset shows aggregate size of the shortest constructs compared to the EDTA control of EC1-MAD10Fc. An additional one-minute rocking step is denoted by R1. Error bars are standard error of the mean (*n* indicated in Table S7). (M) Western blot shows expression and secretion of full-length and truncated versions of *hs* PCDH24.

Given the low sequence identity between the human and mouse N-terminal EC repeats of PCDH24 when compared to CDH23, and the structural differences observed between *hs* PCDH24 EC1-2 and *mm* PCDH24 EC1-3, bead aggregation assays were also performed with the full-length extracellular domain of *mm* PCDH24. Bead aggregation was not observed for *mm* PCDH24 EC1-MAD10Fc (Figure 4A-B, E, G). These results suggest that the large crystallographic antiparallel *trans* interface formed by PCDH24 EC1-3 (Figure S10C) does not mediate bead aggregation under the conditions tested and reveal that PCDH24 mediated homophilic adhesion is species-dependent.

**Figure 4.**
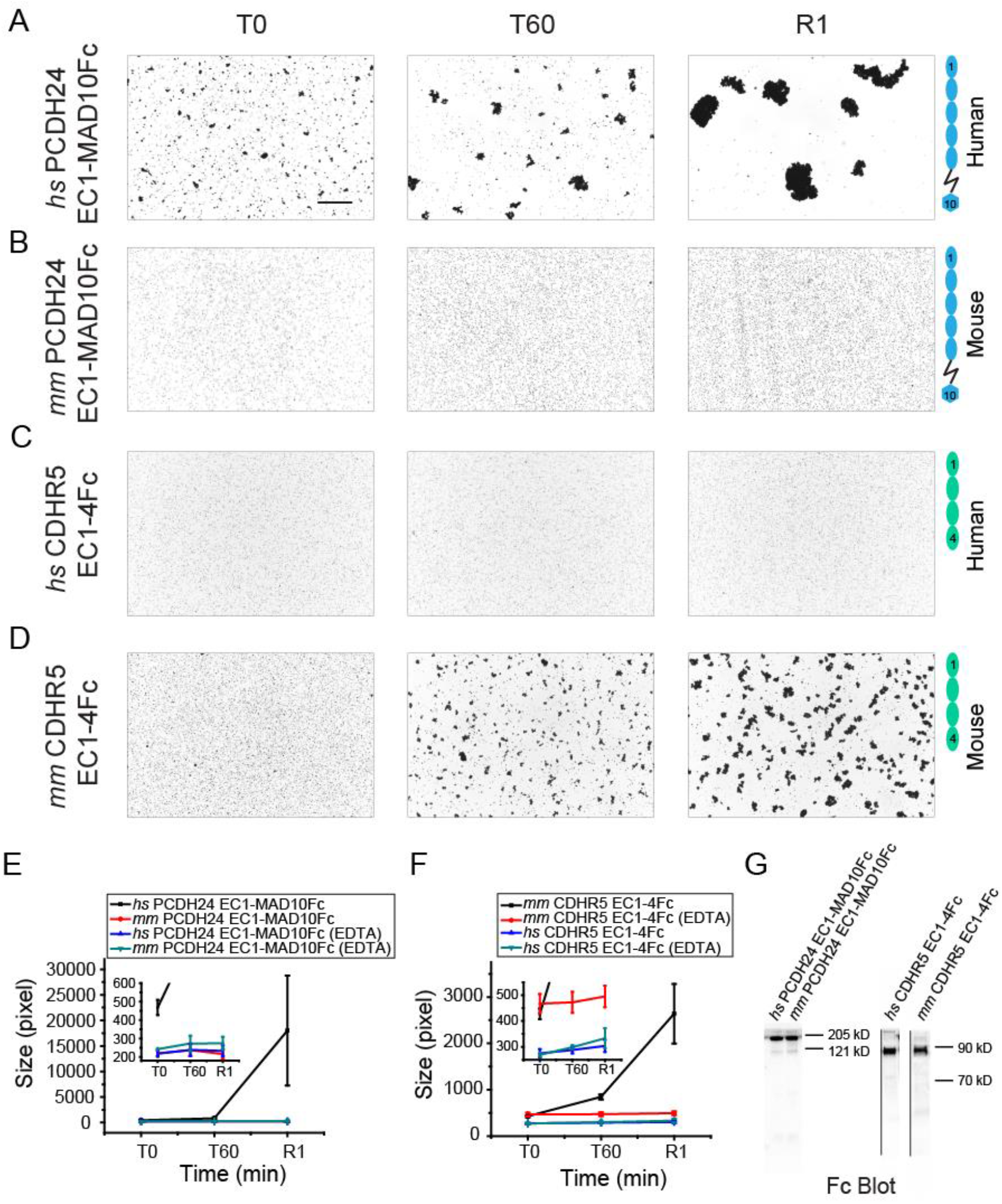
Comparison of homophilic binding assays of *hs* and *mm* PCDH24 and CDHR5. (A-D) Protein G beads coated with full-length *hs* PCDH24 (A), full-length *mm* PCDH24 (B), full-length *hs* CDHR5 (C), and full-length *mm* CDHR5 (D) cadherin extracellular domains, including MAD10 for PCDH24. None of the CDHR5 fragments included the MLD. Images show the aggregation observed at the start of the experiment (T0), after 60 min (T60) followed by rocking for 1 min (R1) in the presence of 2 mM CaCl_2_. Bar – 500 µm. (E) Aggregate size for full-length *hs* and *mm* PCDH24 extracellular domains in the presence of 2 mM CaCl_2_ and 2 mM EDTA at the start of the experiment (T0), after 60 min (T60) followed by rocking for 1 minute (R1). Inset shows aggregate size of full-length *mm* PCDH24 extracellular domain compared to the EDTA control of full-length *hs* and *mm* PCDH24 extracellular domains. (F) Aggregate size for full-length *hs* and *mm* CDHR5 cadherin extracellular domains in the presence of 2 mM CaCl_2_ and 2 mM EDTA at the start of the experiment (T0), after 60 min (T60) followed by rocking for 1 min (R1). Inset shows aggregate size of the full-length *hs* CDHR5 cadherin extracellular domain compared to the EDTA control of full-length *hs* and *mm* CDHR5 cadherin extracellular domain. Error bars in (E) and (F) are standard error of the mean (*n* indicated in Table S7). (G) Western blot shows expression and secretion of full-length *hs* and *mm* PCDH24 and CDHR5 cadherin extracellular domains.

### Mouse, but not human CDHR5 mediates homophilic adhesion

The full-length extracellular domain of *hs* CDHR5 was previously shown to not mediate homophilic adhesion (Crawley et al., 2014b). The low sequence identity between the human and the mouse CDHR5, especially at their N-termini, and the species-dependent homophilic adhesive behavior of PCDH24, prompted us to test if *mm* CDHR5 could mediate adhesion. Full-length cadherin extracellular domains (without MLDs) of both human and mouse CDHR5 were used for bead aggregation assays. As expected, *hs* CDHR5 EC1-4Fc did not mediate bead aggregation in the presence and the absence of calcium. However, *mm* CDHR5 EC1-4Fc did mediate bead aggregation in the presence of calcium (Figure 4C-D, F-G). These results suggest that, like PCDH24, CDHR5 homophilic adhesiveness is species-dependent.

To determine the minimal adhesive unit of *mm* CDHR5, a library of truncated cadherin fragments was generated and used for bead aggregation assays. Calcium-dependent bead aggregation was observed when using beads coated with *mm* CDHR5 EC1-4Fc and EC1-3Fc, but not when using EC1-2Fc and EC1Fc (Figure 5A-D, G-I, Figure S16A-E). Hence, *mm* CDHR5 EC1-3 is sufficient to mediate homophilic adhesion.

**Figure 5.**
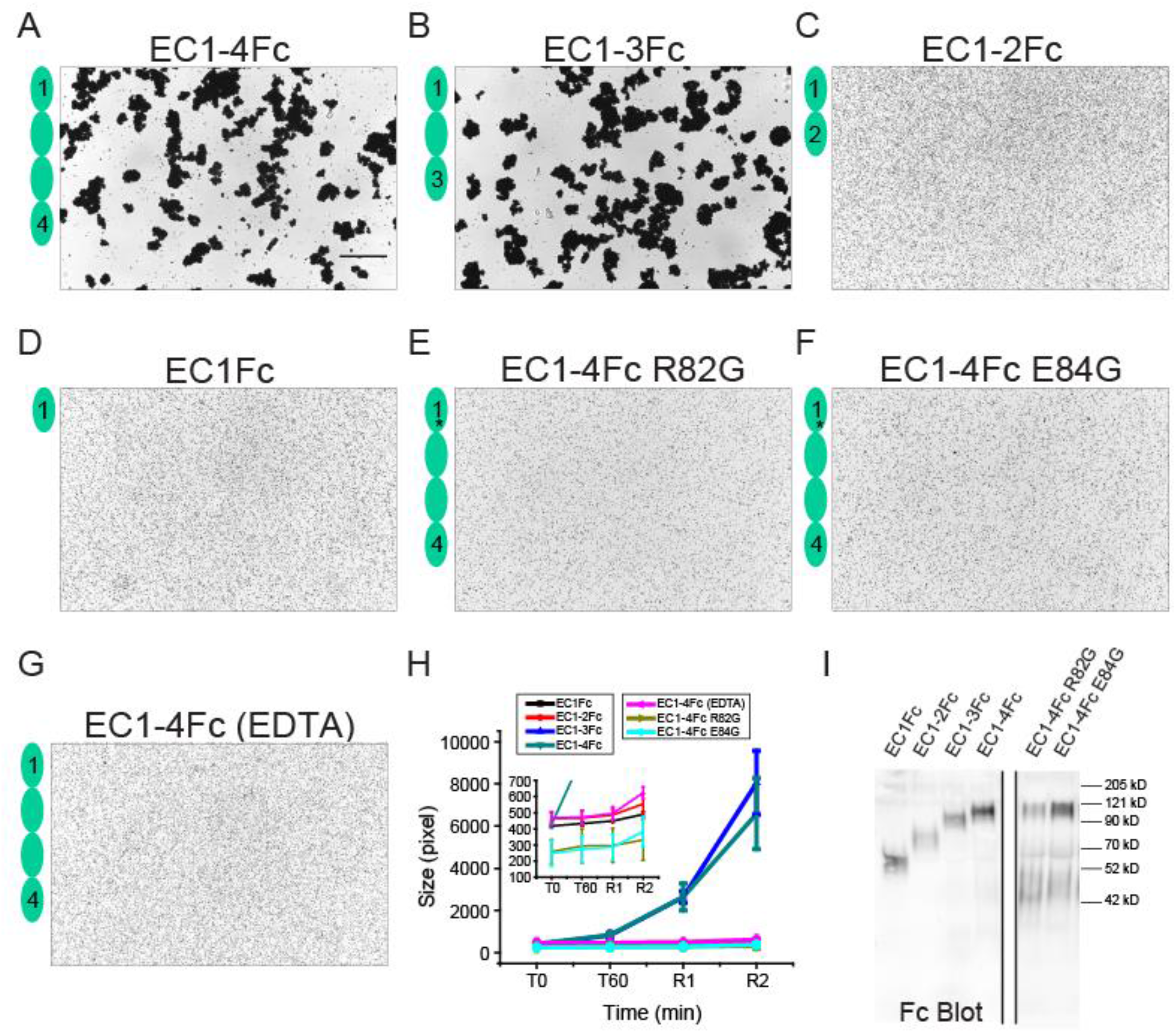
Homophilic binding assays of *mm* CDHR5. (A-F) Protein G beads coated with the full-length *mm* CDHR5 cadherin extracellular domain (A) and its C-terminal truncation versions (B-D). Each image shows bead aggregation observed after 60 min followed by rocking for 2 min in the presence of 2 mM CaCl_2_. (E-F) Protein G beads coated with mutants *mm* CDHR5 EC1-4Fc R82G (E) and EC1-4Fc E84G (F) observed after 60 min followed by rocking for 2 min in the presence of 2 mM CaCl_2_. (G) Protein G beads coated with the full-length *mm* CDHR5 cadherin extracellular domain in the presence of 2 mM EDTA shown as in (A). Bar – 500 µm. (H) Aggregate size for full-length, truncated versions and mutants of *mm* CDHR5 cadherin extracellular domains at the start of the experiment (T0), after 60 min (T60) followed by rocking for 1 min (R1) and 2 min (R2). Inset shows aggregate size of EC1Fc, EC1-2Fc, the EDTA control of EC1-4Fc and EC1-4Fc mutants R82G and E84G. Error bars are standard error of the mean (*n* indicated in Table S7). (I) Western blot shows expression and secretion of full-length, truncated, and mutant versions of *mm* CDHR5 cadherin extracellular domains.

To provide insight into the interface used by *mm* CDHR5 to mediate homophilic adhesion, we tested mutations that alter a site similar to that mutated in *hs* CDHR5 (R84G) preventing heterophilic binding to *hs* PCDH24. Bead aggregation assays were performed with both *mm* CDHR5 EC1-4Fc R82G and E84G, and both mutants abolished bead aggregation of *mm* CDHR5 EC1-4Fc in the presence of calcium (Figure 5E-F, H-I). These results suggest that the interface used for mediating heterophilic adhesion between PCDH24 and CDHR5 and the interface used for mediating homophilic adhesion by *mm* CDHR5 might be similar.

### Structure of human CDHR5 EC1-2 reveals a PCDH15-like fold and suggests adhesive interfaces

To gain insight into CDHR5’s adhesive function we solved the crystal structure of *hs* CDHR5 EC1-2 refined at 1.9 Å resolution (Table 1). This structure, obtained using protein expressed in *E. coli* and refolded from inclusion bodies (see Methods), had one molecule in the asymmetric unit including residues Q1 to L207, with clear electron density for residues A2 to A205 (Figure 2I, J and Figure S4C). Like other cadherins, the structure of *hs* CDHR5 EC1-2 shows straight consecutive EC repeats, each with a typical seven β-strand Greek-key fold (Figure 2I, J). The linker region between the repeats has the canonical acidic residues that coordinate calcium and three calcium ions bound to it (Figure 2K and Figure S12). Similar to PCDH15, a member of the Cr-3 subfamily of non-clustered protocadherins, *hs* CDHR5 EC1 has a disulfide bridge between cysteine residues C5 and C73 on strands A and F respectively (Figure 2L, Figure S4C, S7E). This disulfide bond keeps the N-terminal strand tucked to the protomer in a closed conformation, reminiscent of how calcium at site 0 facilitates closing of the N-terminal strand in EC1 repeats of CDH23 and *hs* PCDH24.

Next, we analyzed crystallographic contacts in the *hs* CDHR5 EC1-2 structure (Figure S13), which could lead to an understanding of how mouse, but not human CDHR5 mediates homophilic adhesion and how the heterophilic complex with PCDH24 is formed. The two largest contacts within the *hs* CDHR5 EC1-2 crystal structure are interfaces in antiparallel arrangements. The largest, at 1433.9 Å^2^, is formed by interactions between the EC1 repeat of one protomer and EC2 of the adjacent one, with a slight shift in the register of the antiparallel dimeric interface (Figure S13A). There are up to three predicted N-glycosylation sites at this interface in the sequences for the human (p.N19 and p.N56) and the mouse (p.N17, p.N56, and p.N107) proteins. The next largest contact involves interactions between antiparallel EC2 repeats facing each other (Figure S13B), with an interface area of 530.2 Å^2^, small relative to likely physiologically relevant interfaces (>856 Å^2^), but comparable to interface area per EC repeat reported for physiologically relevant contacts in protocadherins (Cooper et al., 2016; Modak and Sotomayor, 2019; Nicoludis et al., 2019). In this arrangement, the EC3 repeat (not present in the structure), would interact with EC1, thus increasing the total contact area. None of the predicted N-glycosylation sites for the human or mouse sequences are at the EC2 interface of this *trans* dimer. An X-shaped crystallographic contact is also present (Figure S13C), with a small interface area (364.7 Å^2^) involving EC1 to EC1 interactions. Additional smaller interfaces are present but are unlikely to be physiologically relevant given their atypical arrangement and small interface areas of 347.9 Å^2^, 132.7 Å^2^, and 114.0 Å^2^, respectively (Figure S13D-F).

Although, *hs* CDHR5 does not mediate adhesion in bead aggregation assays, some of the contacts discussed above can serve as templates to predict the interfaces that could be used by *mm* CDHR5 to mediate homophilic adhesion. The largest, antiparallel EC1-2 interface might be impaired by glycosylation at a site predicted to exist in the mouse sequence (N107), but not in the human protein (R107). In addition, bead aggregation assays suggest that repeat EC3 is needed for adhesive interactions, suggesting that this EC1-2 interface is not driving CDHR5 homophilic adhesion. In contrast, the antiparallel EC1-3 interface mediated by EC2 interactions is compatible with results from bead aggregation assays indicating that the first three N-terminal repeats are sufficient for *trans* adhesiveness, but details of the arrangement are likely to be different due to the absence of EC3 in our structure and the low sequence conservation of surface residues (Figure S14). These contacts may also guide the search for interaction modes that facilitate the formation of PCDH24 and CDHR5 heterophilic complexes.

### Human and mouse heterophilic PCDH24 and CDHR5 bonds are distinct

Previous bead aggregation assays showed that full-length extracellular domains of human PCDH24 and CDHR5 mediate heterophilic adhesion (Crawley et al., 2014b). We used our library of Fc-tagged PCDH24 and CDHR5 cadherin fragments to test for heterophilic adhesiveness for both human and mouse proteins. To distinguish PCDH24 from CDHR5 coated beads, we used green and red fluorescent beads, respectively. We confirmed that human PCDH24 and CDHR5 cadherin extracellular domains mediate calcium-dependent heterophilic adhesion, and found that the mouse proteins also facilitate calcium-dependent bead aggregation (Figure 6A-B, I, J, K and Figure S17A-B), suggesting that the heterophilic intermicrovillar cadherin bond driving intestinal brush border assembly is species independent.

**Figure 6.**
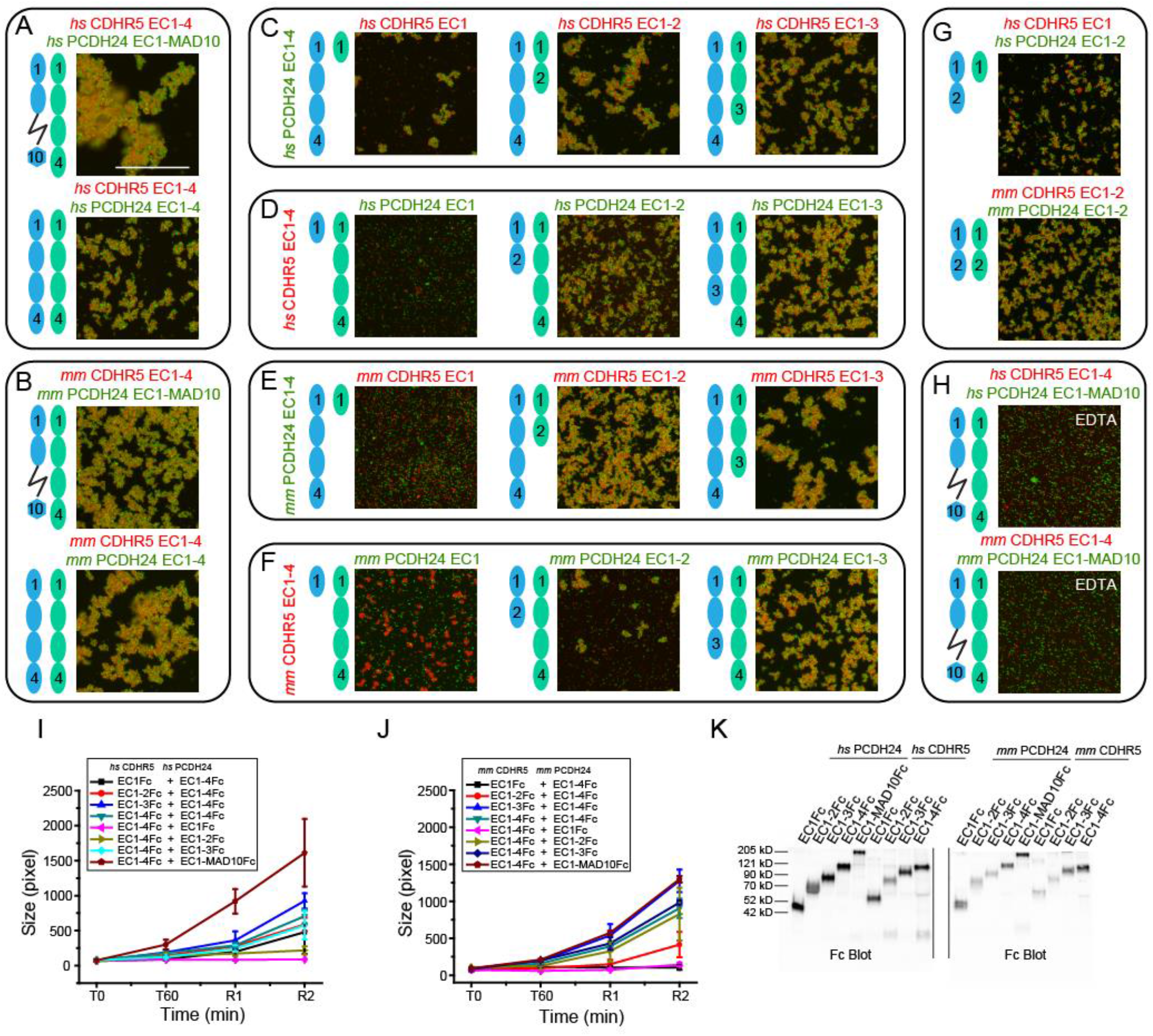
Heterophilic binding assays of *hs* and *mm* PCDH24 and CDHR5. (A-B) Images from binding assays of *hs* PCDH24 (full-length extracellular domain and EC1-4Fc) mixed with the full-length *hs* CDHR5 cadherin extracellular domain (A), and *mm* PCDH24 (full-length extracellular domain and EC1-4Fc) mixed with full-length *mm* CDHR5 cadherin extracellular domain (B). Green fluorescent protein A beads are coated with PCDH24 fragments and red fluorescent protein A beads are coated with CDHR5 in all panels. All images are details of complete views shown in Figure S17. Images show bead aggregation observed after 60 min followed by rocking for 2 min in the presence of 2 mM CaCl_2_. (C-F) Images from binding assays of truncations of *hs* CDHR5 mixed with *hs* PCDH24 EC1-4Fc (C), truncations of *hs* PCDH24 mixed with *hs* CDHR5 EC1-4Fc (D), truncations of *mm* CDHR5 mixed with *mm* PCDH24 EC1-4Fc (E), and truncations of *mm* PCDH24 mixed with *mm* CDHR5 EC1-4Fc (F). Images show bead aggregation observed after 60 min followed by rocking for 2 min in the presence of 2 mM CaCl_2_. (G) Images from binding assays of minimum EC repeats required for heterophilic adhesion of *hs* and *mm* PCDH24 and CDHR5. The minimum units for heterophilic adhesion for the human proteins are CDHR5 EC1Fc and PCDH24 EC1-2Fc. The minimum units for heterophilic adhesion for the mouse proteins are CDHR5 EC1-2Fc and PCDH24 EC1-2Fc. Images show bead aggregation observed after 60 minutes followed by rocking for 2 minutes in the presence of 2 mM CaCl_2_. (H) Protein A beads coated with full-length *hs* and *mm* PCDH24 and CDHR5 cadherin extracellular domains (including MAD10 for PCDH24 and without the MLD for CDHR5) in the presence of 2 mM EDTA shown as in (A-B). Bar – 500 µm. (I) Aggregate size for full-length and truncated constructs of *hs* PCDH24 and *hs* CDHR5 cadherin extracellular domains (including MAD10 for PCDH24 when indicated and without the mucin-like domain for CDHR5) at the start of the experiment (T0), after 60 min (T60) followed by rocking for 1 min (R1) and 2 min (R2). (J) Aggregate size for full-length and truncated constructs of *mm* PCDH24 and *mm* CDHR5 as in (I). Error bars in I and J are standard error of the mean (*n* indicated in Table S7). (K) Western blot shows expression and secretion of full-length and truncated versions of *hs* and *mm* PCDH24 and CDHR5 cadherin extracellular domains (including MAD10 for PCDH24 when indicated and without the MLD for CDHR5).

Next, we used truncated protein fragments and found that repeats EC1-4Fc of PCDH24 and CDHR5 from both species are sufficient to drive heterophilic bead aggregation (Figure 6A-B, I, J, K and Figure S17A-B). Further bead aggregation assays focused on determining the minimum adhesive units for heterophilic adhesion. We carried out heterophilic binding assays using PCDH24 EC1-4Fc mixed with truncated fragments of CDHR5 and vice versa for both human and mouse proteins (Figure 6C-F, I, J, K and Figure S17C-F). The minimal adhesive units for heterophilic binding when using human proteins were PCDH24 EC1-2Fc and CDHR5 EC1Fc, whereas mouse proteins required PCDH24 EC1-2Fc and CDHR5 EC1-2Fc.

Additional experiments using only the required EC repeats of PCDH24 and CDHR5 to test heterophilic adhesiveness for both human and mouse proteins confirmed the results obtained with the truncation series (Figure 6G, K and Figure S17G, I). Binding assays in the presence of 2 mM EDTA abolished aggregation indicating that heterophilic adhesion mediated by PCDH24 and CDHR5 is calcium-dependent (Figure 6H, K and Figure S17H, J). The low sequence similarity of CDHR5 and PCDH24 across different species may explain the differences in their binding mechanisms, as demonstrated by the distinct minimal adhesive units found for the human and mouse heterophilic complexes. Remarkably, while homophilic adhesive behavior is not conserved between human and mouse, the heterophilic bond is robust with slight differences in underlying molecular mechanisms.

## DISCUSSION

Our sequence analyses, structures, and bead aggregation assays suggest that there are various modes of *trans* homophilic and heterophilic interactions involving the extracellular domains of PCDH24 and CDHR5 (Figure 7). We have confirmed that human PCDH24 can form homophilic bonds (Crawley et al., 2014b), and found three potentially distinct modes of *trans* adhesion in bead aggregations assays. The first mode involves weak (small bead aggregates) calcium-independent interactions mediated by EC1. This adhesive mode is not strong enough to drive calcium-dependent PCDH24 adhesion when using larger parts of its extracellular domain, and therefore might be unphysiological or relevant only when a preformed bond is exposed to calcium chelators. The physiological PCDH24 *trans* homophilic bond might be initiated by weak interactions involving EC1 and EC2 repeats (small bead aggregates), with further interdigitation for stronger interactions (larger bead aggregates) mediated by at least EC1 to EC3 repeats.

**Figure 7.**
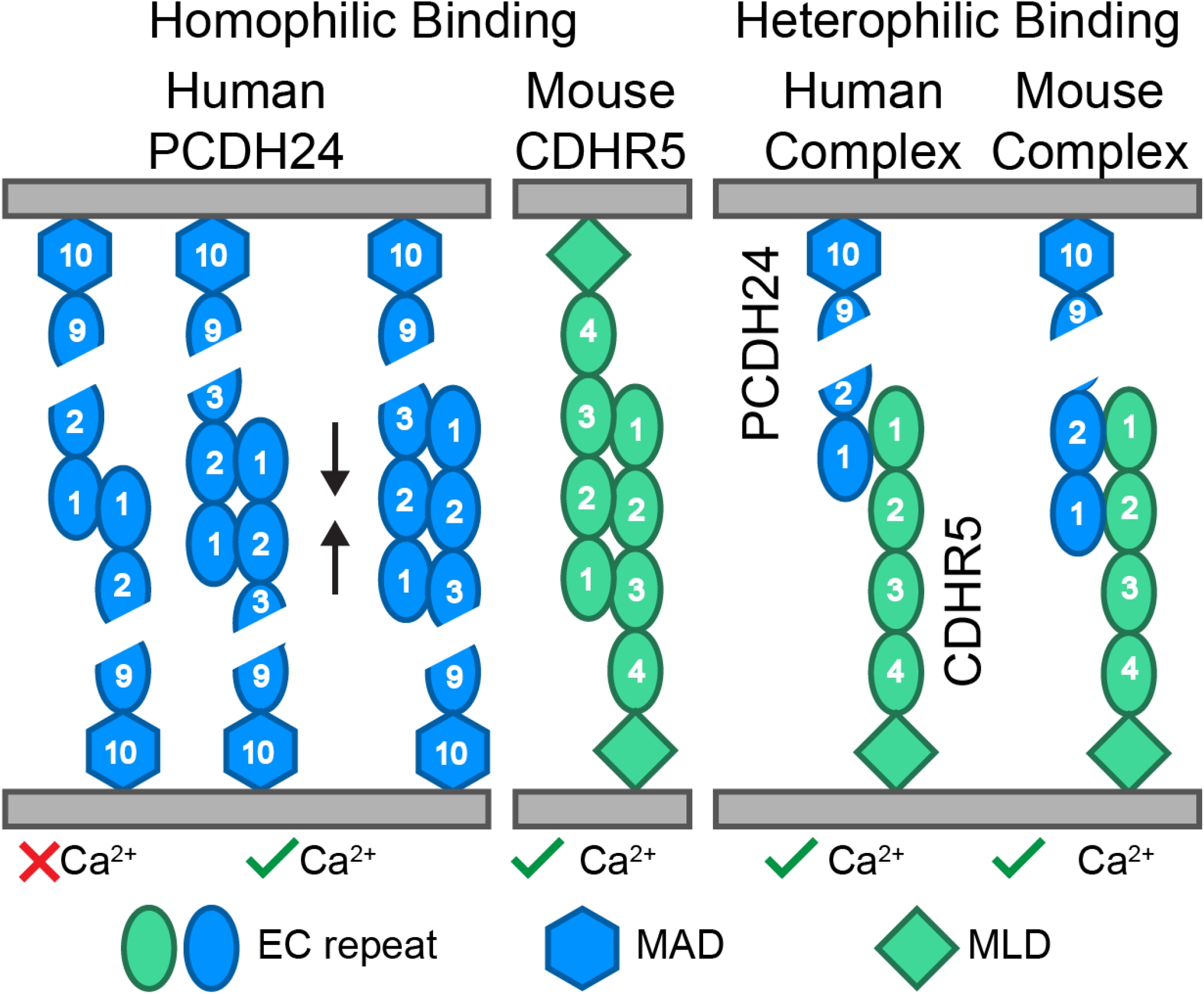
Schematics of homophilic and heterophilic adhesion mediated by human and mouse PCDH24 and CDHR5. Homophilic adhesion that results in small bead aggregates can be mediated by *hs* PCDH24 EC1 (calcium independent) and EC1-2 (calcium dependent), while homophilic adhesion that results in large bead aggregates requires *hs* PCDH24 EC1-3 (calcium dependent). Homophilic adhesion mediated by *mm* CDHR5 requires EC1-3 (calcium dependent). Heterophilic adhesion is mediated by PCDH24 EC1-2 and CDHR5 EC1 when using human proteins and by PCDH24 EC1-2 and CDHR5 EC1-2 when using mouse proteins (both calcium dependent).

Intriguingly, our bead aggregation assays indicate that mouse PCDH24 does not mediate *trans* homophilic adhesion, highlighting a species-dependent behavior that is consistent with poor sequence and structural conservation for the mouse and human proteins. In contrast, experiments with CDHR5 reveal that the mouse protein can mediate *trans* homophilic adhesion, but the human protein cannot. These results are consistent with L929 cell aggregation assays that showed homophilic adhesion mediated by the rat CDHR5 protein (Goldberg et al., 2002, 2000). The physiological relevance of the *trans* homophilic bonds for human PCDH24 and mouse CDHR5 remains to be determined. Mice lacking PCDH24 expression in the intestinal track are viable, but intestinal tissue exhibits various structural defects leading to functional impairment of intestine function and loss of body weight (Pinette et al., 2018). Homophilic CDHR5 bonds might be partially compensating for the lack of heterophilic intermicrovillar links in these mice.

Importantly, adhesion mediated by PCDH24 and CDHR5 heterophilic bonds is species independent in our bead aggregation assays with mouse and human proteins. These results also demonstrate that lack of homophilic adhesion for mouse PCDH24 and for human CDHR5 is not caused by issues with bead coating or with protein misfolding and degradation in our assays. Details of the heterophilic interaction between PCDH24 and CDHR5, however, might differ from species to species, as we found that the minimal adhesive units are PCDH24 EC1-2Fc / CDHR5 EC1Fc for human proteins and PCDH24 EC1-2Fc / CDHR5 EC1-2Fc for mouse proteins.

Our structures of the human PCDH24 EC1-2, mouse PCDH24 EC1-3, and human CDHR5 EC1-2 protein fragments confirm the expected structural similarities between the PCDH24 and CDH23 tips, and between the PCDH15 and CDHR5 tips. These structures also support binding mechanisms that are distinct from those used by classical cadherins, clustered protocadherins, and non-clustered δ protocadherins (Brasch et al., 2019, 2012; Cooper et al., 2016; Goodman et al., 2016; Harrison et al., 2020; Lefebvre, 2017; Modak and Sotomayor, 2019; Nicoludis et al., 2016, 2015; Rubinstein et al., 2015). While crystal contacts in our structures suggest possible interfaces that might facilitate the modes of adhesion discovered through bead aggregation assays, further work will be required to elucidate and validate structural models of *trans* homophilic and heterophilic complexes, to understand the relevance of an *in crystallo cis* complex observed here for mouse PCDH24, to clarify the structural role played by glycosylation *in vivo* (Krahn et al., 2010), and to determine which complexes form intermicrovillar links in enterocytes.

Intermicrovillar links observed in human CACO-2_BBE_ cells and in native mouse tissue are 46.8 ± 8.9 nm and 49.9 ± 8.8 nm in length, respectively (Crawley et al., 2014b). Interestingly, lengths range from 30 to 80 nm for both human and mouse links, perhaps reflecting the challenges of measuring link sizes from two-dimensional electron-microscopy images of cells, with some additional uncertainty coming from difficult-to-define link edges. Nevertheless, this range of lengths hints at multiple possibilities for the architecture of the complex that are consistent with the modes of adhesion we found using bead aggregation assays. Links made of two PCDH24 molecules interacting in *trans* and tip-to-tip could reach lengths of up to ∼90 nm (9 EC repeats + MAD10 per molecule; 4.5 nm long each), while links made of the shortest CDHR5 isoforms with just 4 EC repeats interacting tip-to-tip could be 36 nm in length or even shorter if there is any overlap at the junction or bending at any of the EC linkers. Heterophilic links formed by a tip-to-tip *trans* interaction of the full-length PCDH24 ectodomain and the shortest CDHR5 isoform should be ∼63 nm or shorter. The average value observed *ex vivo* and *in vivo* (∼48 nm) indicates that some overlap at the junction may exist, and that most of the links are likely formed by the heterophilic complex. The minimal adhesive units we found for mouse and human PCDH24 and CDHR5 homophilic and heterophilic *trans* complexes are consistent with these estimates for length ranges, assuming that up to 3 EC repeats might be overlapping to form robust bonds that lead to bead aggregation *in vitro*, and adhesion *in vivo*.

The functional role of PCDH24 and CDHR5 in non-intestinal tissues and cells is unclear. These proteins have also been found in the liver and kidney (Goldberg et al., 2000; Okazaki et al., 2002) and have been implicated in contact inhibition of cell proliferation (Okazaki et al., 2002; Ose et al., 2009), morphogenesis (Behrendt et al., 2012; Goldberg et al., 2000; Krahn et al., 2010), colorectal carcinogenesis (Bujko et al., 2015; Losi and Grande, 2014), and gallstone disease (Chuang et al., 2011). Exploring how different organs in different species express and use PCDH24 and CDHR5 proteins, which have poorly conserved sequences, might provide with a unique opportunity to reveal how evolutionary events structurally encode adhesive and signaling functions in various physiological contexts (Dickinson et al., 2011). It is tempting to speculate that differences in gut proteases’ specificity across species, as well as diverse diets and gut microbiota, put divergent selective pressures driving the molecular evolution of intermicrovillar links. Similarly, other selective pressures relevant to specific organs might be driving the evolution of heterophilic and homophilic PCDH24 and CDHR5 adhesive complexes.

Interestingly, both brush border enterocytes and inner-ear sensory receptor cells use heterophilic cadherin complexes in actin-based structures relevant for their specialized functions. These heterophilic cadherin complexes link membranes within cells, as opposed to mostly homophilic cadherin complexes that mediate adhesion between membranes of adjacent cells. Heterophilic complexes, with a pair of distinct cytoplasmic domains, might be more functionally versatile in signaling, as the number and type of cytoplasmic partners can be independently diversified for each member of the complex. The strength and dynamics of cadherin heterophilic bonds, which are present in systems subjected to mechanical stimuli, might also differ from and provide specific advantages over weak homophilic cadherin complexes that cluster to provide strong adhesiveness. Whether PCDH24 and CDHR5 engage in *cis* interactions that can lead to more mechanically robust intermicrovillar links, as observed for tip links (Choudhary et al., 2019; Dionne et al., 2018; Ge et al., 2018), or to the formation of larger adhesive complexes in non-microvillar membranes, as observed for classical cadherins and clustered protocadherins (Brasch et al., 2019, 2012; Schreiner and Weiner, 2010), remains to be explored.

## METHODS

### Cloning, expression, and purification of PCDH24 and CDHR5 fragments for crystallography

DNA encoding for *hs* PCDH24 EC1-2, *mm* PCDH24 EC1-3 and *hs* CDHR5 EC1-2 were sub cloned into *NdeI* and *XhoI* sites of the pET21a vector including a fused C-terminal hexa-histidine tag. All DNA constructs were sequence verified. These constructs were used to transform BL21(DE3) pLysS cells (Stratagene). The PCDH24 transformants were cultured in TB and induced at OD_600_ ∼ 0.6 with 1 mM IPTG at 37**°**C overnight. The *hs* CDHR5 EC1-2 transformant was cultured in TB and induced at an OD_600_ ∼ 0.6 with 1 mM IPTG at 30°C overnight. Cells were lysed by sonication in denaturing buffer (20 mM TrisHCl [pH7.5], 6 M guanidine hydrochloride [GuHCl], 10 mM CaCl_2_ and 20 mM imidazole). The cleared lysates were loaded onto Ni-Sepharose beads and nutated for ∼1 h (GE Healthcare). Beads were washed twice with denaturing buffer and hexa-histidine-tagged proteins were eluted with the same buffer supplemented with 500 mM imidazole. After purification, 2 mM DTT was added to the protein before refolding. The *hs* PCDH24 EC1-2 protein was refolded in six dialysis steps using a buffer with 20 mM TrisHCl [pH 8.0], 5 mM CaCl_2_ and decreasing amounts of GuHCl (6 M, 3 M, 2 M, 1 M, 0.5 M and 0 M). In steps four, five and six the refolding buffer included 400 mM arginine. Steps three through six also included 150 mM NaCl. The *mm* PCDH24 EC1-3 and *hs* CDHR5 EC1-2 proteins were refolded overnight using a buffer with 400 mM Arginine, 20 mM TrisHCl [pH 8.0], 150 mM NaCl, and 5 mM CaCl_2_. All dialysis steps were performed using MWCO 2000 membranes and used refolding buffers for 12 hours at 4**°**C. Refolded protein was further purified on a Superdex200 16/600 column (GE Healthcare) in 20 mM TrisHCl [pH 8.0], 150 mM NaCl and 2 mM CaCl_2_, and concentrated by ultrafiltration to ∼5 mg/ml for crystallization (Vivaspin 10 KDa).

### Crystallization, data collection, and structure determination

Crystals were grown by vapor diffusion at 4°C by mixing 1 μL of protein and 0.5 μL of reservoir solution including (0.2 M MgCl, 0.1 M HEPES [pH 7.3], 10% PEG 4000) for *hs* PCDH24 EC1-2, (3.0 M LiCl, 0.1 M HEPES [pH 7.5]) for *mm* PCDH24 EC1-3 and (0.1 M Na Acetate pH 4.4, 0.7 M CaCl_2_) for *hs* CDHR5 EC1-2. Crystals were cryo-protected in reservoir solution plus a cryoprotectant (25% PEG 400 for PCDH24 and 25% glycerol for *hs* CDHR5 EC1-2) and cryo-cooled in liquid N_2_. X-ray diffraction datasets were collected as indicated in Table 1 and processed with HKL2000 (Otwinowski and Minor, 1997).

All structures were solved by molecular replacement using PHASER (McCoy et al., 2007). The first two N-terminal repeats of CDH23 EC1-2 (PDB: 4AQE) (Sotomayor et al., 2012) were used as an initial model for the *mm* PCDH24 EC1-3 structure. The *mm* PCDH24 EC3 repeat was built using BUCCANEER (Cowtan, 2007, 2006). The *hs* PCDH24 EC1-2 structure was determined by using the first two N-terminal EC repeats of the *mm* PCDH24 EC1-3 structure (PDB: 5CYX). For the *hs* CDHR5 EC1-2 structure we used a model of CDH23 EC22-24 (PDB: 5UZ8) (Jaiganesh et al., 2018) created using RaptorX (Källberg et al., 2012). Model building was done with COOT (Emsley et al., 2010) and refinement was performed with REFMAC5 (Murshudov et al., 2011). Restrained TLS refinement was used for the PCDH24 structures. Data collection and refinement statistics are provided in Table 1.

### Cloning of Fc-fusions proteins for bead aggregation assays

DNA constructs encoding C-terminal truncations of *hs* and *mm* PCDH24 as well as of *hs* and *mm* CDHR5, all fused to a C-terminal Fc-tag, were prepared for bead aggregation assays. Truncations were made at DXNDN or similar ends of each EC repeat (Figure S1 and S2). Truncations were cloned into a modified CMV vector (Jontes laboratory, OSU) using *XhoI* and *KpnI* (*hs* PCDH24, *mm* PCDH24), *NheI* and *XhoI* (*hs* CDHR5), and *XhoI* and *BamHI* (*mm* CDHR5) cutsites. The sequence for the linker between the *hs* CDHR5 protein fragments and their Fc tags is LELKLRILQSTVPRARDPPV, while the sequence for the linker between the *mm* CDHR5 protein fragments and their Fc tags is TDPPV. Mutations were generated using the QuikChange Lightning kit (Agilent). All DNA constructs were sequence verified.

### Bead aggregation assays

Bead aggregation assays were performed as previously described (Cooper et al., 2016; Crawley et al., 2014b; Emond et al., 2011; Emond and Jontes, 2014; Modak and Sotomayor, 2019). The PCDH24Fc and CDHR5Fc fusion constructs were transfected into HEK293T cells via calcium-phosphate transfection (Barry et al., 2014; Jiang and Chen, 2006; Kwon and Firestein, 2013). A solution of 10 µg of DNA and 250 mM CaCl_2_ was added dropwise to 2X HBS with mild vortexing. The solution was immediately added drop-wise to a 100 mm dish of HEK293T cells cultured in DMEM with FBS, L-glut, and PenStrep. For each construct, two plates of cells were transfected. The following day, cells were rinsed twice with 1X PBS and allowed to grow in serum-free media for two days prior to collection. The collected media contained the secreted Fc fusion proteins and was concentrated (Amicon 10 kD, 30 kD concentrators) to a volume of 500 µL before being incubated with 1.5 µL of Protein G Dynabeads (Invitrogen). The beads were incubated with rotation for 2 hr at 4°C. Incubated beads were washed with binding buffer (50 mM Tris [pH 7.5], 100 mM NaCl, 10 mM KCl, and 0.2% BSA) and split into two tubes, either with 2 mM CaCl_2_ or 2 mM EDTA. Beads were then allowed to aggregate on a glass depression slide in a humidified chamber for 1 h, then rocked for 1 to 2 min at 8 oscillations/min. Images were taken at various time points using a Nikon Eclipse Ti microscope with a 10X objective as detailed in Table S7. Sizes of bead aggregates were quantified with ImageJ as described previously (Emond et al., 2011; Emond and Jontes, 2014). Images were thresholded to exclude background and the area of the detected particles was measured in pixels. Average aggregate size (± SEM) was plotted in Origin Pro. Number of independent biological replicates are listed in Table S7.

### Fluorescent bead aggregation assays

Fluorescently labeled protein A Dynabeads were produced for the heterophilic bead aggregation assays. Protein A (5 mg; Thermo Fisher Pierce) was resuspended in 1X PBS to obtain a final concentration of 2 mg/mL. Using the Thermo Fisher Dylight Antibody Labeling kits, 1 mg of Protein A was labeled with Dylight 488 to produce 0.5 moles of dye/1 mole of Protein A. Labeled Protein A (200 µg) was then conjugated to Dynabeads M-280 Tosylactivated (Invitrogen) following the provided protocol to produce 10 mg of beads at a final concentration of 20 mg/mL. A similar protocol was used to obtain Dylight 594 conjugated Protein A Dynabeads. Conjugation to the beads was confirmed by observing the beads in depression slides under 10X magnification with the GFP filter cube (Nikon) for the Dylight 488 fluorophore and the Texas Red filter cube (Nikon) for the Dylight 594 fluorophore. Beads of both colors were then mixed on a depression slide and no overlap of fluorescence was observed. Further confirmation that the beads were functional was done by performing a bead aggregation assay with a positive control (*hs* PCDH24 EC1-4).

Fluorescent bead aggregation assays were performed as done with non-fluorescent beads but with noted exceptions. Concentrated media was added to 2.2 µL of fluorescently labeled beads to achieve the same concentration as the Protein G Dynabeads on the slides. Dylight 488 conjugated beads were used for all PCDH24Fc fusions and Dylight 594 conjugated beads were used for all CDHR5Fc fusions. Prior to addition of beads to the CaCl_2_ or EDTA tubes, 160 µL of Dylight 488 beads containing the PCDH24 constructs were mixed with 160 µL Dylight 594 beads containing CDHR5 constructs, then 150 µL of the mixture was added to the CaCl_2_ or EDTA tubes.

### Western Blots

Western blots were performed for Fc-tagged proteins to confirm expression and secretion of the protein. Protein samples were mixed with 4X SDS with β-mercaptoethanol, boiled for 5 min, and loaded onto SDS-PAGE gels (BioRad) for electrophoresis. Proteins were electroblotted onto PVDF membrane (GE Healthcare), blocked with 5% nonfat milk in TBS with 0.1% Tween-20 for 1 h, and incubated overnight at 4°C with goat anti-human IgG (1:200; Jackson ImmunoResearch Laboratories, Catalog-109-025-003, Lot-117103, 135223). After washing with 1X TBS, the membranes were incubated with mouse anti-goat HRP-conjugated secondary antibody (1:5000; Santa Cruz Biotechnology, Catalog-sc-2354, Lot-A2017, J1718) for 1 h at room temperature. Membranes were washed in 1X TBS and developed using the ECL Select Western Blot Detection kit (GE Healthcare) for chemiluminescent detection (Omega Lum G). Western blots were performed on samples obtained immediately after media collection and concentration as well as protein samples obtained from the Protein G/A beads following aggregation assays.

### Sequence alignments

Sequence alignments were performed on individual EC repeats for a variety of species of PCDH24 (Table S2) and CDHR5 (Table S3) in MUSCLE (Edgar, 2004). Sequences that covered the complete extracellular domain were selected to include a variety of species of each protein. Sequence conservation per EC repeat in percent identity was obtained from the MUSCLE alignments in Geneious (Kearse et al., 2012) (Table S1). All EC repeats of both human and mouse PCDH24 (Figure S5) and CDHR5 (Figure S12) were aligned to one another as well as described above. Conservation of alignments were visualized in JalView (Waterhouse et al., 2009) using percent identity (Henikoff and Henikoff, 1992) with a conservation cutoff of 40%. Additionally, sequence alignments of individual EC repeats for a variety of species of PCDH15 (Choudhary et al., 2019) (Table S4) and CDH23 (Jaiganesh et al., 2018) (Table S5) were performed in MUSCLE, and sequence conservation per EC repeat was obtained in Geneious (Table S1). A separate set of alignments, which include up to 5 species for the N-terminal repeats (EC1-3) of PCDH24, CDHR5, CDH23, PCDH15, CDH1, and CDH2 (Table S6), were performed in MUSCLE. The Sequence Identity and Similarity Server (SIAS) was used to determine identity between pairs of different species for each protein (Figure 1B). A broader conservation analysis of PCDH24 EC1-3 and CDHR5 EC1-2 was done in Consurf (Ashkenazy et al., 2016, 2010; Celniker et al., 2013; Glaser et al., 2003; Landau et al., 2005) using additional sequences to include a larger variety of species (Table S8 for PCDH24 and Table S9 for CDHR5). The values obtained from Consurf (1 = lowest conservation, 9 = highest conservation) were matched to the corresponding residues in the structures and used to make a heat map of conservation. N-glycosylation and O-glycosylation sites for the extracellular domains of PCDH24 and CDHR5 were predicted using NetNGlyc 1.0 and NetOgly 4.0 (Steentoft et al., 2013) and highlighted in the multiple sequence alignment. All positives for O-glycosylation were labeled, but predictions for N-glycosylation were only labeled if the prediction received a 9/9 prediction score from the server.

### Simulated systems

The psfgen, solvate, and autoionize VMD (Humphrey et al., 1996) plugins were used to build all systems (Table S10), adding hydrogen atoms to the protein structure and crystallographic water molecules. For the *mm* PCDH24 EC1-3 systems, the missing loop in EC1 (residues 32 to 37) was built in COOT (Emsley et al., 2010) using the human PCDH24 EC1 structure as a template. Non-native N-terminal residues were removed. Residues D, E, K, and R were considered to be charged and neutral histidine protonation states were determined on favorable hydrogen bond formation. The proteins were placed in additional water and randomly placed ions to solvate and ionize the systems at 150 mM NaCl. For SMD simulations, protein molecules were aligned to the *x-*axis. Additional information for systems is presented in (Table S10).

### Simulations Parameters

Equilibrium and SMD simulations were performed using NAMD 2.11 (Phillips et al., 2005), the CHARMM36 forcefield for proteins with the CMAP correction, and the TIP3P model for water (Huang and MacKerell, 2014). Van der Waals interactions were computed with a cutoff at 12 Å and using a switching function starting at 10 Å. Computation of long-range electrostatic forces was done with the Particle Mesh Ewald method with a cutoff of 12 Å and a grid point density of >1 Å^-3^. A 2 fs uniform integration time was used with SHAKE. When indicated, constant temperature (*T* = 300K) was enforced using Langevin dynamics with a dampening coefficient of 0.1 ps^-1^ unless otherwise noted. The hybrid Nose-Hoover Langevin piston method was used to maintain constant pressure (*NpT*) at 1 atm with a 200 fs decay period and a 100 fs dampening time constant. In simulation S3a (Table S10), the *mm* PCDH24 EC1-3 protein had harmonic constraints (*k* = 1 kcal mol^-1^ Å^-2^) applied to the C_α_ atoms of residues S110, V123, and L182 in EC2 to avoid rotations leading to interactions between periodic images. Constant velocity stretching simulations (S1b-d and S3b-d, Table S10) were carried out using the SMD method and the NAMD Tcl interface (Franz et al., 2020; Grubmüller, 2005; Isralewitz et al., 2001; Sotomayor and Schulten, 2007). Virtual springs (*k*_s_ = 1 kcal mol^-1^ Å^-2^) were attached to Cα atoms of C-terminal ends for unbinding simulations (S1b-d) and to the center of mass of EC1 and EC3 for unfolding simulations (S3b-d). The free ends of the stretching springs were moved in opposite directions, away from the protein(s), at a constant velocity. Applied forces were computed using the extension of the virtual springs. Maximum force peaks and their averages were computed from 50 ps running averages.

## ACCESSION NUMBERS

Coordinates for *hs* PCDH24 EC1-2, *mm* PCDH24 EC1-3 and *hs* CDHR5 EC1-2 have been deposited in the Protein Data Bank with entry codes 5CZR, 5CYX, and 6OAE, respectively.

## ACKNOWLEDGEMENTS

We thank members of the Sotomayor laboratory for assistance and discussions. This work was supported in part by the Ohio State University and NIH (NIDDK R01 DK095811 to M.J.T.). Use of APS NE-CAT beamlines was supported by NIH (P41 GM103403 & S10 RR029205) and the Department of Energy (DE-AC02-06CH11357) through grants GUP 49774 and 59251. Use of TACC-Stampede and PSC-Bridges supercomputers was supported by the National Science Foundation through XSEDE (XRAC MCB140226). Use of the OSC-Owens supercomputer was supported by grant PAS1037. DM was a Pelotonia fellow and MS was an Alfred P. Sloan fellow (FR-2015-65794).

## AUTHOR CONTRIBUTIONS

M.E.G. did sequence alignments, solved the structure for *hs* CDHR5 EC1-2, did cloning of *hs* and *mm* PCDH24Fc constructs, performed homophilic binding assays of PCDH24, prepared the fluorescence magnetic beads for heterophilic bead aggregation assays, and did simulations. Z.R.J. did cloning, protein expression and protein purification, crystallization trials, and solved structures for *hs* PCDH24 EC1-2, *mm* PCDH24 EC1-3, and *hs* CDHR5 EC1-2. D.M. did cloning of *hs* and *mm* CDHR5Fc constructs, carried out homophilic bead aggregation assays of CDHR5 and did all heterophilic bead aggregation assays. M.J.T. provided guidance, reagents, and secured funding. M.S. trained M.E.G., Z.R.J., and D.M. and assisted with crystal fishing, cryo-cooling, solving crystal structures, and simulations. M.S. supervised work and assisted with data analysis. M.E.G., D.M., and M.S. prepared figures and wrote the manuscript with feedback from M.J.T. and Z.R.J.

## COMPETING INTERESTS

The authors declare no competing interests.

## SUPPLEMENTARY MATERIAL

**Figure S1.**
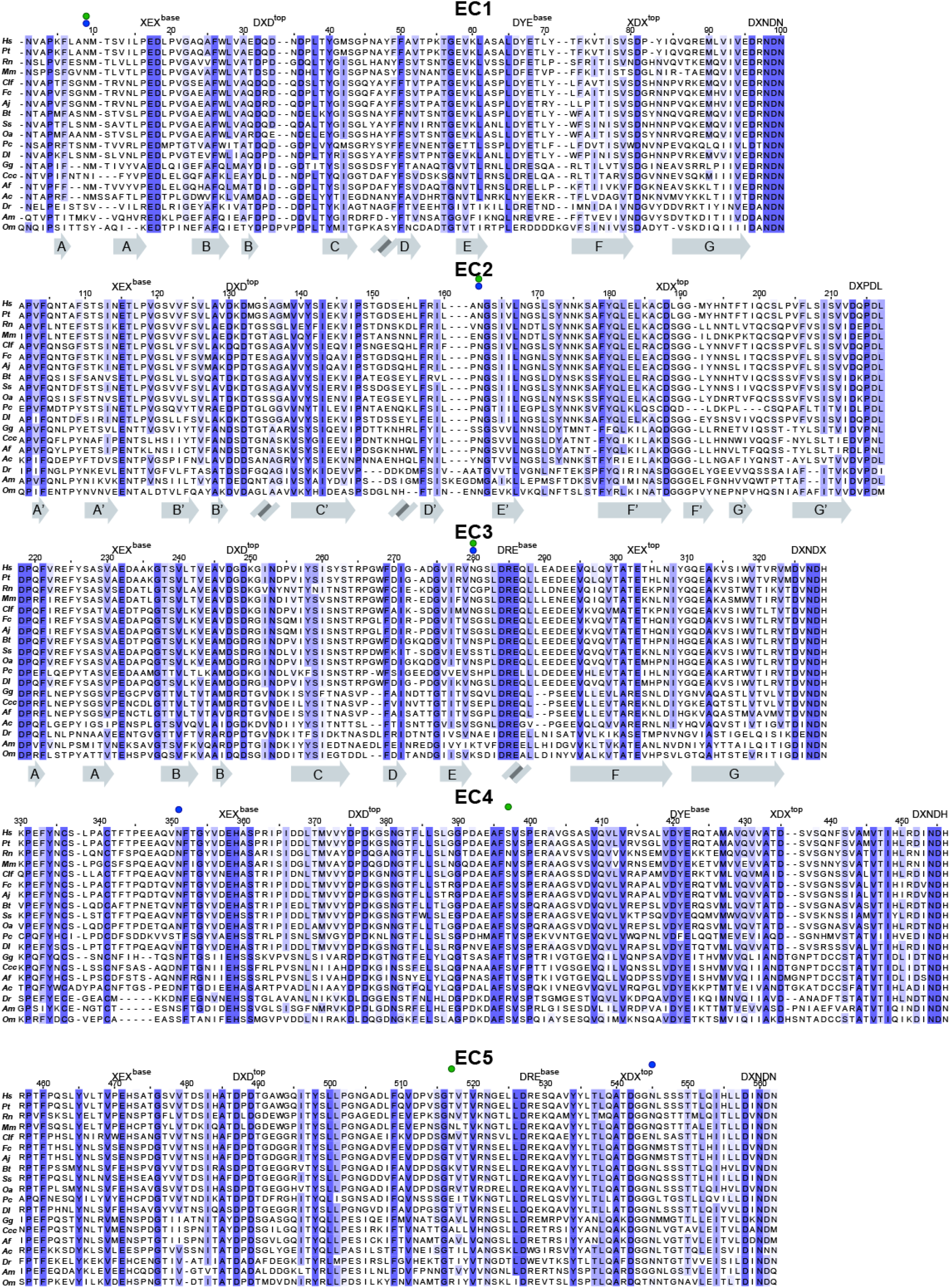
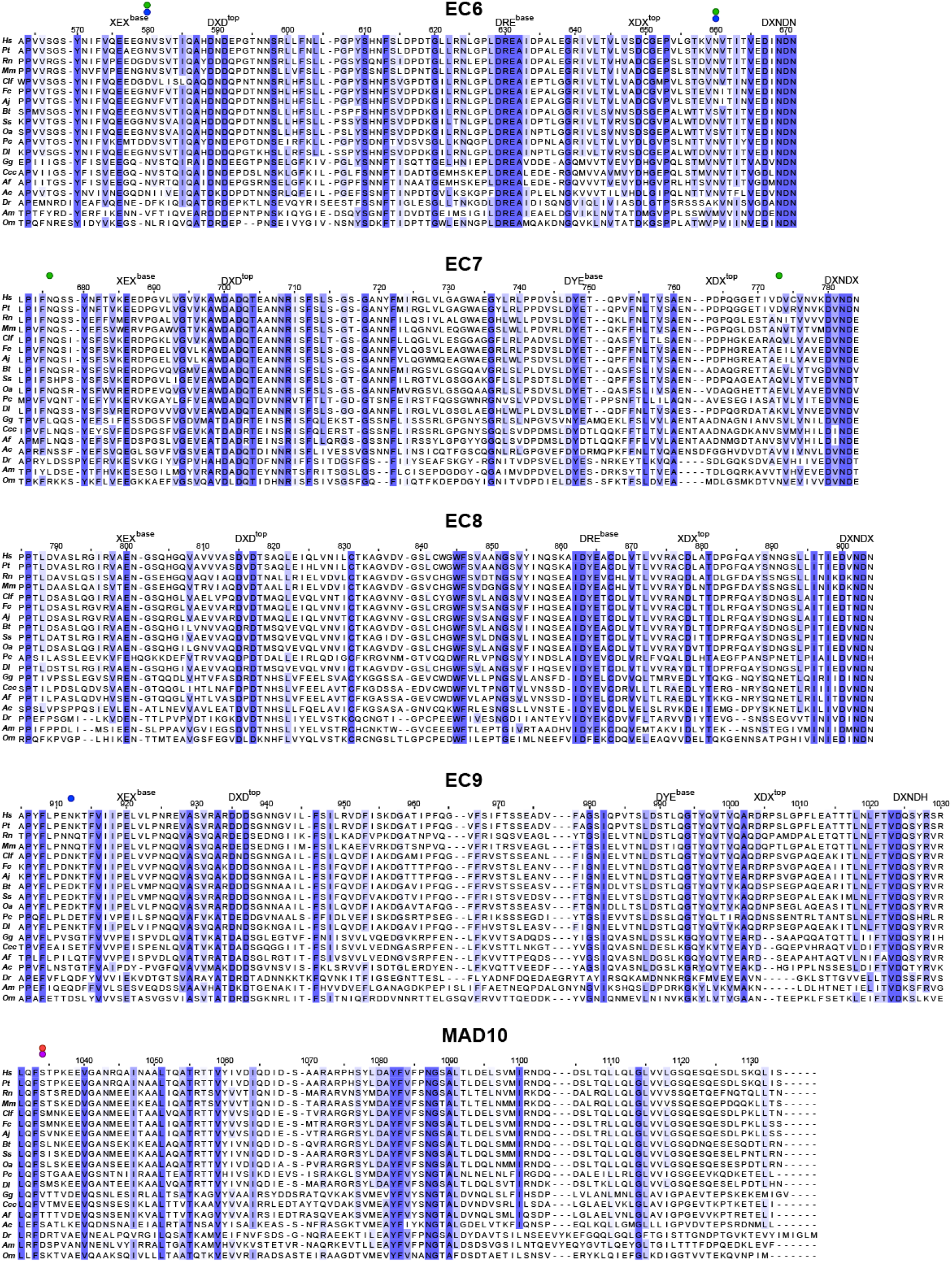
Sequence alignments of individual EC repeats of PCDH24. Multiple sequence alignments comparing each EC repeat and MAD10 of PCDH24 from 19 different species. Each alignment is colored by percent identity, with white being the lowest percent identity and dark blue being the highest. Sites of N-linked (blue in human, green in mouse) and O-linked (purple in human, red in mouse) glycosylation are denoted by a colored circle. Secondary structure elements observed in the crystal structures of *hs* PCDH24 EC1-2 and *mm* PCDH24 EC1-3 are illustrated below the respective repeats. Calcium-binding motifs are indicated above the sequences, which are numbered according to the human protein. Species are abbreviated as follows: *Homo sapiens (hs), Pan troglodytes (Pt), Rattus norvegicus (Rn), Mus musculus (mm), Canis lupus familiaris (Clf), Felis catus (Fc), Acinoyx jubatus (Aj), Bos Taurus (Bt), Sus scrofa (Ss), Ovis aries (Oa), Phascolarctos cinereus (Pc), Delphinapterus leucas (Dl), Gallus gallus (Gg), Corvus cornix cornix (Ccc), Aptenodytes forsteri (Af), Anolis carolinensis (Ac), Danio rerio (Dr), Astyanax mexicanus (Am)*, and *Oncorhynchus mykiss (Om)*. Species were chosen based on sequence availability and taxonomical diversity. Accession numbers and species can be found in Table S2.

**Figure S2.**
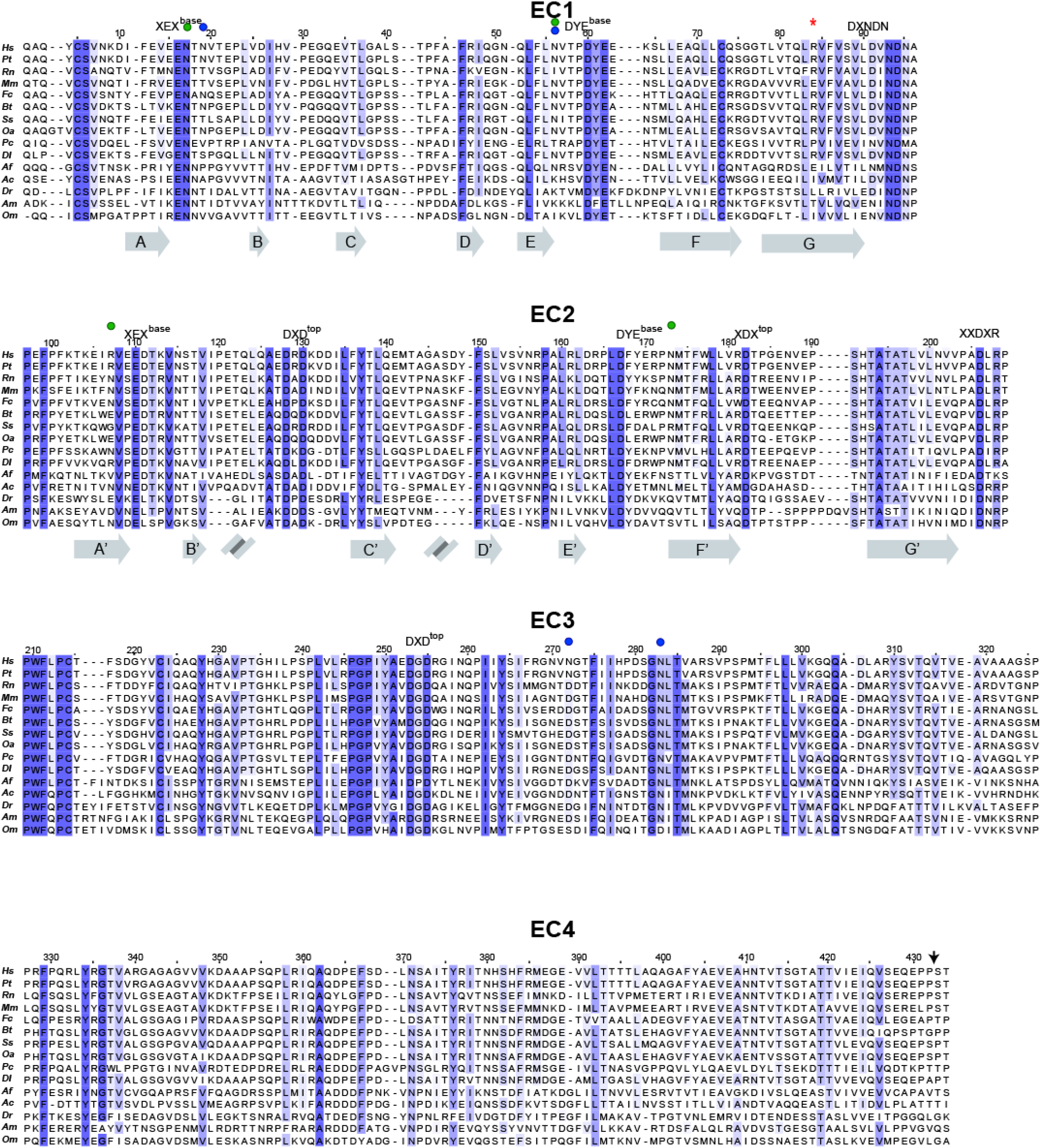
Sequence alignments of individual EC repeats of CDHR5. Multiple sequence alignments comparing each EC repeat of CDHR5 from 15 different species, shown as in Figure S1. An asterisk (*) indicates site R84 mutated in binding assays. An arrow indicates the end of EC1-EC4 protein fragments used in binding assays. Secondary structure elements observed in the crystal structures of *hs* CDHR5 EC1-2 are illustrated below the respective repeats. Calcium-binding motifs are indicated above the sequences, which are numbered according to the human protein. Species are abbreviated as follows: *Homo sapiens (hs), Pan troglodytes (Pt), Rattus norvegicus (Rn), Mus musculus (mm), Felis catus (Fc), Bos Taurus (Bt), Sus scrofa (Ss), Ovis aries (Oa), Phascolarctos cinereus (Pc), Delphinapterus leucas (Dl), Aptenodytes forsteri (Af), Anolis carolinensis (Ac), Danio rerio (Dr), Astyanax mexicanus (Am)*, and *Oncorhynchus mykiss (Om)*. Species were chosen based on sequence availability and taxonomical diversity. Accession numbers and species can be found in Table S3.

**Figure S3.**
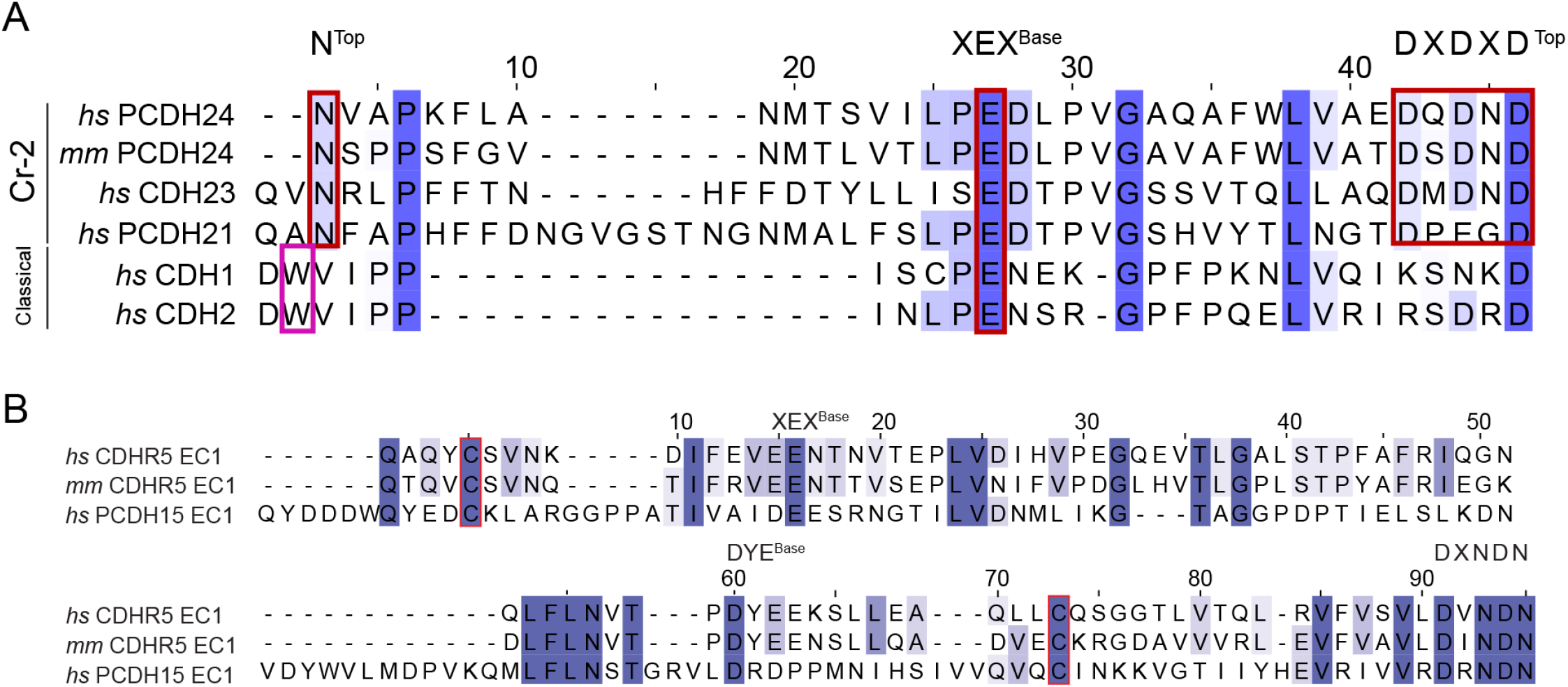
Comparison of PCDH24 and CDHR5 N-termini with other cadherins. (A) Alignment of the processed N-terminal sequences of Cr-2 protocadherins *hs* PCDH24, *mm* PCDH24, *hs* CDH23 and *hs* PCDH21, and of two classical cadherins, *hs* CDH1 and *hs* CDH2. Calcium-binding sites are labeled and boxed in red in the sequence to show the additional calcium-bindings sites in Cr-2 protocadherins not present in classical cadherins. Residue W2 involved in classical cadherin binding is boxed in pink. (B) Alignments of the processed sequences of EC1 for *hs* CDHR5, *mm* CDHR5, and *hs* PCDH15. Calcium-binding sites are labeled. The two cysteine residues that form a disulfide bond between β-strands A and F are highlighted by a red box.

**Figure S4.**
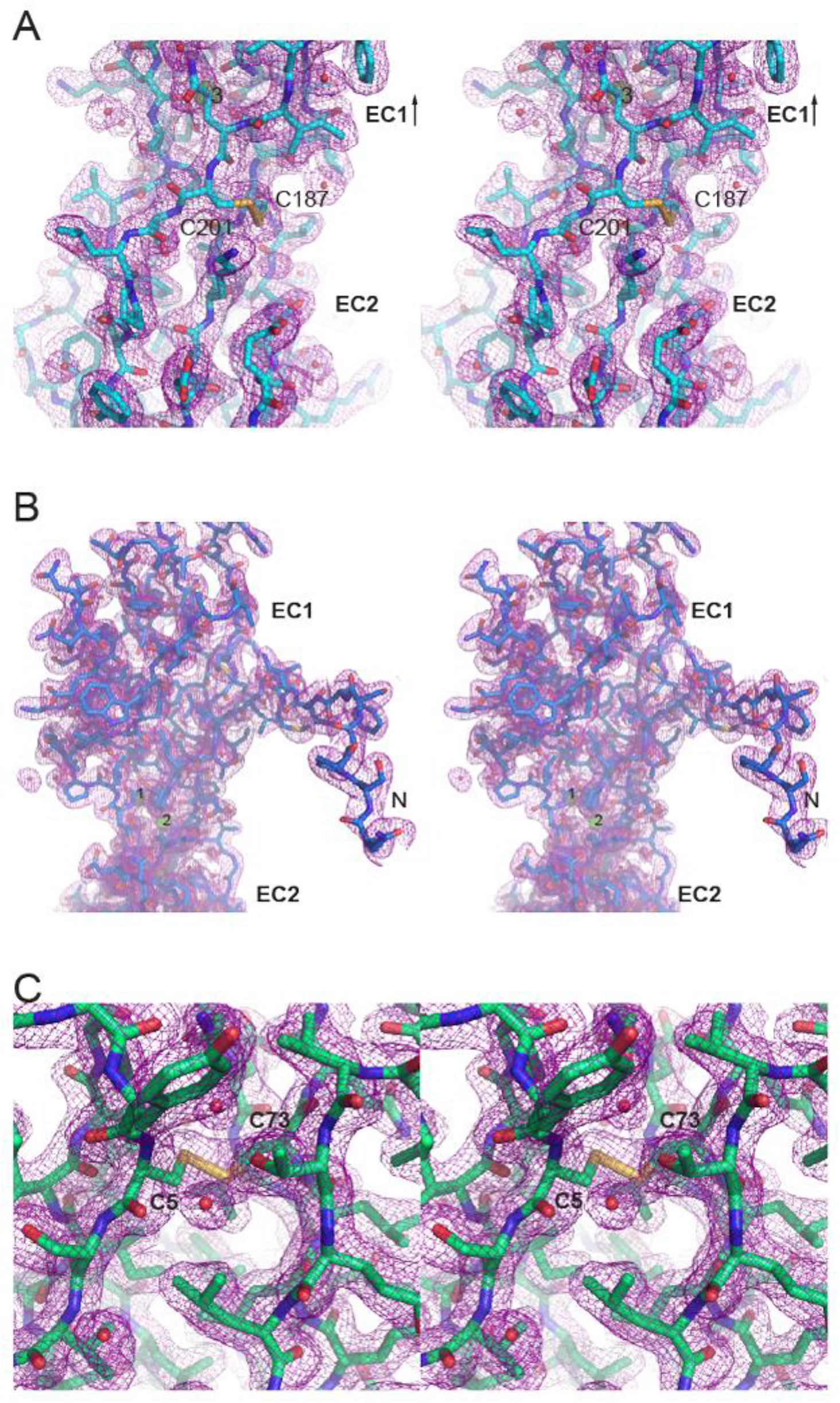
Electron density for unique elements of *hs* PCDH24 EC1-2, *mm* PCDH24 EC1-3 and *hs* CDHR5 EC1-2. (A) Extended F-G loop of EC2 in *hs* PCDH24 where the disulfide bond is located (C187 - C201) shown at 1.0 σ using carve of 1.6. This loop is also present in the *mm* PCDH24 structure. (B) The atypical extended N-terminus seen in *mm* PCDH24 EC1 shown at 1.0 σ and carve of 1.8. (C) Disulfide bond between residue C5 in β-strand A and residue C73 in β-strand F of *hs* CDHR5 EC1-2 shown at 1.0 σ and carve of 1.6.

**Figure S5.**
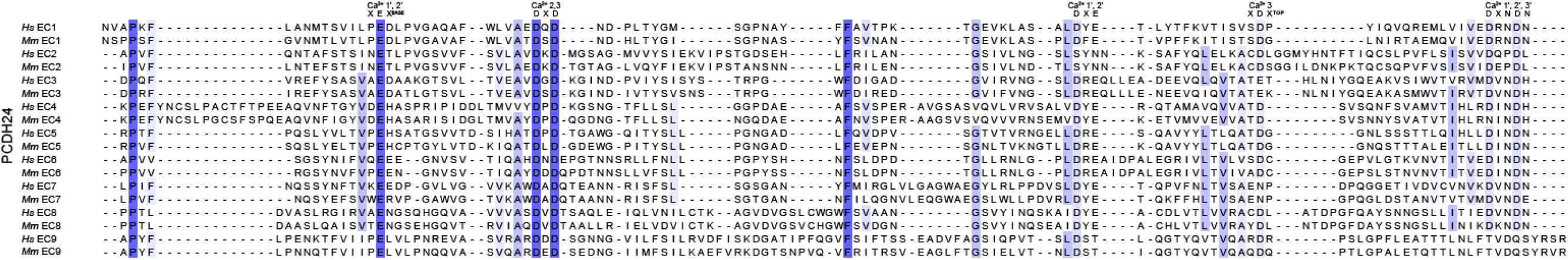
Sequence alignment of *hs* and *mm* PCDH24 EC repeats. All 9 EC repeats for each species are aligned to each other (EC1-9). Conserved calcium-binding motifs are labeled on top of the alignment.

**Figure S6.**
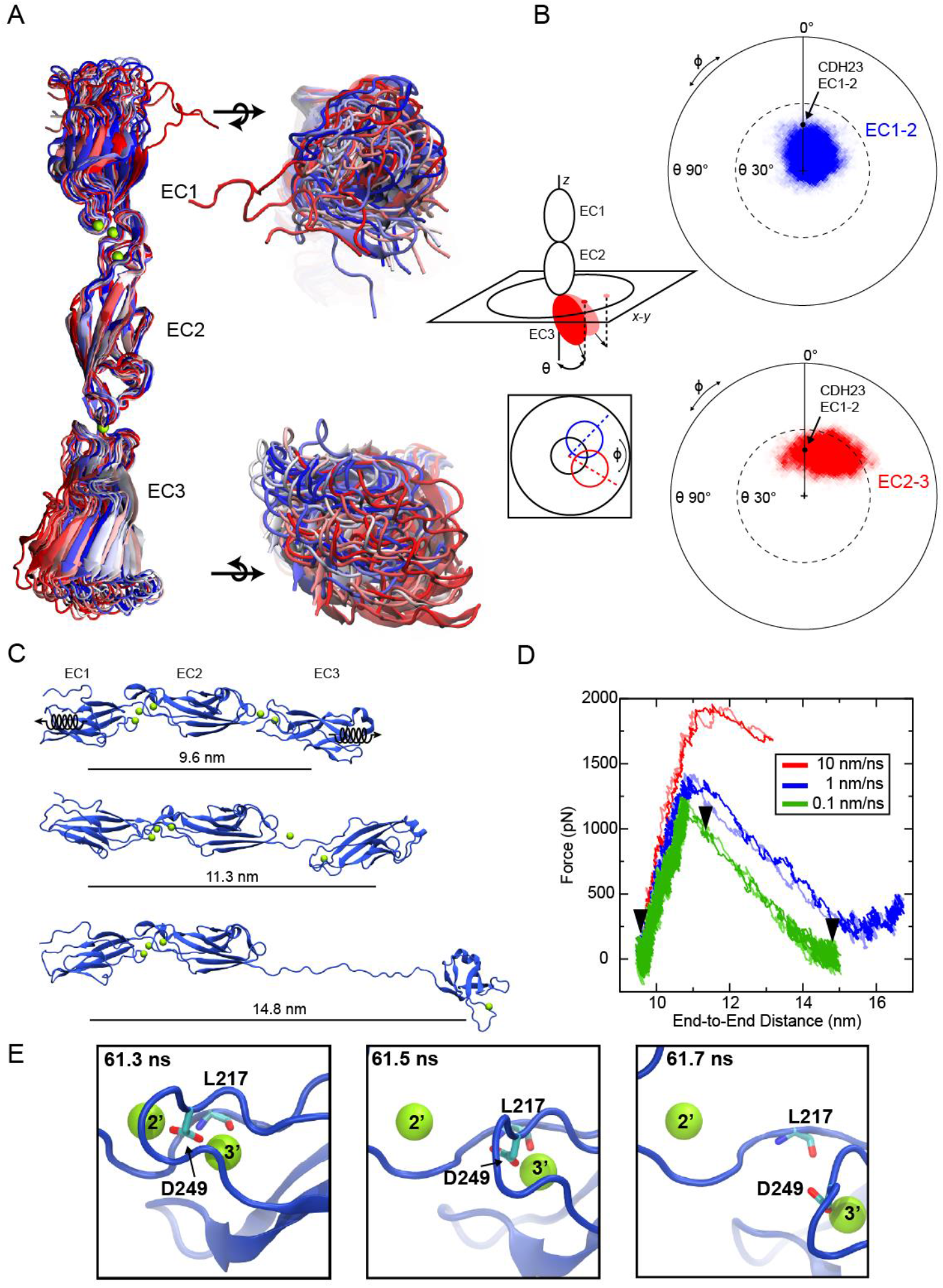
Equilibrium and stretching simulations of *mm* PCDH24 EC1-3. (A) Superposition of *mm* PCDH24 EC1-3 conformations taken every 5 ns from a ∼99 ns long trajectory (simulation S2, Table S10). Repeat EC2 was used as a reference. Side, top, and bottom views are shown. Color indicates time step (red-white-blue). (B) Orientation projections illustrating the conformational freedom of the EC1-2 and EC2-3 linkers throughout equilibrium simulations. To quantify the conformational freedom of EC2 relative to EC1 (top), and of EC3 relative to EC2 (bottom), the longest principal axes of EC1 and EC3 were aligned to the *z* axis and then the projections of the longest principal axes of EC2 (blue) and EC3 (red) in the *x-y* plane were plotted. The initial orientation of a control protomer, CDH23 EC1-2 (2WHV), is shown as a black dot (Sotomayor et al., 2010). The EC2-3 linker behavior is not dramatically different than the behavior observed for the EC1-2 linker, suggesting similar flexibility. (C) Trajectory snapshots during the slowest speed stretching simulation at 0.1 nm/ns for *mm* PCDH24 EC1-3 (S3d, Table S10). Springs indicate position (center of mass) and direction of applied forces. Unfolding of the EC2-3 linker is observed first. End-to-end distances between the centers of mass of EC1 and EC3 are indicated for each snapshot. (D) Force versus end-to-end distance for simulations of the *mm* PCDH24 EC1-3 monomer (S3a-d) at stretching speeds of 10 nm/ns (red), 1 nm/ns (blue) and 0.1 nm/ns (green). Dark and light colors indicate forces applied at opposite ends. Black arrowheads indicate time points illustrated in (C). (E) Detail of the *mm* PCDH24 EC2-3 linker at the force peak during the slowest stretching simulation at 0.1 nm/ns (S3d, Table S10). Residues involved in two-step unbinding from calcium ions are shown.

**Figure S7.**
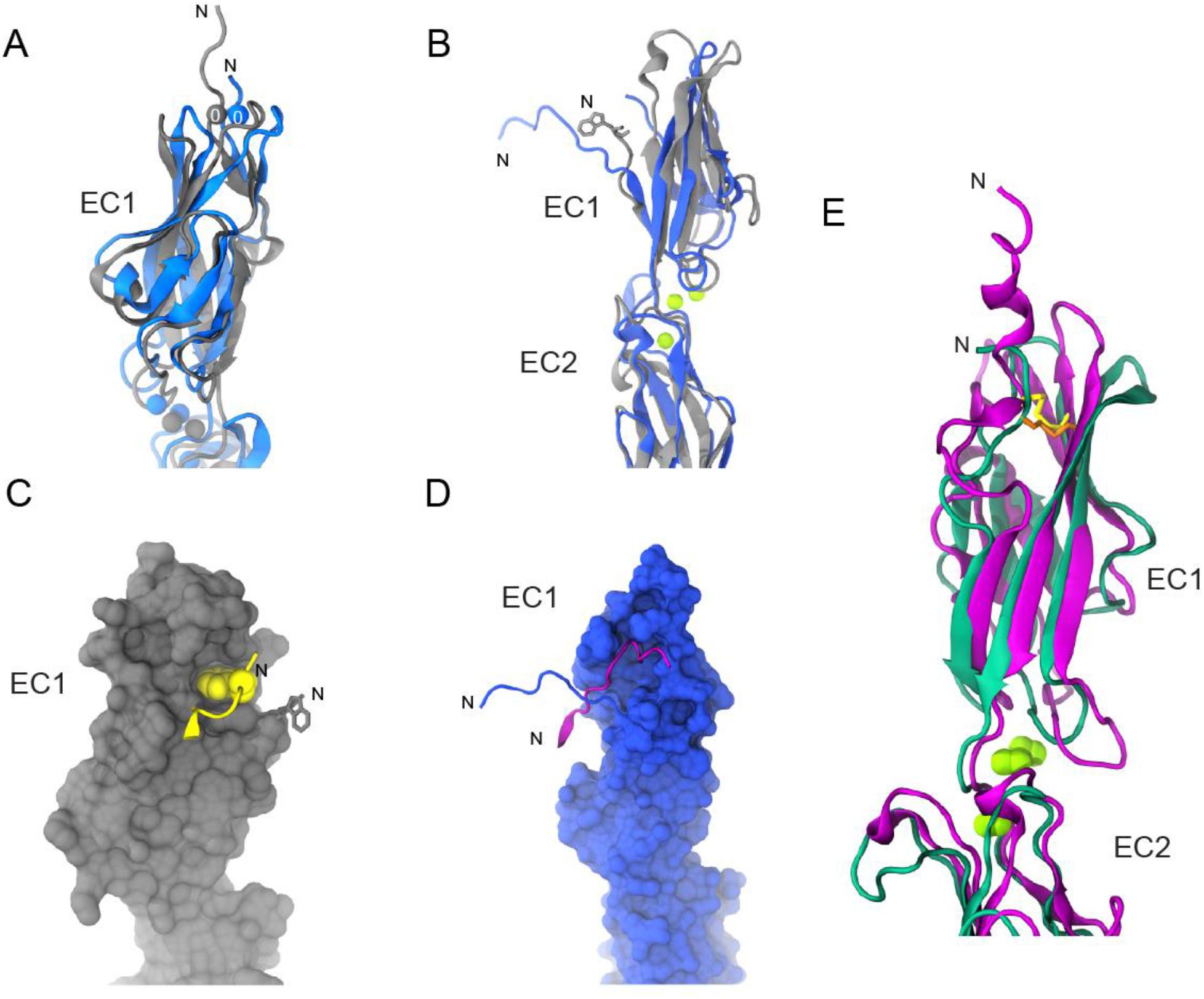
Structural comparison of PCDH24 and CDHR5 EC1 repeats with other cadherins. (A) Detail of superposed EC1 N-termini of *hs* PCDH24 (blue) and *mm* CDH23 (grey; PDB: 2WHV). Both have a calcium ion at the calcium-binding site 0, a feature of Cr-2 protocadherins. (B) Detail of superposed EC1 N-termini of *mm* PCDH24 (blue) and *hs* CDH1 (grey; PDB: 2O72). Both have N-termini protruding away from the protomer to interact with another neighboring EC1, as shown in (C) for *hs* CDH1 and (D) for *mm* PCDH24. (C-D) Surface representations of *hs* CDH1 and *mm* PCDH24, respectively. Highlighted in yellow is β-strand A with W2 from its binding partner. Residues 25, 26, and 27 are not shown to facilitate visualization of the tryptophan binding pocket. Highlighted in purple is the N-terminus of *mm* PCDH24 from a neighboring protomer in the crystal lattice (Figure 2D). (E) Detail of *hs* CDHR5 EC1 (green) and *mm* PCDH15 EC1 (purple) superposed. Disulfide bonds are shown in orange and yellow. PCDH15 has longer loops and a longer N-terminus, with a helix that interacts with CDH23.

**Figure S8.**
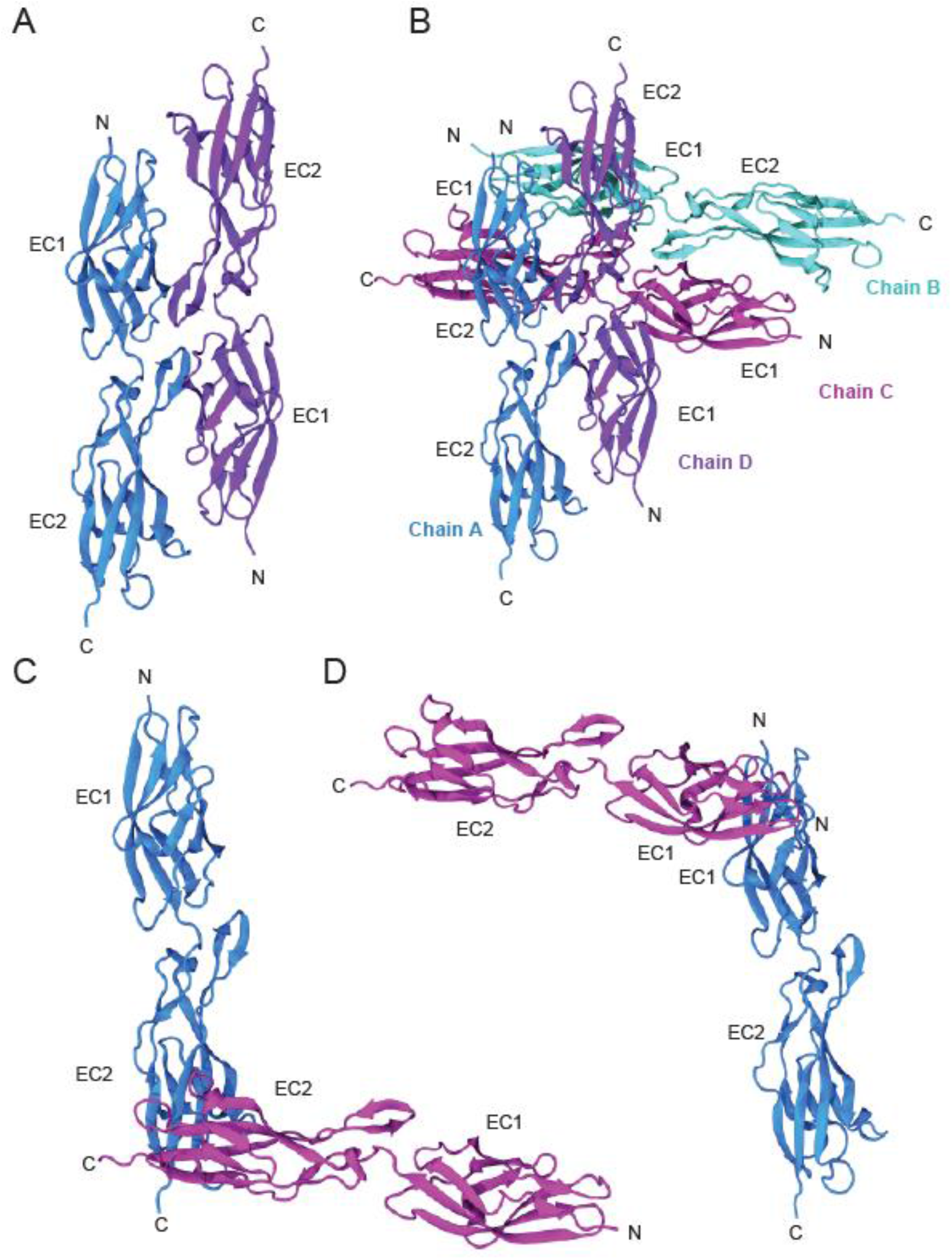
Crystal contacts in the *hs* PCDH24 EC1-2 structure. (A-D) Ribbon diagram of two protomers showing a crystal contact between chains A and D with an interface area of 979.2 Å^2^. The arrangement is antiparallel, likely describing a possible *trans* interface (A). Crystal contacts in the entire asymmetric unit include four additional interfaces with areas of 173.6 Å^2^ (chains A and B), 231.1 Å^2^ (chains A and C), 255.3 Å^2^ (chains D and B), and 307.2 Å^2^ (chains D and C). Two Additional interfaces between chains A and C (387.4 Å^2^ and 507.3 Å^2^) are shown in (C) and (D). Equivalent interfaces for the one shown in (A) between chains B and C, and those shown in (C, D) between chains D and B, have similar interface areas of 938.6 Å^2^, 390.8 Å^2^, and 515.1 Å^2^, respectively, but are not shown.

**Figure S9.**
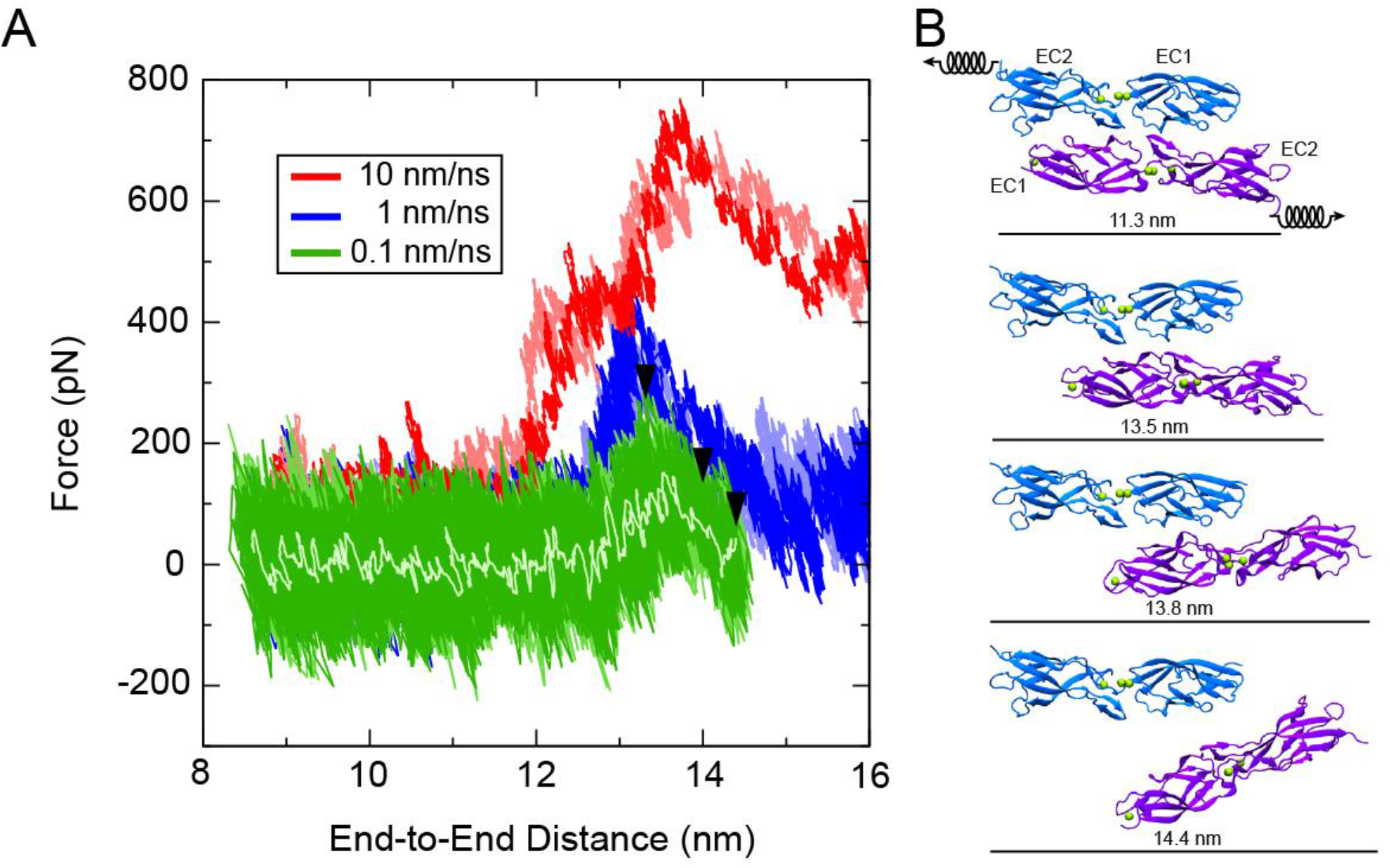
Stretching simulations testing the strength of the *hs* PCDH24 EC1-2 *trans* interface. (A) Force versus end-to-end distance for simulations of the largest crystallographic *trans* interface (Figure S8A) observed for *hs* PCDH24 EC1-2 (simulations S1b-S1d, Table S10). Forced unbinding simulations were carried out at stretching speeds of 10 nm/ns (red), 1 nm/ns (blue), and 0.1 nm/ns (green, 50 ps running average in light green). (B) Snapshots of unbinding trajectory during stretching simulation at 0.1 nm/ns (S1d, Table S10). Springs indicate position and direction of applied forces. Top panel shows complex at the beginning of the simulation, other panels show snapshots at three time points indicated with black arrow heads in (A). End-to-end distance indicates the distance between the stretched atoms on the C-termini of EC2.

**Figure S10.**
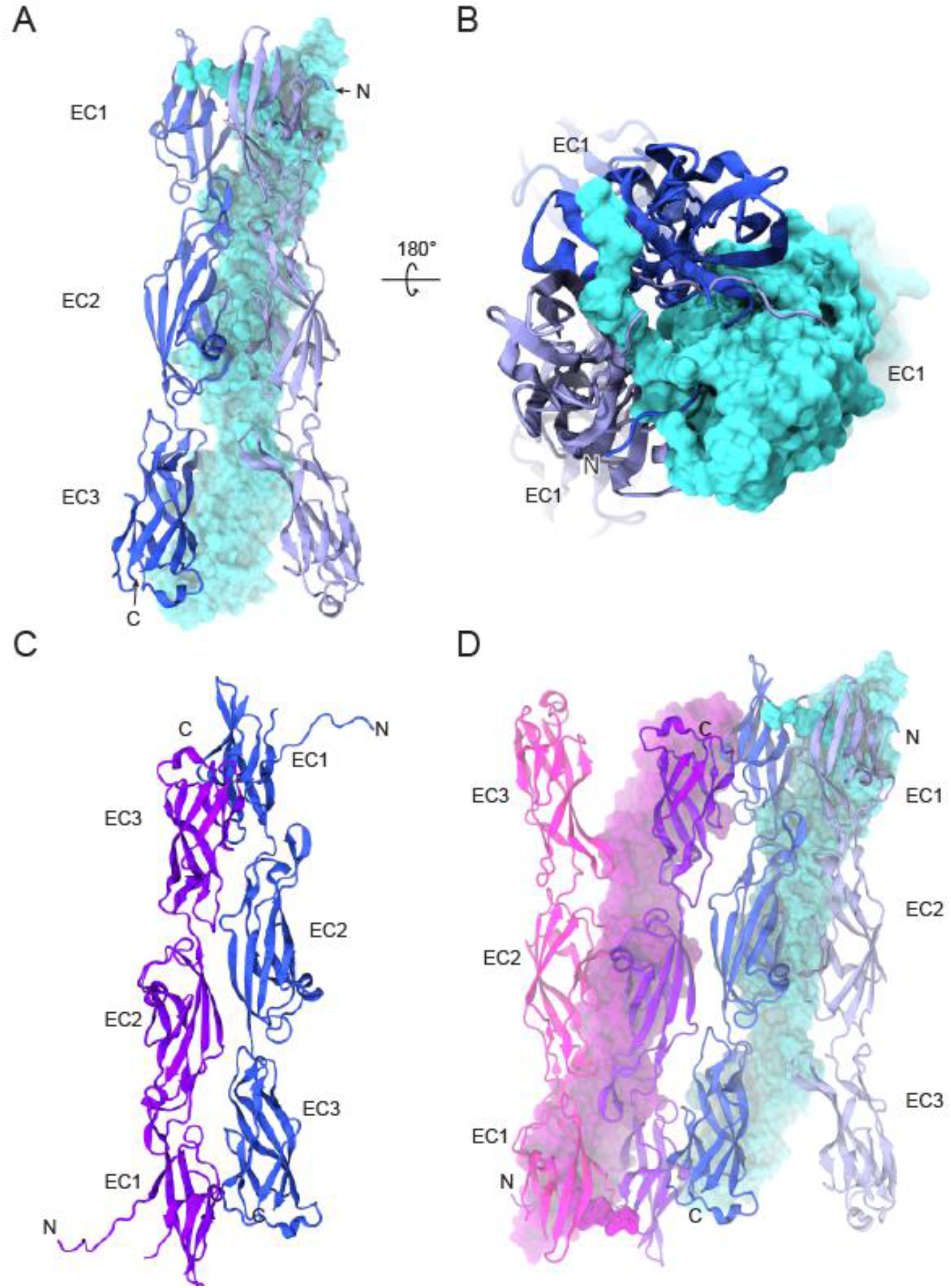
Crystal contacts in the *mm* PCDH24 EC1-3 structure. (A) Crystal contacts show a potential *cis* trimer formed by three parallel monomers. The interface area between two monomers is 1352.6 Å^2^. Two monomers are shown as ribbons while the third one is shown in molecular surface representation. (B) Detail of trimer seen from EC1 (top) shows that the extended N-terminal strands interlace between monomers. (C) Crystal contacts show an antiparallel *trans* dimer with an interface area of 1221.2 Å^2^. (D) Taken together, the potential *cis* and *trans* interfaces form a large complex with two *cis* trimers forming an antiparallel *trans* dimer.

**Figure S11.**
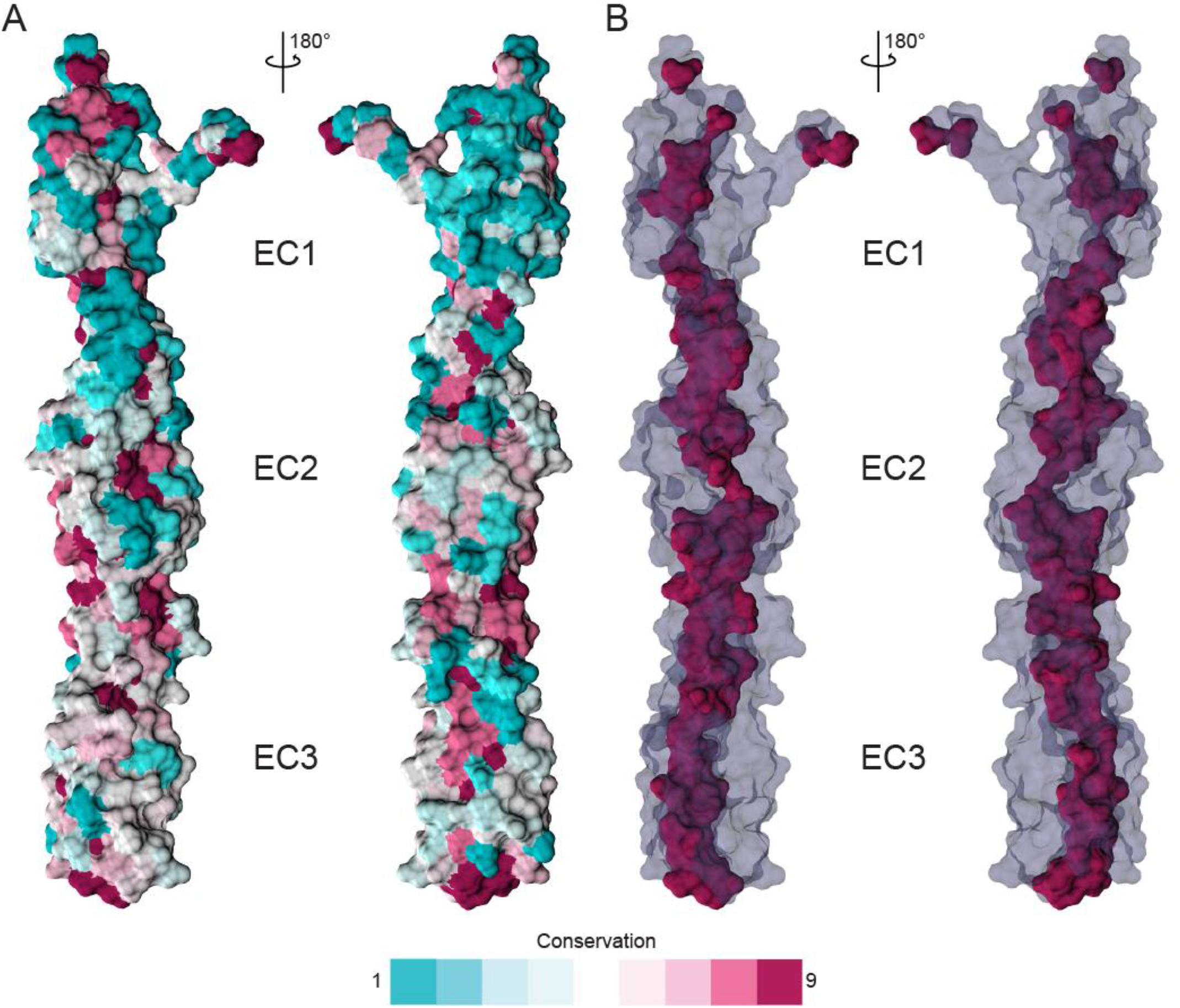
Sequence conservation of PCDH24 EC1-3. (A) Surface representation of *mm* PCDH24 EC1-3 structure with residues colored according to sequence conservation determined using Consurf and a sequence alignment including over 94 species. Teal colors indicate residues that are least conserved while magenta indicates residues that are most conserved among species. (B) Transparent surface representation of *mm* PCDH24 EC1-3 with most conserved residues shown as an opaque magenta surface. Protein core is conserved.

**Figure S12.**
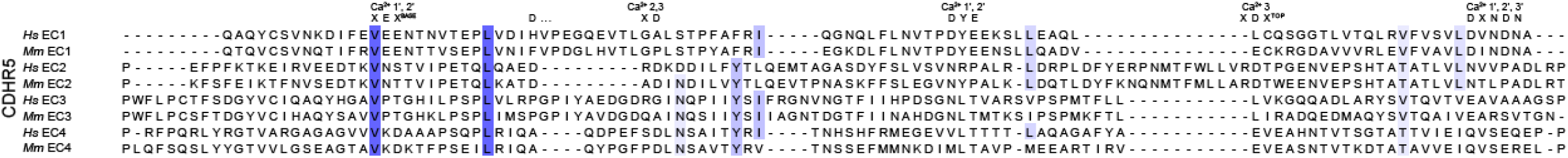
Sequence alignment of *hs* and *mm* CDHR5 EC repeats. All 4 EC repeats for each species are aligned to each other (EC1-4).

**Figure S13.**
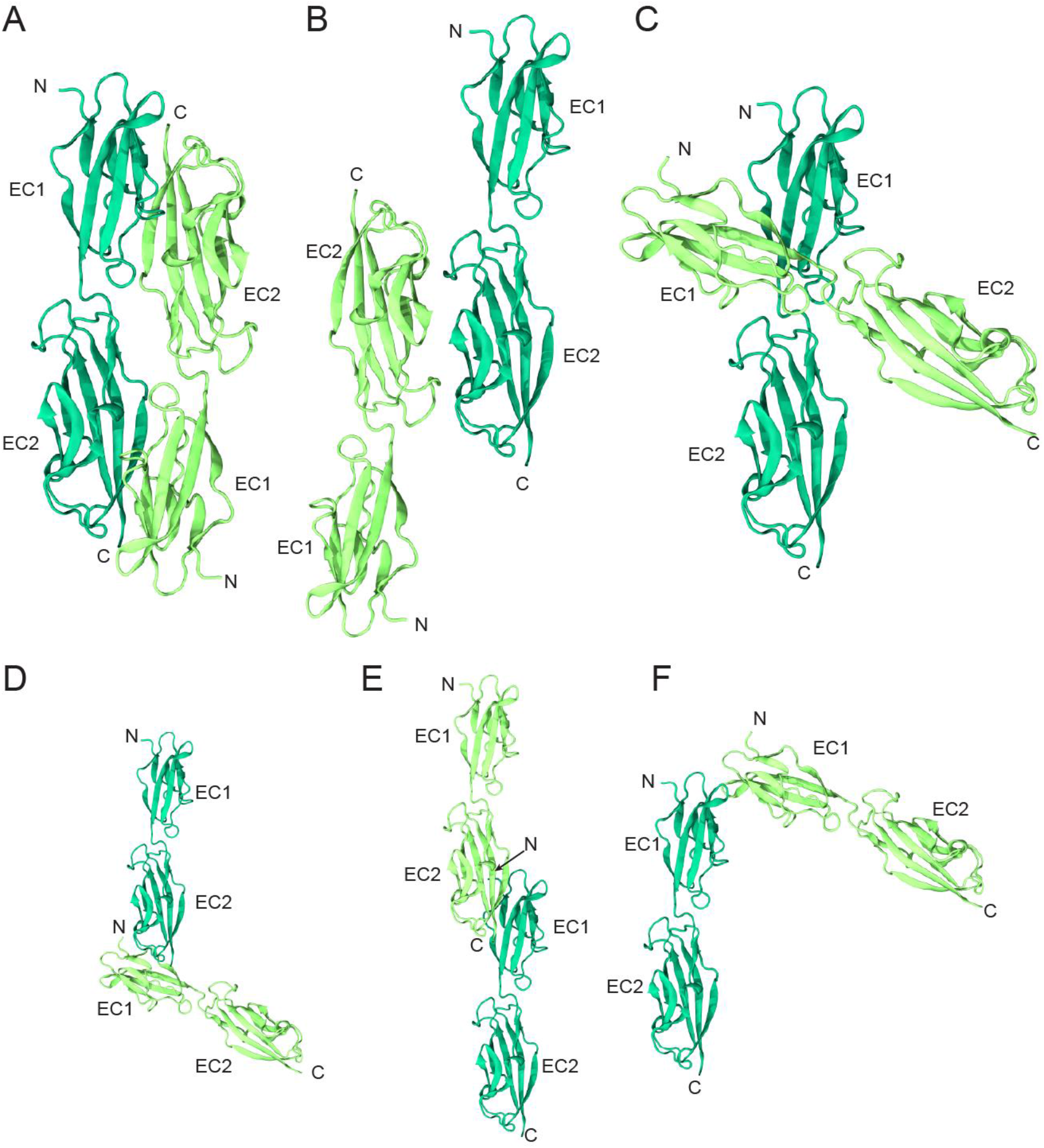
Crystal contacts in the *hs* CDHR5 EC1-2 structure. (A-F) Ribbon diagram of two protomers showing various crystal contacts between monomers of *hs* CDHR5 EC1-2. Interface areas are 1433.9 Å^2^ (A), 530.2 Å^2^ (B), 364.7 Å^2^ (C), 347.9 Å^2^ (D), 132.7 Å^2^ (E), and 114.0 Å^2^ (F), respectively. Interfaces in (A) and (B) correspond to possible *trans* overlaps of EC1-2 and EC1-3, respectively. While *hs* CDHR5 does not mediate *trans* homophilic adhesion based on the homophilic binding assay data presented in Figure 4, these interfaces may serve as templates for the mouse CDHR5 protein. Crystal contacts in (D-F) are unlikely to be of physiological relevance due to the arrangement of the monomers.

**Figure S14.**
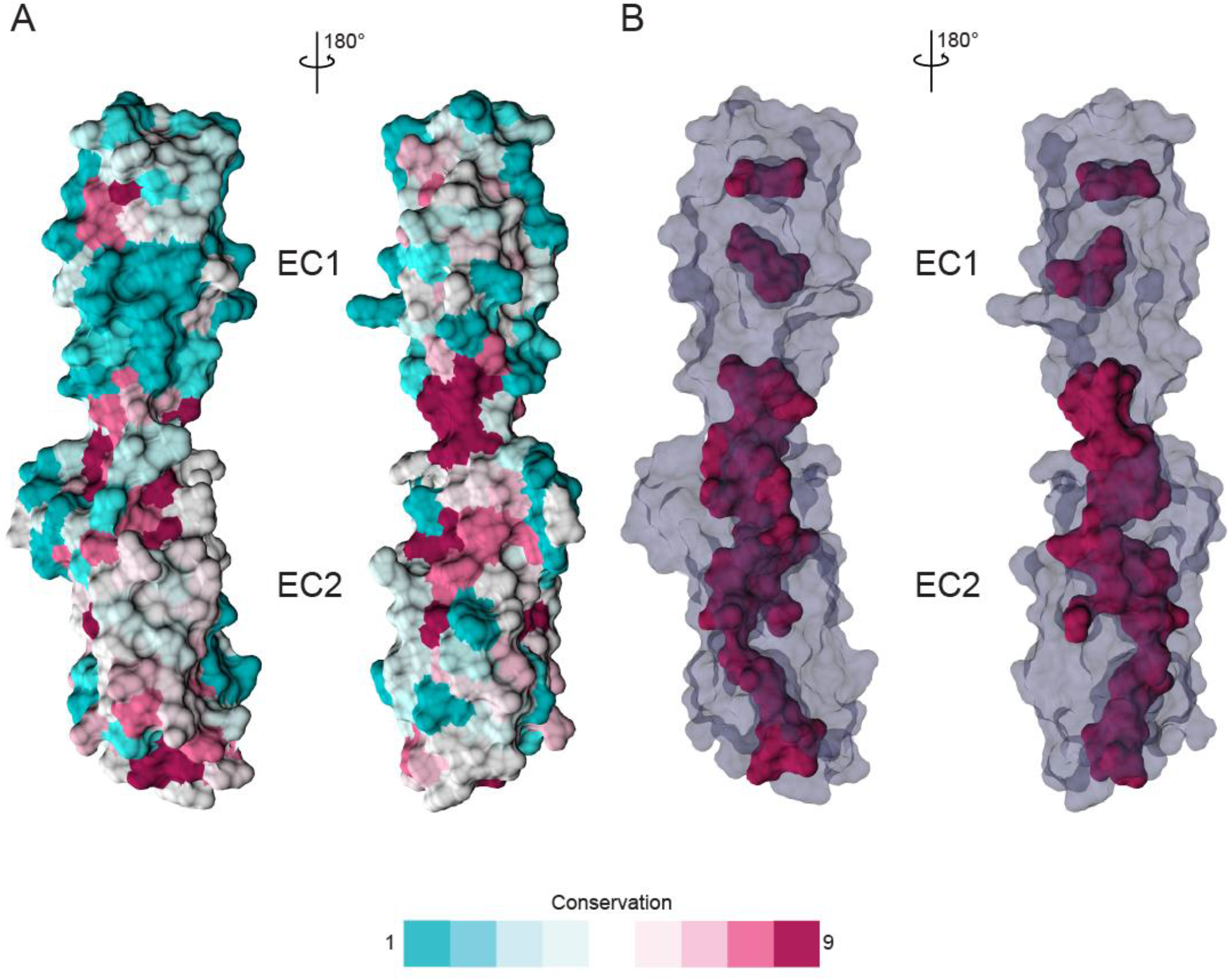
Sequence conservation of CDHR5 EC1-2. (A) Surface representation of *hs* CDHR5 EC1-2 structure with residues colored according to sequence conservation determined using Consurf and a sequence alignment including over 69 species. Teal colors indicate residues that are least conserved while magenta indicates residues that are most conserved among species. (B) Transparent surface representation of *hs* CDHR5 EC1-2 with most conserved residues shown as an opaque magenta surface.

**Figure S15.**
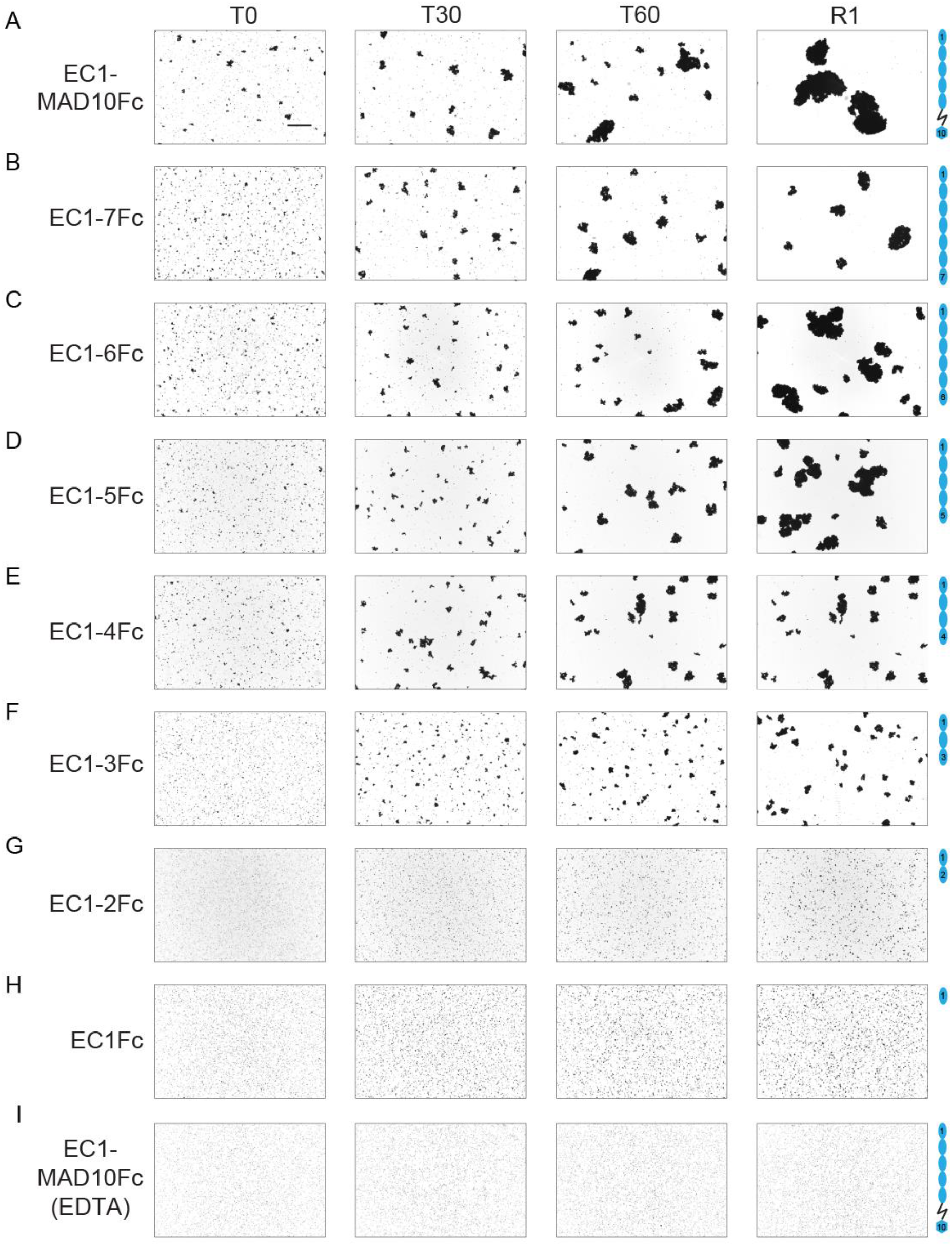
Homophilic binding assays of *hs* PCDH24 at various time points. (A-H) Protein G beads coated with the full-length extracellular domain of *hs* PCDH24 (A) and its C-terminal truncation versions (B-H). Images show bead aggregation observed at the start of the experiment (T0), after 30 min (T30), after 60 min (T60) followed by rocking for 1 min (R1), all in the presence of 2 mM CaCl_2_. Bar – 500 µm. (I) Protein G beads coated with the full-length extracellular domain of *hs* PCDH24 in the presence of 2 mM EDTA, shown as in (A).

**Figure S16.**
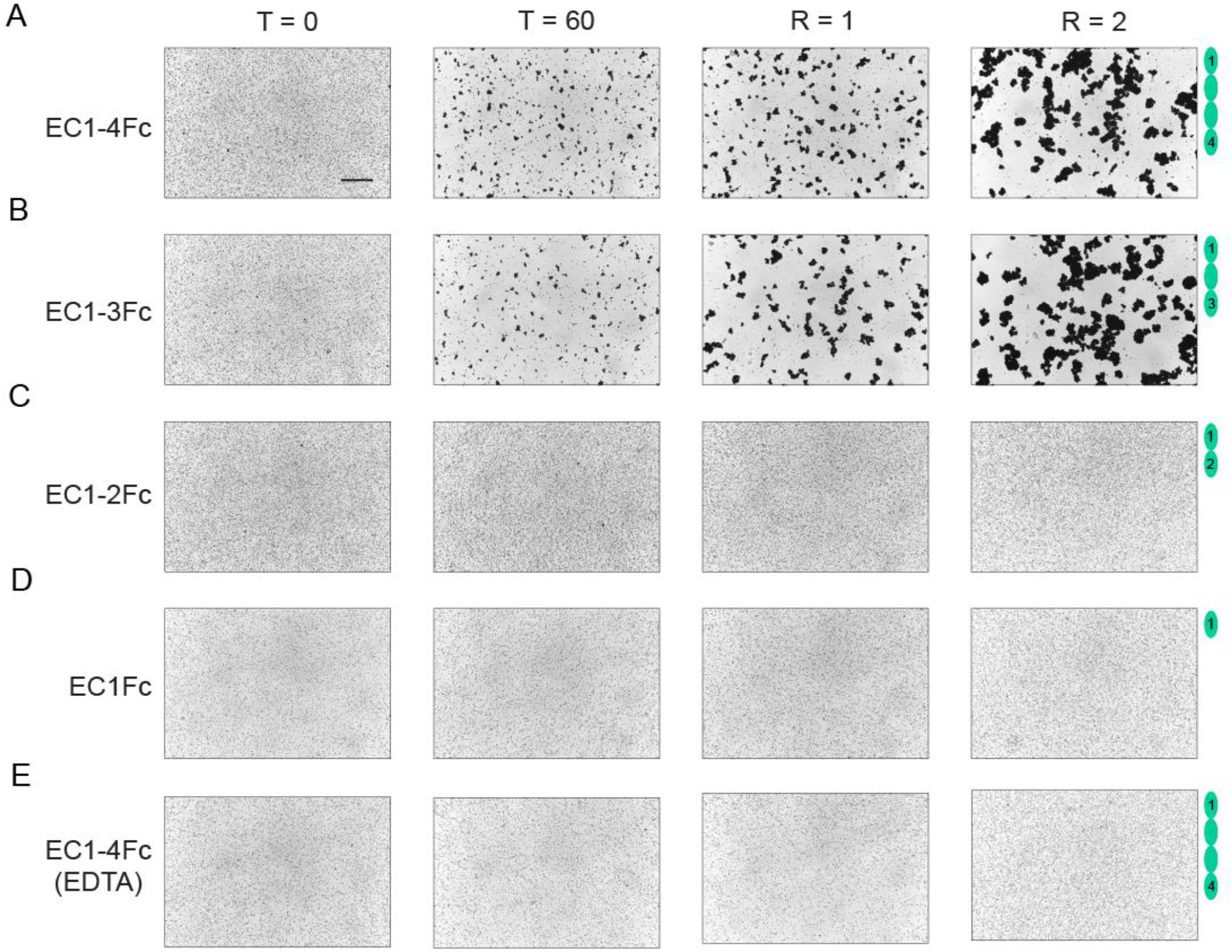
Homophilic binding assays of *mm* CDHR5 at various time points. (A-D) Protein G beads coated with the full-length cadherin extracellular domain of *mm* CDHR5 (A) and its C-terminal truncation versions (B-D). Images show the aggregation observed at the start of experiment (T0), after 60 min (T60) followed by rocking for 1 min (R1) and 2 min (R2) in the presence of 2 mM CaCl_2_. Bar – 500 µm. (E) Protein G beads coated with the full-length cadherin extracellular domain of *mm* CDHR5 in the presence of 2 mM EDTA shown as in (A).

**Figure S17.**
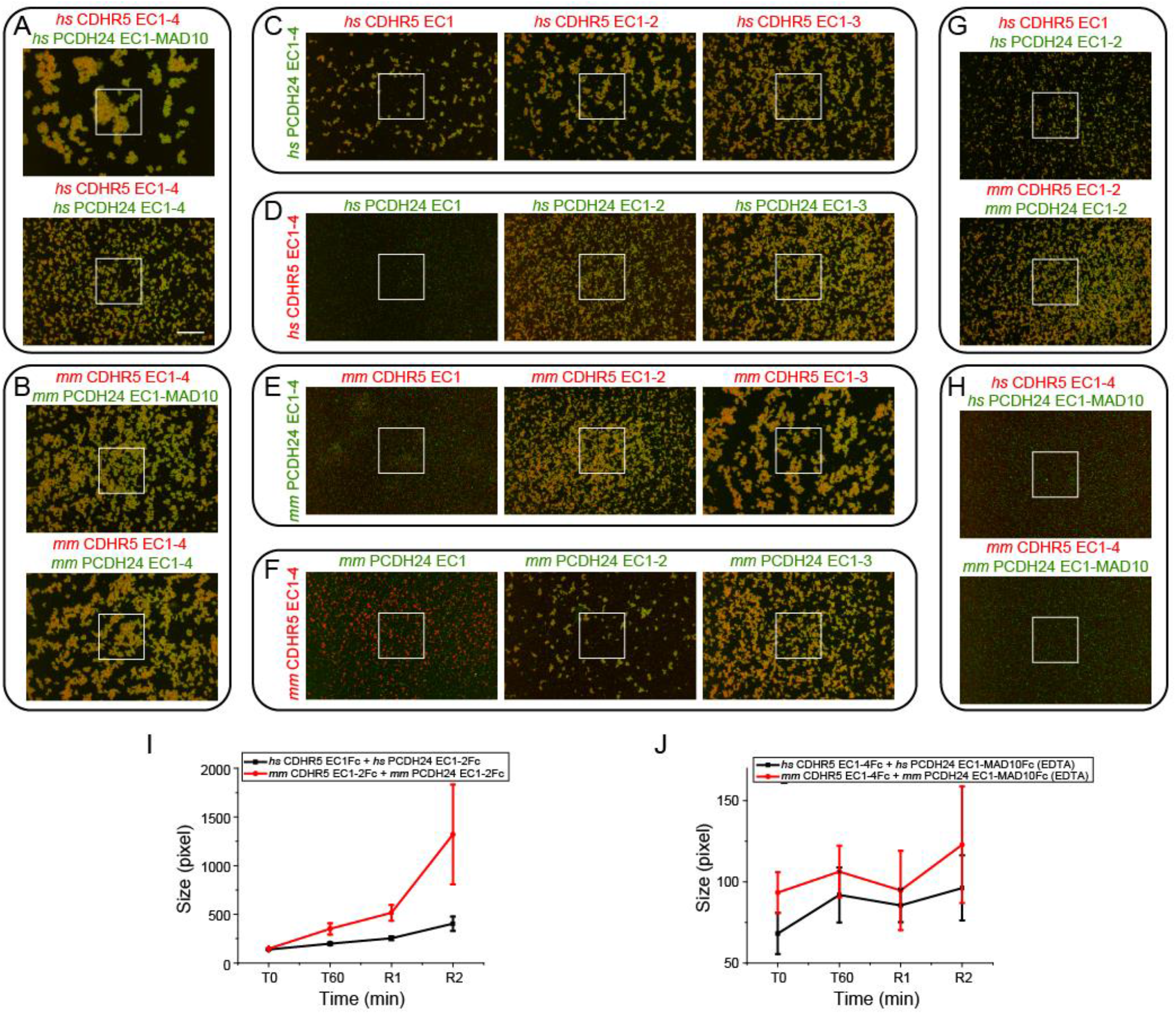
Heterophilic binding assays of *hs* and *mm* PCDH24 and CDHR5. (A-B) Images from binding assays of *hs* PCDH24 (full-length extracellular domain and EC1-4Fc) mixed with the full-length *hs* CDHR5 cadherin extracellular domain (A), and *mm* PCDH24 (full-length extracellular domain and EC1-4Fc) mixed with full-length *mm* CDHR5 cadherin extracellular domain (B). Green fluorescent protein A beads are coated with PCDH24 fragments and red fluorescent protein A beads are coated with CDHR5 in all panels. The white boxes indicate the regions shown in Figure 6. Images show bead aggregation observed after 60 min followed by rocking for 2 min in the presence of 2 mM CaCl_2_. (C-F) Images from binding assays of truncations of *hs* CDHR5 mixed with *hs* PCDH24 EC1-4Fc (C), truncations of *hs* PCDH24 mixed with *hs* CDHR5 EC1-4Fc (D), truncations of *mm* CDHR5 mixed with *mm* PCDH24 EC1-4Fc (E), and truncations of *mm* PCDH24 mixed with *mm* CDHR5 EC1-4Fc (F). Images show bead aggregation observed after 60 min followed by rocking for 2 min in the presence of 2 mM CaCl_2_. (G) Images from binding assays of minimum EC repeats required for heterophilic adhesion of *hs* and *mm* PCDH24 and CDHR5. The minimum units for heterophilic adhesion for the human proteins are CDHR5 EC1Fc and PCDH24 EC1-2Fc. The minimum units for heterophilic adhesion for the mouse proteins are CDHR5 EC1-2Fc and PCDH24 EC1-2Fc. Images show bead aggregation observed after 60 minutes followed by rocking for 2 minutes in the presence of 2 mM CaCl_2_. (H) Protein A beads coated with full-length *hs* and *mm* PCDH24 and CDHR5 cadherin extracellular domains (including MAD10 for PCDH24 when indicated and without the mucin-like domain for CDHR5) in the presence of 2 mM EDTA shown as in (A-B). Bar – 500 µm. (I) Aggregate size for minimum units of heterophilic adhesion of *hs* and *mm* PCDH24 and CDHR5 at the start of the experiment (T0), after 60 min (T60) followed by rocking for 1 min (R1) and 2 min (R2). (J) Aggregate size for full-length *hs* and *mm* PCDH24 and CDHR5 at the start of the experiment (T0), after 60 min (T60) followed by rocking for 1 min (R1) and 2 min (R2). Error bars in I and J are standard error of the mean (*n* indicated in Table S7).

**Table S1.**
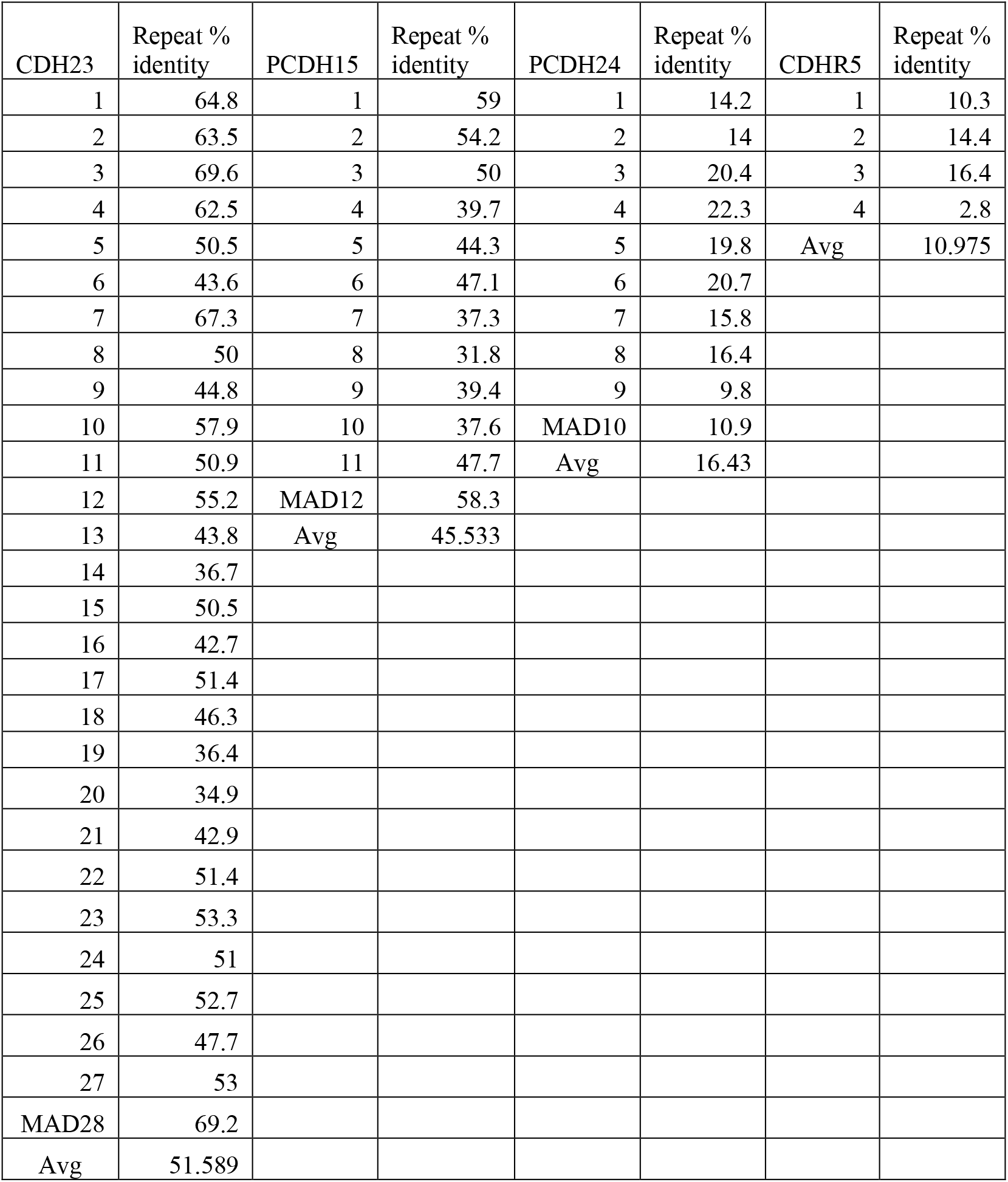
Percent Identity of EC repeats of CDH23, PCDH15, PCDH24 and CDHR5 proteins across species. The accession numbers of the species used are listed in Tables S2-5.

**Table S2.** Accession numbers of PCDH24 sequences used for alignment of EC repeats across different species.

| Abbreviation | Accession Number | Species | Common Name |
| --- | --- | --- | --- |
| *Hs* | NP_001165447.1 | *Homo sapiens* | Human |
| *Pt* | XP_016809791.1 | *Pan troglodytes* | Chimpanzee |
| *Rn* | XP_214434.4 | *Rattus norvegicus* | Norwegian rat |
| *Mm* | NP_001028536.2 | *Mus musculus* | Mouse |
| *Clf* | XP_005619238.1 | *Canis lupus familiaris* | Dog |
| *Fc* | XP_019693959.2 | *Felis catus* | Domestic cat |
| *Aj* | XP_014928203.2 | *Acinonyx jubatus* | Cheetah |
| *Bt* | NP_001179233.2 | *Bos taurus* | Cow |
| *Ss* | XP_013850328.2 | *Sus scrofa* | Pig |
| *Oa* | XP_004008751.3 | *Ovis aries* | Sheep |
| *Pc* | XP_020854166.1 | *Phascolarctos cinereus* | Koala |
| *Dl* | XP_022420063.1 | *Delphinapterus leucas* | Beluga whale |
| *Gg* | XP_015149366.1 | *Gallus gallus* | Chicken |
| *Ccc* | XP_019141545.2 | *Corvus cornix cornix* | Crow |
| *Af* | XP_019329391.1 | *Aptenodytes forsteri* | Emperor penguin |
| *Ac* | XP_008119531.1 | *Anolis carolinensis* | Green anole |
| *Dr* | XP_017214654.2 | *Danio rerio* | Zebrafish |
| *Am* | XP_022532075.1 | *Astyanax mexicanus* | Mexican tetra |
| *Om* | XP_021417551.1 | *Oncorhynchus mykiss* | Rainbow trout |

**Table S3.** Accession numbers of CDHR5 sequences used for alignment of EC repeats across different species.

| Abbreviation | Accession Number | Species | Common Name |
| --- | --- | --- | --- |
| *Hs* | NP_068743.2 | *Homo sapiens* | Human |
| *Pt* | XP_016775501.1 | *Pan troglodytes* | Chimpanzee |
| *Rn* | XP_006230571.1 | *Rattus norvegicus* | Norway rat |
| *Mm* | NP_001107794.1 | *Mus musculus* | Mouse |
| *Fc* | XP_019668438.1 | *Felis catus* | Domestic cat |
| *Bt* | NP_001096796.1 | *Bos taurus* | Cow |
| *Ss* | XP_013845324.2 | *Sus scrofa* | Pig |
| *Oa* | XP_027815739.1 | *Ovis aries* | Sheep |
| *Pc* | XP_020835078.1 | *Phascolarctos cinereus* | Koala |
| *Dl* | XP_022421100.1 | *Delphinapterus leucas* | Beluga whale |
| *Af* | XP_009281978.1 | *Aptenodytes forsteri* | Emperor penguin |
| *Ac* | XP_008106729.1 | *Anolis carolinensis* | Green anole |
| *Dr* | XP_021326278.1 | *Danio rerio* | Zebrafish |
| *Am* | XP_022525707.1 | *Astyanax mexicanus* | Mexican tetra |
| *Om* | XP_021454527.1 | *Oncorhynchus mykiss* | Rainbow trout |

**Table S4.** Accession numbers of PCDH15 sequences used for EC alignment and conservation analysis.

| Accession Number | Species | Common Name |
| --- | --- | --- |
| NP_001136235.1 | *Homo sapiens* | Human |
| NP_001136241.1 | *Homo sapiens* | Human |
| XP_019792025.1 | *Tursiops truncatus* | Bottlenose dolphin |
| XP_020929194.1 | *Sus scrofa* | pig |
| XP_022266127.1 | *Canis lupus familiaris* | Dog |
| NP_075604.2 | *Mus musculus* | mouse |
| XP_024416857.1 | *Desmodus rotundus* | Common vampire bat |
| XP_015143564.1 | *Gallus gallus* | Chicken |
| XP_021147002.1 | *Columbia livia* | Pigeon |
| XP_021391314.1 | *Lonchura striata domestica* | Finch |
| XP_017589248.1 | *Corvus brachyrhynchos* | American Crow |
| XP_020662251.1 | *Pogona vitticeps* | Bearded dragon |
| XP_016851436.1 | *Anolis carolinensis* | Green anole |
| XP_019394135.1 | *Crocodylus porosus* | Australian crocodile |
| NP_001012500.1 | *Danio rerio* | Zebrafish |
| XP_012675462.1 | *Clupea harengus* | Atlantic herring |
| XP_022525074.1 | *Astyanax mexicanus* | Mexican tetra |
| XP_007897895.1 | *Callorhinchus milii* | Australian ghostshark |

**Table S5.** Accession numbers of CDH23 sequences used for EC alignment and conservation analysis.

| Accession Number | Species | Common Name |
| --- | --- | --- |
| NP_071407.4 | *Homo sapiens* | Human |
| XP_016774005.2 | *Pan troglodytes* | Chimpanzee |
| XP_001925718.2 | *Sus scrofa* | Pig |
| NP_001178135.2 | *Bos taurus* | Cow |
| XP_014695776.1 | *Equus asinus* | Donkey |
| XP_023096304.1 | *Felis catus* | Cat |
| XP_022273252.1 | *Canis lupus familiaris* | Dog |
| XP_007454751.1 | *Lipotes vexillifer* | Yangtze river dolphin |
| NP_446096.1 | *Rattus norvegicus* | Norwegian rat |
| NP_075859.2 | *Mus musculus* | Mouse |
| XP_013028252.1 | *Anser cygnoides domesticus* | Domestic goose |
| XP_015143732.1 | *Gallus gallus* | Chicken |
| XP_015278377.1 | *Gekko japonicus* | Japanese gekko |
| XP_019339897.1 | *Alligator mississippiensis* | American alligator |
| XP_007897937.1 | *Callorhynchus milii* | Australian ghostshark |
| NP_999974.1 | *Danio rerio* | Zebrafish |
| XP_012672424.1 | *Clupea harengus* | Atlantic herring |

**Table S6.**
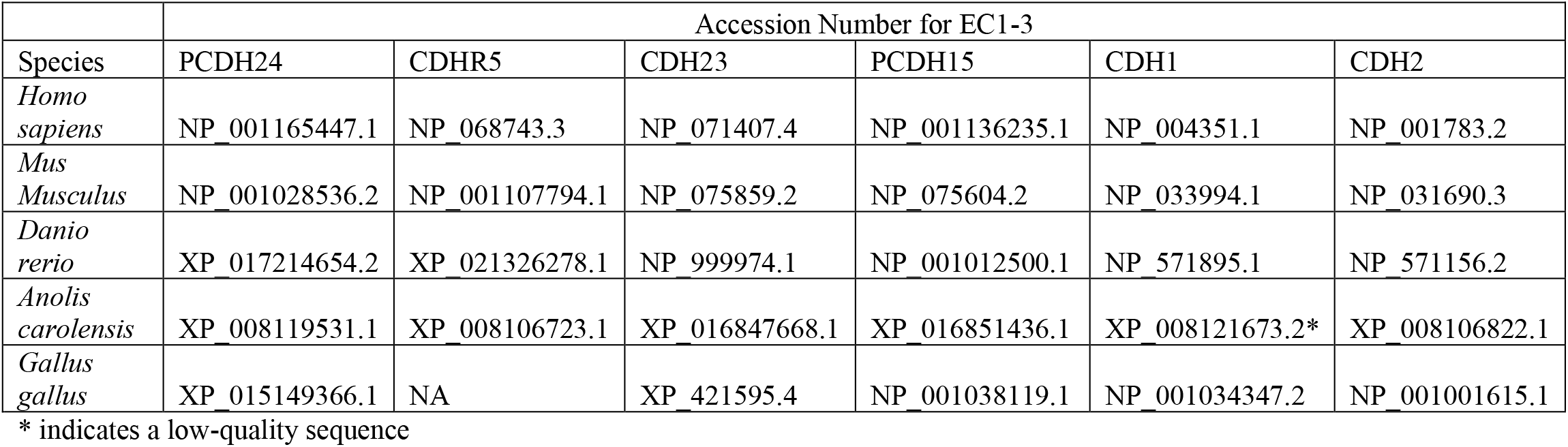
Accession numbers of sequences of PCDH24, CDHR5, CDH23, PCDH15, CDH1, and CDH2 used for generating Figure 1B.

**Table S7.**
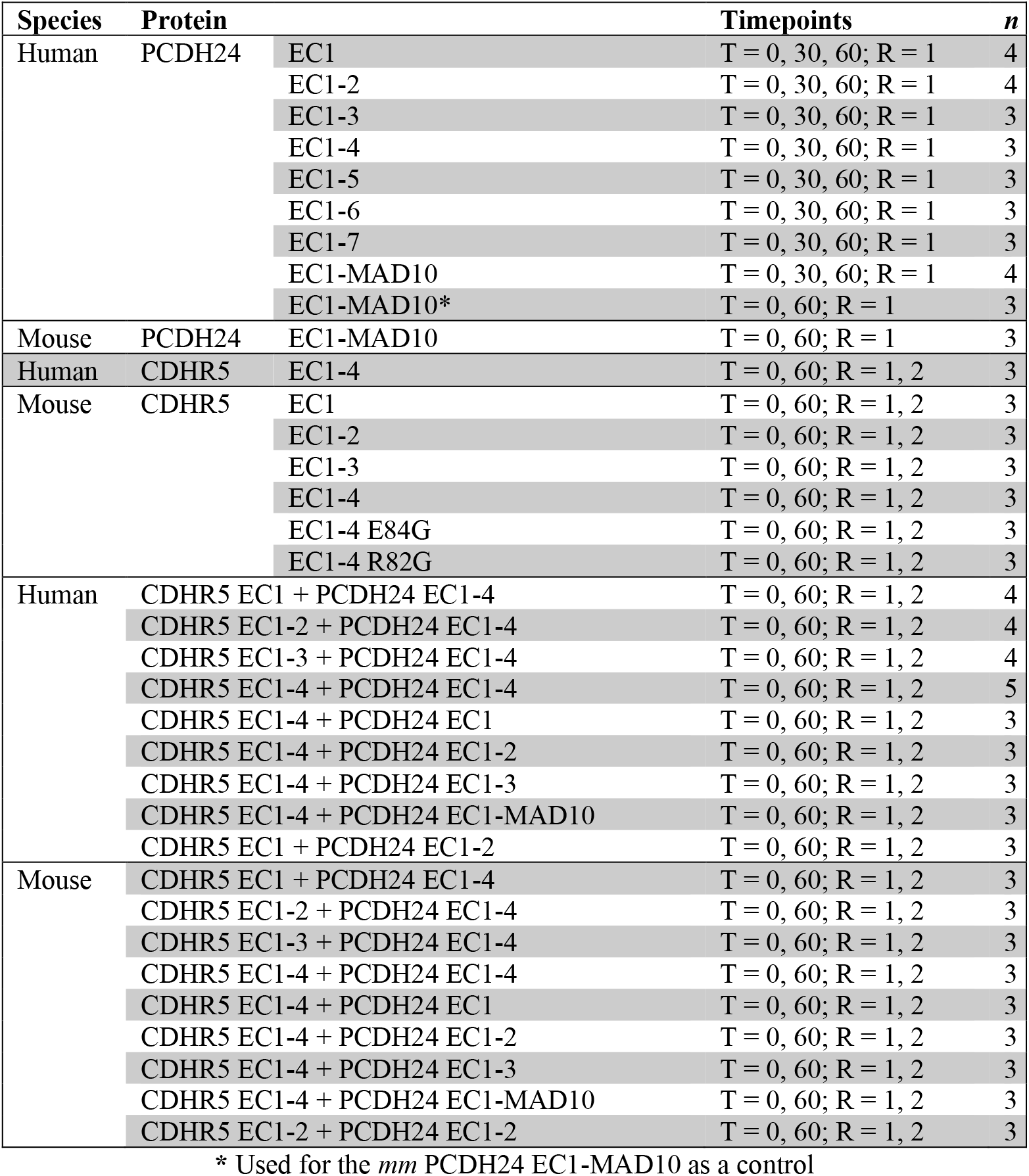
Number of biological replicates for bead aggregation assays.

| Species | Protein |  | Timepoints | *n* |
| --- | --- | --- | --- | --- |
| Human | PCDH24 | EC1 | T = 0, 30, 60; R = 1 | 4 |
|  |  | EC1-2 | T = 0, 30, 60; R = 1 | 4 |
|  |  | EC1-3 | T = 0, 30, 60; R = 1 | 3 |
|  |  | EC1-4 | T = 0, 30, 60; R = 1 | 3 |
|  |  | EC1-5 | T = 0, 30, 60; R = 1 | 3 |
|  |  | EC1-6 | T = 0, 30, 60; R = 1 | 3 |
|  |  | EC1-7 | T = 0, 30, 60; R = 1 | 3 |
|  |  | EC1-MAD10 | T = 0, 30, 60; R = 1 | 4 |
|  |  | EC1-MAD10* | T = 0, 60; R = 1 | 3 |
| Mouse | PCDH24 | EC1-MAD10 | T = 0, 60; R = 1 | 3 |
| Human | CDHR5 | EC1-4 | T = 0, 60; R = 1, 2 | 3 |
| Mouse | CDHR5 | EC1 | T = 0, 60; R = 1, 2 | 3 |
|  |  | EC1-2 | T = 0, 60; R = 1, 2 | 3 |
|  |  | EC1-3 | T = 0, 60; R = 1, 2 | 3 |
|  |  | EC1-4 | T = 0, 60; R = 1, 2 | 3 |
|  |  | EC1-4 E84G | T = 0, 60; R = 1, 2 | 3 |
|  |  | EC1-4 R82G | T = 0, 60; R = 1, 2 | 3 |
| Human | CDHR5 EC1 + PCDH24 EC1-4 | CDHR5 EC1 + PCDH24 EC1-4 | T = 0, 60; R = 1, 2 | 4 |
|  |  | CDHR5 EC1-2 + PCDH24 EC1-4 | T = 0, 60; R = 1, 2 | 4 |
|  |  | CDHR5 EC1-3 + PCDH24 EC1-4 | T = 0, 60; R = 1, 2 | 4 |
|  |  | CDHR5 EC1-4 + PCDH24 EC1-4 | T = 0, 60; R = 1, 2 | 5 |
|  |  | CDHR5 EC1-4 + PCDH24 EC1 | T = 0, 60; R = 1, 2 | 3 |
|  |  | CDHR5 EC1-4 + PCDH24 EC1-2 | T = 0, 60; R = 1, 2 | 3 |
|  |  | CDHR5 EC1-4 + PCDH24 EC1-3 | T = 0, 60; R = 1, 2 | 3 |
|  |  | CDHR5 EC1-4 + PCDH24 EC1-MAD10 | T = 0, 60; R = 1, 2 | 3 |
|  |  | CDHR5 EC1 + PCDH24 EC1-2 | T = 0, 60; R = 1, 2 | 3 |
| Mouse | CDHR5 EC1 + PCDH24 EC1-4 | CDHR5 EC1 + PCDH24 EC1-4 | T = 0, 60; R = 1, 2 | 3 |
|  |  | CDHR5 EC1-2 + PCDH24 EC1-4 | T = 0, 60; R = 1, 2 | 3 |
|  |  | CDHR5 EC1-3 + PCDH24 EC1-4 | T = 0, 60; R = 1, 2 | 3 |
|  |  | CDHR5 EC1-4 + PCDH24 EC1-4 | T = 0, 60; R = 1, 2 | 3 |
|  |  | CDHR5 EC1-4 + PCDH24 EC1 | T = 0, 60; R = 1, 2 | 3 |
|  |  | CDHR5 EC1-4 + PCDH24 EC1-2 | T = 0, 60; R = 1, 2 | 3 |
|  |  | CDHR5 EC1-4 + PCDH24 EC1-3 | T = 0, 60; R = 1, 2 | 3 |
|  |  | CDHR5 EC1-4 + PCDH24 EC1-MAD10 | T = 0, 60; R = 1, 2 | 3 |
|  |  | CDHR5 EC1-2 + PCDH24 EC1-2 | T = 0, 60; R = 1, 2 | 3 |

\* Used for the *mm* PCDH24 EC1-MAD10 as a control

**Table S8.**
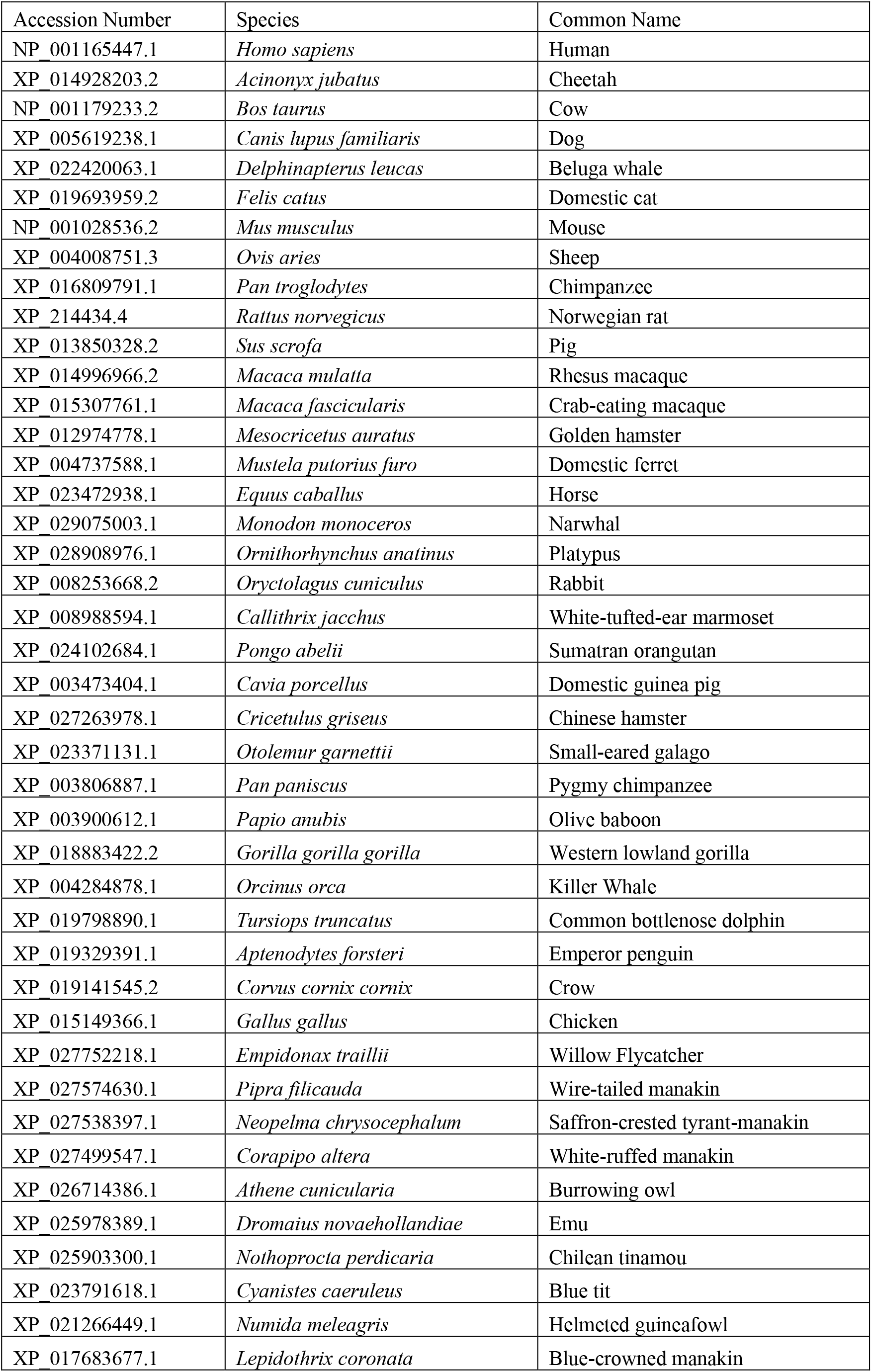
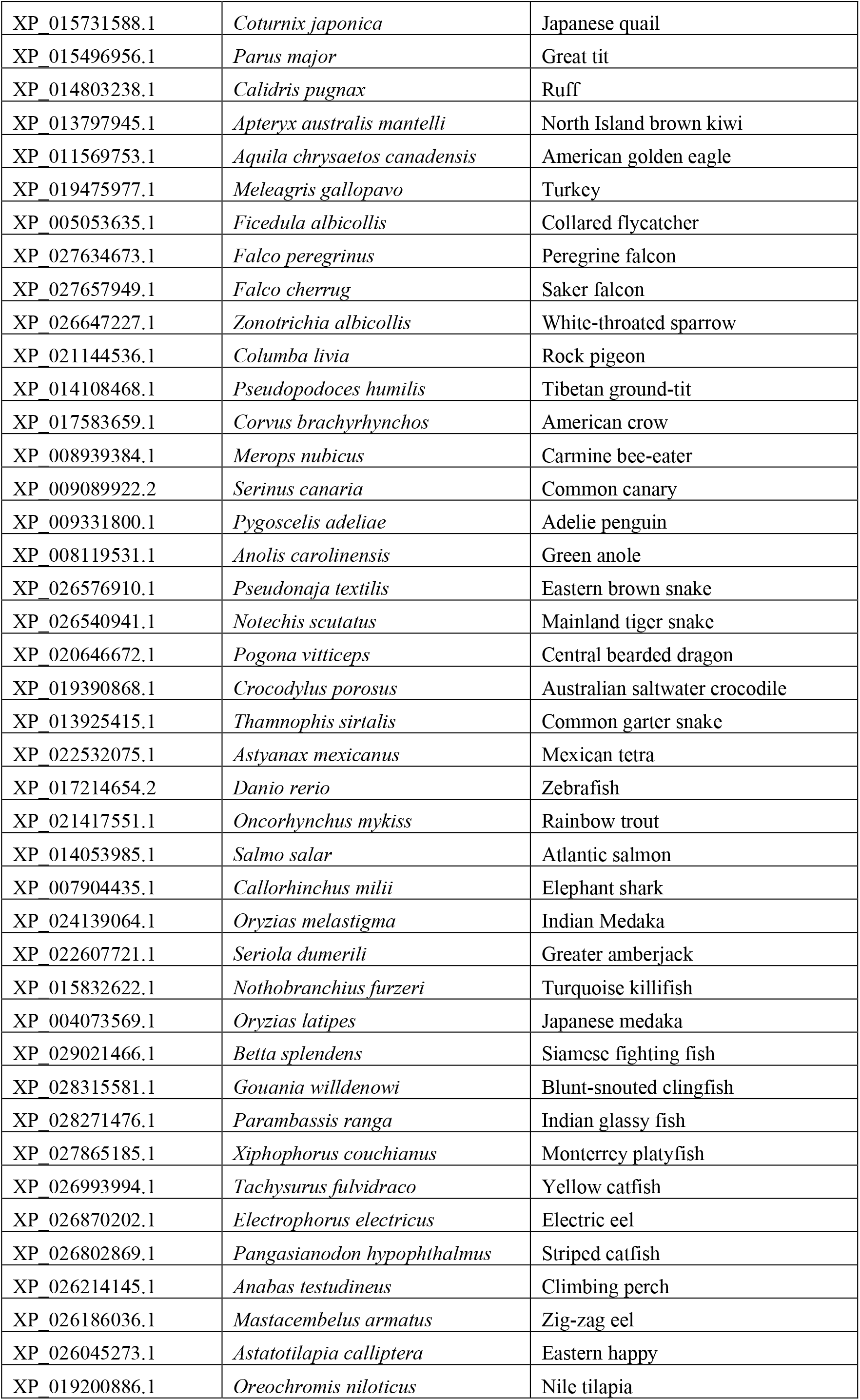
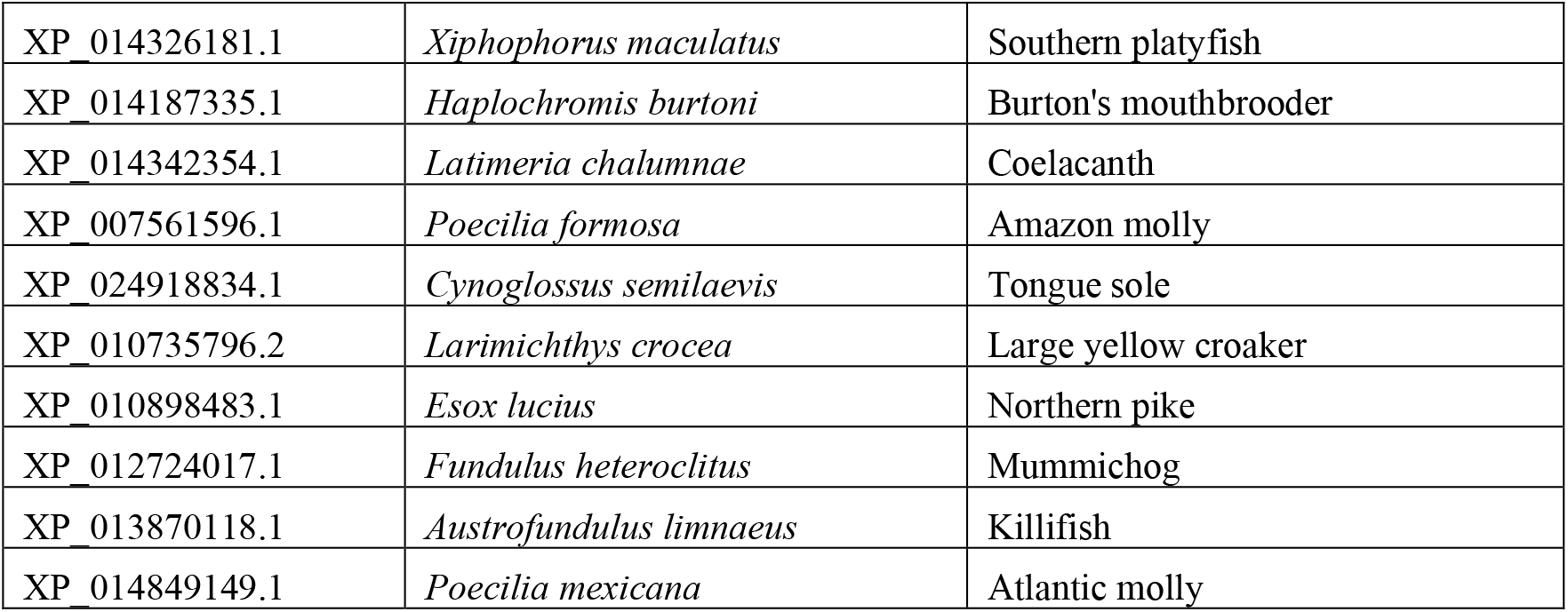
Accession numbers of sequences of PCDH24 used in Consurf.

| Accession Number | Species | Common Name |
| --- | --- | --- |
| NP_001165447.1 | *Homo sapiens* | Human |
| XP_014928203.2 | *Acinonyx jubatus* | Cheetah |
| NP_001179233.2 | *Bos taurus* | Cow |
| XP_005619238.1 | *Canis lupus familiaris* | Dog |
| XP_022420063.1 | *Delphinapterus leucas* | Beluga whale |
| XP_019693959.2 | *Felis catus* | Domestic cat |
| NP_001028536.2 | *Mus musculus* | Mouse |
| XP_004008751.3 | *Ovis aries* | Sheep |
| XP_016809791.1 | *Pan troglodytes* | Chimpanzee |
| XP_214434.4 | *Rattus norvegicus* | Norwegian rat |
| XP_013850328.2 | *Sus scrofa* | Pig |
| XP_014996966.2 | *Macaca mulatta* | Rhesus macaque |
| XP_015307761.1 | *Macaca fascicularis* | Crab-eating macaque |
| XP_012974778.1 | *Mesocricetus auratus* | Golden hamster |
| XP_004737588.1 | *Mustela putorius furo* | Domestic ferret |
| XP_023472938.1 | *Equus caballus* | Horse |
| XP_029075003.1 | *Monodon monoceros* | Narwhal |
| XP_028908976.1 | *Ornithorhynchus anatinus* | Platypus |
| XP_008253668.2 | *Oryctolagus cuniculus* | Rabbit |
| XP_008988594.1 | *Callithrix jacchus* | White-tufted-ear marmoset |
| XP_024102684.1 | *Pongo abelii* | Sumatran orangutan |
| XP_003473404.1 | *Cavia porcellus* | Domestic guinea pig |
| XP_027263978.1 | *Cricetulus griseus* | Chinese hamster |
| XP_023371131.1 | *Otolemur garnettii* | Small-eared galago |
| XP_003806887.1 | *Pan paniscus* | Pygmy chimpanzee |
| XP_003900612.1 | *Papio anubis* | Olive baboon |
| XP_018883422.2 | *Gorilla gorilla gorilla* | Western lowland gorilla |
| XP_004284878.1 | *Orcinus orca* | Killer Whale |
| XP_019798890.1 | *Tursiops truncatus* | Common bottlenose dolphin |
| XP_019329391.1 | *Aptenodytes forsteri* | Emperor penguin |
| XP_019141545.2 | *Corvus cornix cornix* | Crow |
| XP_015149366.1 | *Gallus gallus* | Chicken |
| XP_027752218.1 | *Empidonax traillii* | Willow Flycatcher |
| XP_027574630.1 | *Pipra filicauda* | Wire-tailed manakin |
| XP_027538397.1 | *Neopelma chrysocephalum* | Saffron-crested tyrant-manakin |
| XP_027499547.1 | *Corapipo altera* | White-ruffed manakin |
| XP_026714386.1 | *Athene cunicularia* | Burrowing owl |
| XP_025978389.1 | *Dromaius novaehollandiae* | Emu |
| XP_025903300.1 | *Nothoprocta perdicaria* | Chilean tinamou |
| XP_023791618.1 | *Cyanistes caeruleus* | Blue tit |
| XP_021266449.1 | *Numida meleagris* | Helmeted guineafowl |
| XP_017683677.1 | *Lepidothrix coronata* | Blue-crowned manakin |
| XP_015731588.1 | *Coturnix japonica* | Japanese quail |
| XP_015496956.1 | *Parus major* | Great tit |
| XP_014803238.1 | *Calidris pugnax* | Ruff |
| XP_013797945.1 | *Apteryx australis mantelli* | North Island brown kiwi |
| XP_011569753.1 | *Aquila chrysaetos canadensis* | American golden eagle |
| XP_019475977.1 | *Meleagris gallopavo* | Turkey |
| XP_005053635.1 | *Ficedula albicollis* | Collared flycatcher |
| XP_027634673.1 | *Falco peregrinus* | Peregrine falcon |
| XP_027657949.1 | *Falco cherrug* | Saker falcon |
| XP_026647227.1 | *Zonotrichia albicollis* | White-throated sparrow |
| XP_021144536.1 | *Columba livia* | Rock pigeon |
| XP_014108468.1 | *Pseudopodoces humilis* | Tibetan ground-tit |
| XP_017583659.1 | *Corvus brachyrhynchos* | American crow |
| XP_008939384.1 | *Merops nubicus* | Carmine bee-eater |
| XP_009089922.2 | *Serinus canaria* | Common canary |
| XP_009331800.1 | *Pygoscelis adeliae* | Adelie penguin |
| XP_008119531.1 | *Anolis carolinensis* | Green anole |
| XP_026576910.1 | *Pseudonaja textilis* | Eastern brown snake |
| XP_026540941.1 | *Notechis scutatus* | Mainland tiger snake |
| XP_020646672.1 | *Pogona vitticeps* | Central bearded dragon |
| XP_019390868.1 | *Crocodylus porosus* | Australian saltwater crocodile |
| XP_013925415.1 | *Thamnophis sirtalis* | Common garter snake |
| XP_022532075.1 | *Astyanax mexicanus* | Mexican tetra |
| XP_017214654.2 | *Danio rerio* | Zebrafish |
| XP_021417551.1 | *Oncorhynchus mykiss* | Rainbow trout |
| XP_014053985.1 | *Salmo salar* | Atlantic salmon |
| XP_007904435.1 | *Callorhynchus milii* | Elephant shark |
| XP_024139064.1 | *Oryzias melastigma* | Indian Medaka |
| XP_022607721.1 | *Seriola dumerili* | Greater amberjack |
| XP_015832622.1 | *Nothobranchius furzeri* | Turquoise killifish |
| XP_004073569.1 | *Oryzias latipes* | Japanese medaka |
| XP_029021466.1 | *Betta splendens* | Siamese fighting fish |
| XP_028315581.1 | *Gouania willdenowi* | Blunt-snouted clingfish |
| XP_028271476.1 | *Parambassis ranga* | Indian glassy fish |
| XP_027865185.1 | *Xiphophorus couchianus* | Monterrey platyfish |
| XP_026993994.1 | *Tachysurus fulvidraco* | Yellow catfish |
| XP_026870202.1 | *Electrophorus electricus* | Electric eel |
| XP_026802869.1 | *Pangasianodon hypophthalmus* | Striped catfish |
| XP_026214145.1 | *Anabas testudineus* | Climbing perch |
| XP_026186036.1 | *Mastacembelus armatus* | Zig-zag eel |
| XP_026045273.1 | *Astatotilapia calliptera* | Eastern happy |
| XP_019200886.1 | *Oreochromis niloticus* | Nile tilapia |
| XP_014326181.1 | *Xiphophorus maculatus* | Southern platyfish |
| XP_014187335.1 | *Haplochromis burtoni* | Burton's mouthbrooder |
| XP_014342354.1 | *Latimeria chalumnae* | Coelacanth |
| XP_007561596.1 | *Poecilia formosa* | Amazon molly |
| XP_024918834.1 | *Cynoglossus semilaevis* | Tongue sole |
| XP_010735796.2 | *Larimichthys crocea* | Large yellow croaker |
| XP_010898483.1 | *Esox lucius* | Northern pike |
| XP_012724017.1 | *Fundulus heteroclitus* | Mummichog |
| XP_013870118.1 | *Austrofundulus limnaeus* | Killifish |
| XP_014849149.1 | *Poecilia mexicana* | Atlantic molly |

**Table S9.** Accession numbers of sequences of CDHR5 used in Consurf.

| Accession Number | Species | Common Name |
| --- | --- | --- |
| NP_068743.2 | *Homo sapiens* | Human |
| NP_001107794.1 | *Mus musculus* | Mouse |
| XP_027815739.1 | *Ovis aries* | Sheep |
| XP_022421100.1 | *Delphinapterus leucas* | Beluga whale |
| XP_020835078.1 | *Phascolarctos cinereus* | Koala |
| XP_016775501.1 | *Pan troglodytes* | Chimpanzee |
| XP_006230571.1 | *Rattus norvegicus* | Norway rat |
| XP_013845324.2 | *Sus scrofa* | Pig |
| NP_001096796.1 | *Bos taurus* | Cow |
| XP_019668438.1 | *Felis catus* | Domestic cat |
| XP_023510521.1 | *Equus caballus* | Horse |
| XP_014437989.1 | *Tupaia chinensis* | Chinese tree shrew |
| XP_012973076.1 | *Mesocricetus auratus* | Golden hamster |
| XP_029064308.1 | *Monodon monoceros* | Narwhal |
| XP_028642741.1 | *Grammomys surdaster* | African woodland thicket rat |
| XP_028370928.1 | *Phyllostomus discolor* | Pale spear-nosed bat |
| XP_025847484.1 | *Vulpes vulpes* | Red fox |
| XP_025717837.1 | *Callorhinus ursinus* | Northern ful seal |
| XP_024430321.1 | *Desmodus rotundus* | Common vampire bat |
| XP_020767741.1 | *Odocoileus virginianus texanus* | White-tailed deer |
| XP_009281978.1 | *Aptenodytes forsteri* | Emperor penguin |
| XP_025966497.1 | *Dromaius novaehollandiae* | Emu |
| XP_025928961.1 | *Apteryx rowi* | Okarito brown kiwi |
| XP_025900649.1 | *Nothoprocta perdicaria* | Chilean tinamou |
| XP_023784276.1 | *Cyanistes caeruleus* | Blue tit |
| XP_015485075.1 | *Parus major* | Great tit |
| XP_015718593.1 | *Coturnix japonica* | Japanese quail |
| XP_014796130.1 | *Calidris pugnax* | Ruff |
| XP_014734396.1 | *Sturnus vulgaris* | Common starling |
| XP_013800822.1 | *Apteryx australis mantelli* | North Island brown kiwi |
| XP_013053829.1 | *Anser cygnoides domesticus* | Domestic goose |
| XP_011595784.1 | *Aquila chrysaetos canadensis* | American golden eagle |
| XP_010709142.1 | *Meleagris gallopavo* | Turkey |
| XP_010562687.1 | *Haliaeetus leucocephalus* | Bald eagle |
| XP_010297449.1 | *Balearica regulorum gibbericeps* | East African grey crowned-crane |
| XP_010292632.1 | *Phaethon lepturus* | White-tailed tropicbird |
| XP_010225820.1 | *Tinamus guttatus* | White-throated tinamou |
| XP_010207507.1 | *Colius striatus* | Speckled mousebird |
| XP_007907548.1 | *Callorhynchus milii* | Elephant shark |
| XP_021454527.1 | *Oncorhynchus mykiss* | Rainbow trout |
| XP_021326278.1 | *Danio rerio* | Zebrafish |
| XP_022525707.1 | *Astyanax mexicanus* | Mexican tetra |
| XP_024120964.1 | *Oryzias melastigma* | Indian medaka |
| XP_023807528.1 | *Oryzias latipes* | Japanese medaka |
| XP_029312546.1 | *Cottoperca gobio* | Channel bull blenny |
| XP_027008843.1 | *Tachysurus fulvidraco* | Yellow catfish |
| XP_026168289.1 | *Mastacembelus armatus* | Zig-zag eel |
| XP_019112627.2 | *Larimichthys crocea* | Large yello croaker |
| XP_017341402.1 | *Ictalurus punctatus* | Channel catfish |
| XP_016392038.1 | *Sinocyclocheilus rhinoceros* | Rayfinned fish |
| XP_016324417.1 | *Sinocyclocheilus anshuiensis* | Rayfinned fish |
| XP_015227070.1 | *Cyprinodon variegatus* | Sheepshead minnow |
| XP_021163353.1 | *Fundulus heteroclitus* | Mummichog |
| XP_012674652.1 | *Clupea harengus* | Atlantic herring |
| XP_005938389.1 | *Haplochromis burtoni* | Burton's mouthbrooder |
| XP_004556350.2 | *Maylandia zebra* | Zebra mbuna |
| XP_008106729.1 | *Anolis carolinensis* | Green anole |
| XP_019340716.1 | *Alligator mississippiensis* | American alligator |
| XP_028591121.1 | *Podarcis muralis* | Common wall lizard |
| XP_026562223.1 | *Pseudonaja textilis* | Eastern brown snake |
| XP_026523924.1 | *Notechis scutatus* | Mainland tiger snake |
| XP_026513403.1 | *Terrapene carolina triunguis* | Three-toed box turtle |
| XP_019402850.1 | *Crocodylus porosus* | Australian saltwater crocodile |
| XP_019375347.1 | *Gavialis gangeticus* | Gharial |
| XP_015745583.1 | *Python bivittatus* | Burmese python |
| XP_013925033.1 | *Thamnophis sirtalis* | Common garter snake |
| XP_014429985.1 | *Pelodiscus sinensis* | Chinese softshell turtle |
| XP_025054327.1 | *Alligator sinensis* | Chinese alligator |
| XP_023967188.1 | *Chrysemys picta bellii* | Western painted turtle |

**Table S10.**
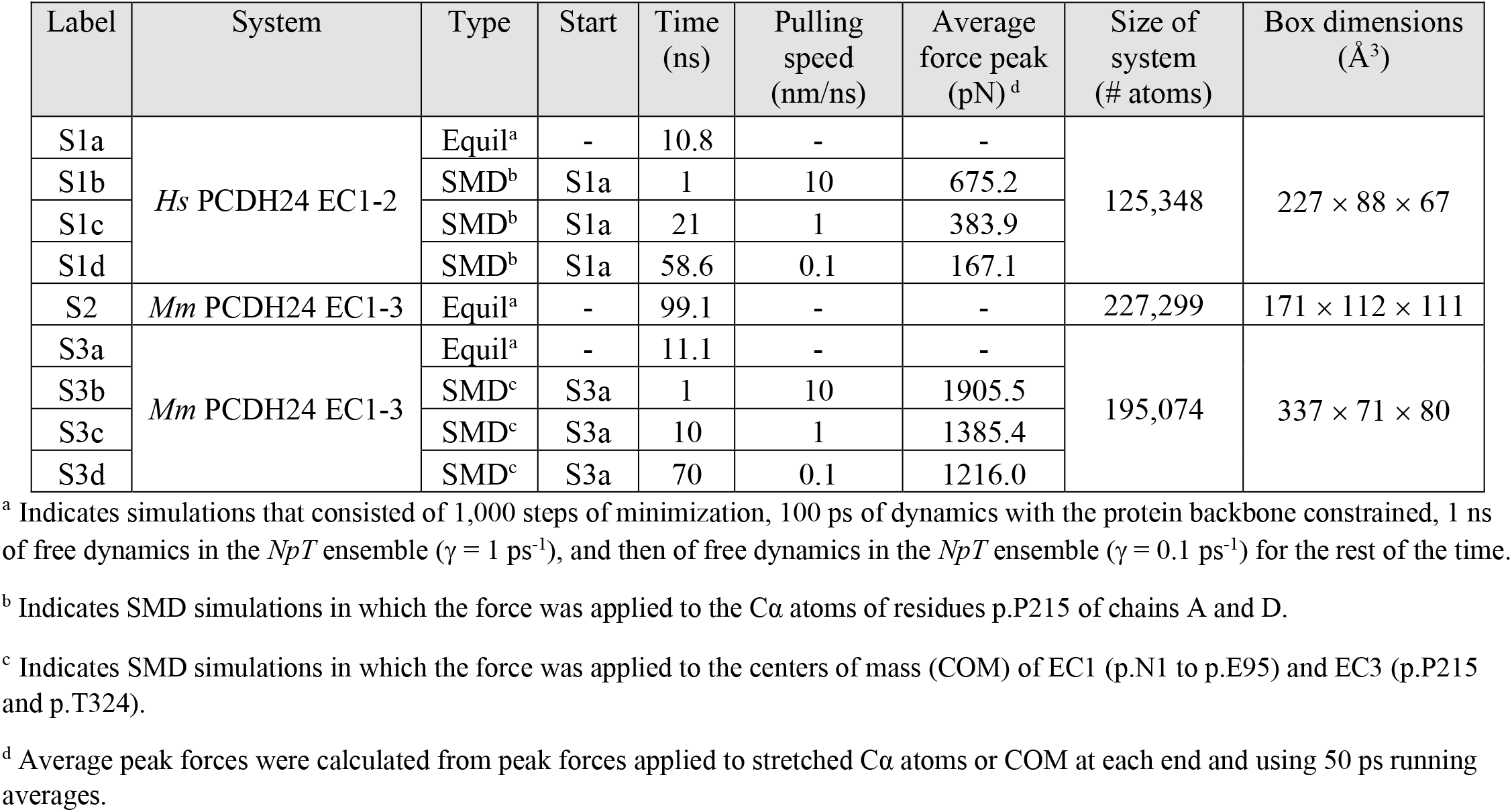
Summary of simulations

| Label | System | Type | Start | Time (ns) | Pulling speed (nm/ns) | Average force peak (pN)<sup>d</sup> | Size of system (# atoms) | Box dimensions (Å<sup>3</sup>) |
| --- | --- | --- | --- | --- | --- | --- | --- | --- |
| S1a | *Hs* PCDH24 EC1-2 | Equil<sup>a</sup> | - | 10.8 | - | - | 125,348 | 227 × 88 × 67 |
| S1b |  | SMD<sup>b</sup> | S1a | 1 | 10 | 675.2 |  |  |
| S1c |  | SMD<sup>b</sup> | S1a | 21 | 1 | 383.9 |  |  |
| S1d |  | SMD<sup>b</sup> | S1a | 58.6 | 0.1 | 167.1 |  |  |
| S2 | *Mm* PCDH24 EC1-3 | Equil<sup>a</sup> | - | 99.1 | - | - | 227,299 | 171 × 112 × 111 |
| S3a | *Mm* PCDH24 EC1-3 | Equil<sup>a</sup> | - | 11.1 | - | - | 195,074 | 337 × 71 × 80 |
| S3b |  | SMD<sup>c</sup> | S3a | 1 | 10 | 1905.5 |  |  |
| S3c |  | SMD<sup>c</sup> | S3a | 10 | 1 | 1385.4 |  |  |
| S3d |  | SMD<sup>c</sup> | S3a | 70 | 0.1 | 1216.0 |  |  |

<sup>a</sup> Indicates simulations that consisted of 1,000 steps of minimization, 100 ps of dynamics with the protein backbone constrained, 1 ns of free dynamics in the *NpT* ensemble ( $\gamma = 1 \text{ ps}^{-1}$ ), and then of free dynamics in the *NpT* ensemble ( $\gamma = 0.1 \text{ ps}^{-1}$ ) for the rest of the time.
<sup>b</sup> Indicates SMD simulations in which the force was applied to the Cα atoms of residues p.P215 of chains A and D.
<sup>c</sup> Indicates SMD simulations in which the force was applied to the centers of mass (COM) of EC1 (p.N1 to p.E95) and EC3 (p.P215 and p.T324).
<sup>d</sup> Average peak forces were calculated from peak forces applied to stretched Cα atoms or COM at each end and using 50 ps running averages.

## Notes

### Competing Interest Statement

The authors have declared no competing interest.

